# Capture of Post-Pluripotency Human Neuromesodermal and Neural Tube Axial Stem Cell States

**DOI:** 10.1101/2024.03.26.586760

**Authors:** Dolunay Kelle, Enes Ugur, Ejona Rusha, Dmitry Shaposhnikov, Alessandra Livigni, Sandra Horschitz, Mahnaz Davoudi, Atilgan Yilmaz, Kilian Winiger, Anahi Binagui-Casas, Angela Moreno Anguita, Xiaolong Yang, Mazène Hochane, Andreas Blutke, Judith Bushe, Michael Sterr, Ksenia Arkhipova, Benjamin Tak, Ruben de Vries, Alena Boos, Yuting Wang, Nina Schulten, Britte Spruijt, Raffee Wright, Ankit Sinha, Aicha Haji Ali, Christian Schröter, Heiko Lickert, Annette Feuchtinger, Zhenyu Xiao, Hongmei Wang, Pascal Giehr, Philipp Koch, Matthias Mann, Heinrich Leonhardt, Valerie Wilson, Micha Drukker

**Affiliations:** Institute of Stem Cell Research, Helmholtz Zentrum München, German Research Center for Environmental Health, Neuherberg, 85764, Germany; Induced Pluripotent Stem Cell Core Facility, Helmholtz Zentrum München, German Research Center for Environmental Health, Neuherberg, 85764, Germany; Faculty of Biology and Center for Molecular Biosystems (BioSysM), Human Biology and BioImaging, Ludwig-Maximilians-Universität München, Munich, 81377, Germany; Department of Proteomics and Signal Transduction, Max-Planck Institute of Biochemistry, Martinsried, 82152, Germany; Institute for Regeneration and Repair, Centre for Regenerative Medicine, Institute for Stem Cell Research, School of Biological Sciences, University of Edinburgh, 5 Little France Drive, Edinburgh EH16 4UU, Scotland; Central Institute of Mental Health, Heidelberg University/ Medical Faculty Mannheim, Mannheim, Germany; Hector Institute for Translational Brain Research (HITBR gGmbH), Mannheim, Germany; German Cancer Research Center, Heidelberg, Germany; Leuven Stem Cell Institute, Department of Development and Regeneration, KU Leuven, 3000 Leuven, Belgium; Department of Chemistry, Institute for Chemical Epigenetics, Ludwig-Maximilians-Universität München, Butenandtstr. 5-13, 81377 München, Germany; School of Life Science, Beijing Institute of Technology, Beijing 100081, China; Division of Cell Systems and Drug Safety, Leiden Academic Centre for Drug Research (LACDR), Leiden University, 2333 CC Leiden, The Netherlands; Core Facility Pathology & Tissue Analytics, Helmholtz Munich, 85764 Neuherberg, Germany; Institute of Diabetes and Regeneration Research, Helmholtz Center, Munich, Germany; German Center for Diabetes Research, Neuherberg, Germany; Core Facility Bioinformatics and Statistics, Helmholtz Zentrum München GmbH - German Research Center for Environmental Health, 85764 Neuherberg, Germany; School of Medicine, Technical University of Munich, Munich, Germany; Beijing Institute for Stem Cell and Regenerative Medicine, Beijing, China; Department of Internal Medicine, Leiden University Medical Center, Albinusdreef 2, 2333 ZA Leiden, The Netherlands

## Abstract

The capture of pluripotent and adult tissue stem cells has transformed biomedicine, yet stable stem cell states between these developmental extremes have remained elusive. Here, we capture two human axial stem cell (AxSC) states corresponding to neuromesodermal and derivative neural tube progenitors. These states maintain telomere length without teratoma formation, and neuromesodermal AxSCs were recaptured from human embryonic tail buds. Mechanistically, genome-wide knockout screening and multiomics analyses revealed capture-associated acquisition of a stemness regulatory network with Trithorax-Polycomb dual functionality. Integration with genome-wide DNA methylation analysis identified a novel HOX timer driven by time-grouped DNA demethylation of target genes. Additionally, stage-specific glycolytic/OXPHOS remodeling in AxSCs was linked to bioenergetic pathways and histone acetylation dynamics. Finally, we captured mouse AxSCs that contribute to chimeras and exhibit an evolutionary divergent neuromesodermal-neural combined state. AxSCs are authentic post pluripotent stem cells for upscalable interrogation of development, with potential for regenerative medicine and drug discovery.

**Highlights:**

- Capture of human AxSCs reveal two post-pluripotent stem cell states
- AxSCs represent neuromesodermal and neural tube progenitors with activated self-renewal
- Long-term self-renewal pauses developmental progression while sustaining axial identity
- CRISPR screening reveals capture-associated acquisition of a stemness network
- Timed and grouped demethylation of HOX targets links epigenetics to axial progression
- OXPHOS and glycolysis ensure axial stage specific bioenergetics and epigenetic remodeling
- Mouse AxSCs reveal evolutionary divergence in mammalian axial progenitors

## INTRODUCTION

Embryonic development proceeds through successive transient progenitor states that, if they self-renew, do so only briefly during differentiation. The derivation of induced and embryonic pluripotent stem cells (i/ePSCs) demonstrated the capture in permanent self-renewal of early embryonic states. It revealed that ground, rosette, formative, and primed PSC states are maintained by distinct signaling, epigenetic and metabolic mechanisms.^1–7^ However, mechanisms for capturing transient post-pluripotent states remain largely unexplored.

Multipotent axial progenitors that form the elongating amniote body axis are attractive candidates for uncovering such mechanisms. These transient progenitors self-renew while producing the posterior neural tube (NT) and somitic mesoderm. These tissues then further differentiate into the spinal cord, peripheral motor and sensory structures, and musculoskeletal tissues, extending from the cervical region to the tail. ^8–10^ However, whether the progenitors of these tissues can themselves be captured as stable long-term self-renewing stem cells remains unresolved.

In the mouse, axial elongation is driven by neuromesodermal progenitors (NMPs) that are located at the posterior end of the embryo: the node-streak border (NSB) and the caudal lateral epiblast (CLE) at early stages, and the chordo-neural hinge (CNH) of the tail bud later in axis elongation. ^11,12^ Mouse NMPs are characterized by co-expression of *Sox2, T (Bra)* and *Cdx2.*^13–15^ In addition to bipotent SOX2⁺/T (BRA)⁺ NMPs, chick embryos appear to harbor a distinct proliferative neural progenitor population expressing *Sox2* at the caudal tip of the NT.^16,17^ Whether axial progenitors in humans and primates resemble the NMP population in the mouse, or comprise distinct NT progenitors and bipotent NMPs remains unclear.

During axial elongation, NMPs and their progeny activate *Hox* genes. These are organized within four clusters into 13 paralogous groups (PG), whose nested anteroposterior expression establishes positional identity. Early mouse NMPs express neck and upper trunk HOX PG5–8, while later ones express lower trunk and tail PG9–13.^12,14^ Consistently, a human embryo at post conception week (PCW) 3 shows *HOXA2* expression in the hindbrain, whereas more posterior HOX paralog groups are expressed in the NT.^9^ Mechanisms underlying unfolding of axial HOX programs through tail-level identities include chromatin remodeling.^18^ In particular, progressive removal of H3K27me3 permits activation of increasingly posterior HOX programs during axial elongation.^19^ However, transient post-pluripotent human cell states are not readily expandable, limiting the mechanistic biochemical interrogation of HOX regulation.

Formation of mouse NMPs relies on WNT and FGF signaling, which promote renewal of posterior progenitors and regulate *Hox* expression through *Cdx* genes.^15,20–23^ This signaling framework has been leveraged in NMP differentiation protocols of human and mouse i/ePSCs. Protocols typically employ 2–3 day activation of Wnt/β-catenin and FGF2 in a monolayer to generate transient NMP-like cells expressing *SOX2*, *T*, and *CDX2*.^22,24–27^ Transient NMP-like cells generated with this strategy have been used as an intermediate to differentiate regionally patterned NT progenitors expressing *SOX2* and *PAX6* with integration potential in chick embryo.^28–33^ While mouse NMPs can be passaged through serial embryos suggesting self-renewal,^10^ *in vitro* human and murine NMP- and NT-like progenitors appear transient.^24,25,27^ In accordance, human progenitor cultures with the capacity to differentiate into neuronal and skeletal muscle lineages have been maintained through passaging.^34^ However, these cultures display a mixture of T (BRA)^+^ and PAX6^+^ cells, require EGF that is not aligned with NMP/NT signaling, and remain unverified in human and animal embryos. Thus, stable self-renewing AxSC lines remain to be established and validated in both human and murine embryos.

During PSC differentiation, chromatin resolves from a predominantly open euchromatic state, in which lineage-specific genes are maintained in configurations marked by both H3K4me3 and H3K27me3, into progressively more defined locally active and repressed states.^35,36^ Changes in this bivalency and chromatin accessibility are paralleled by metabolic reprogramming from glycolysis toward mitochondrial respiration.^37^ It has been shown that redirecting acetyl-CoA toward mitochondrial metabolism restricts histone acetylation during differentiation, thereby linking epigenetic and metabolic programs.^38–41^ Capturing stable AxSCs therefore requires recapitulating these epigenetic and metabolic transitions, creating a tractable system to uncover the biochemical mechanisms governing stemness acquisition, developmental timing, and axial identity.

Here, we identify signaling regimes that support AxSC derivation from PSCs and embryos across genetic backgrounds and species. These cocktails combine the Wnt/β-catenin activator CHIR99021, the TGF-β inhibitor SB431542, with or without FGF2 (CFS and CS), and support the derivation and long-term self-renewal of two distinct AxSC states. Through functional differentiation and mapping to embryonic transcriptomes, we demonstrate that permanent human CFS and CS lines capture states that reflect embryonic NMPs and NT progenitors, respectively. Consistent with the differentiation trajectory of progenitors *in vivo* from NMP to NT progenitors, CFS cells readily undergo differentiation to a CS phenotype but not vice versa. Mouse posterior progenitors could only be maintained under CFS conditions and converged on a hybrid state between NMPs and NT progenitors. This finding hints at fundamental primate-rodent differences in axial formation.

To characterise the molecular integration underlying capture and self-renewal of AxSCs, we performed integrated multiomic analyses. We combined scRNA-seq, genome-wide CRISPR/Cas9 screening, proteomics of whole cells, chromatin fractions as well as histone post-translational modifications (PTMs), DNA methylation analysis, and functional metabolic and epigenetic perturbations. These analyses uncover stemness acquisition pathways, molecular timer linked to stage-specific epigenetic-metabolic reconfigurations, and telomere maintenance in post pluripotency stem cell states. More broadly, these findings reveal post-pluripotent developmental stages in which specific chromatin and metabolic configurations become permissive for long-term self-renewal, suggesting the existence of a broader landscape of such developmental windows. AxSC multiomic datasets are accessible through an interactive web resource at https://axialstemcells.shinyapps.io/atlas/.

## RESULTS

### Capture of human long-term self-renewing multipotent axial stem cell states

To derive long-term post-pluripotent human stem cells, we first sought to transition ePSCs to an optimal capture window between pluripotency and early organogenesis programs. Time-course RNA-seq upon Wnt/β-catenin activation using both CHIR99021 and functionally constitutive polyclonal ΔN90 β-catenin revealed strong upregulation of *T (BRA)* and *CDX2* and maintenance of *SOX2* expression at 24h (Figure S1A–C). This was followed by induction of more advanced developmental markers, including *GATA6* and *FGF17*, and downregulation of *SOX2* at 48-72h. We therefore selected the 24h CHIR99021 condition as a post-pluripotent posterior gastrulation-like state for subsequent self-renewal screening (Figure 1A).

**Figure 1.**
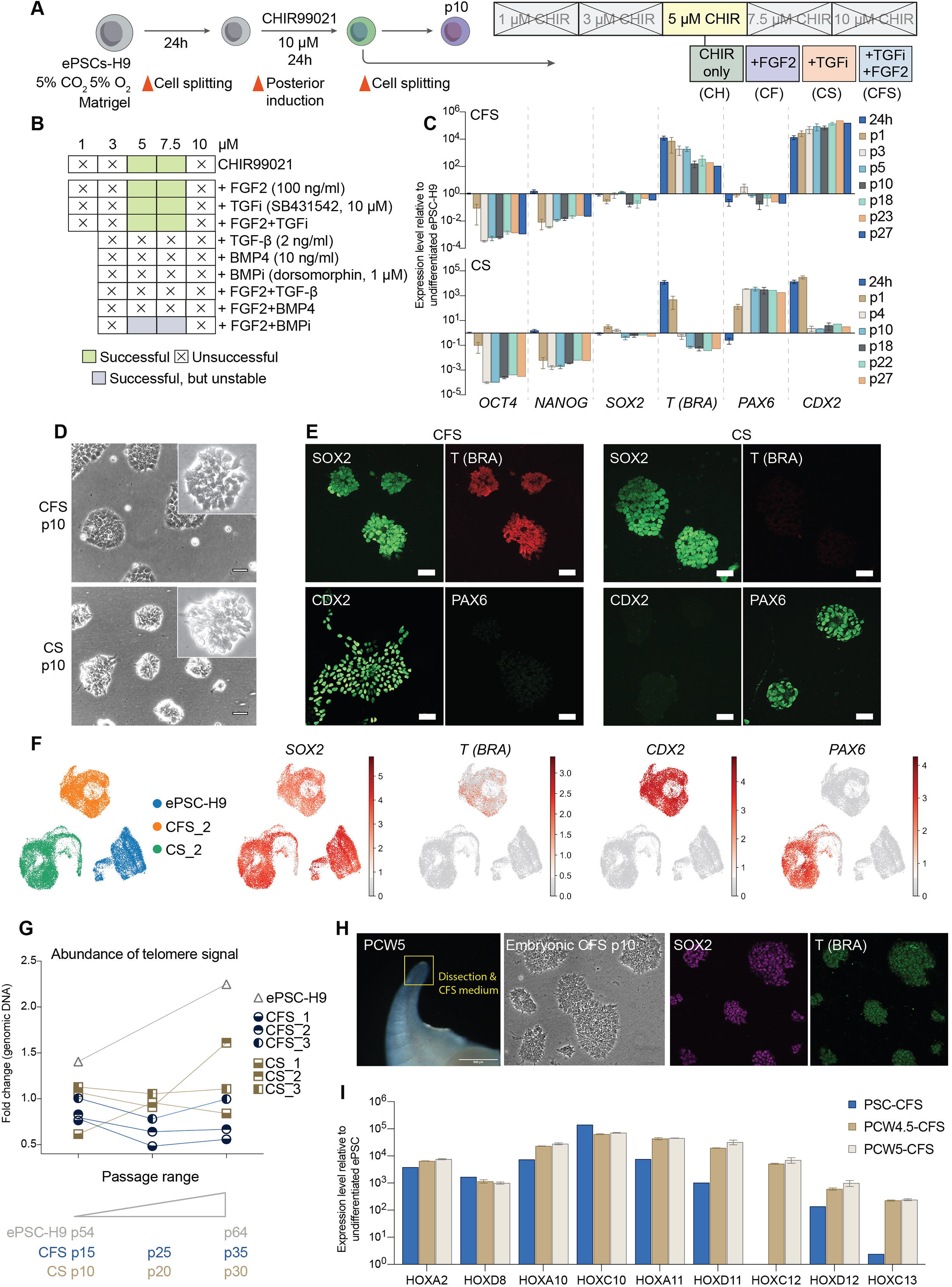
Capture of axial stem cell lines from human PSCs and embryos. **A**, The screening strategy for identifying conditions promoting self-renewal of posterior progenitors from human ES cells following 24h WNT/β-catenin induction by CHIR99021 and cell splitting in conjunction with cultivation with cocktails of small molecule and growth factors (as indicated) for 10 passages. Cultures were dissociated to single cells when passaged on Matrigel-coated plates in 5% CO_2_ / 5% O_2_, and B27 medium without vitamin A or antibiotics. **B**, The green rectangles indicate conditions that enabled successful establishment of cell lines for >10 passages, and gray rectangles represent cell lines that were discontinued after 9 passages. Crossed-out rectangles represent screened conditions that did not support the establishment of renewing cell lines. CFS and CS denote cell lines that were established in CHIR99021, FGF2, and TGF-β pathway inhibitor SB431542, or without FGF2, respectively, with passaging every 3-4 and 5-7 days respectively. The analysis of CH and CF lines is shown in Figure S1F. **C**, RT-qPCR characterization of H9 CFS and CS lines using pluripotency markers, NMP and neural development associated genes. n=3 independent derivations of CFS and CS lines (marked CFS/CS_1-3) induced from different passages of parental H9 ES cells at passage range 55-60 (error bars represent the SEM for three cell line per condition). **D**, Phase contrast images displaying the distinct loose colony morphologies of H9 CFS and CS lines which are markedly distinct from parental H9 ES cells (scale bars: 50 µM). **E**, Representative images of CFS and CS lines (p26-28) immunostained for SOX2, T (BRA), CDX2, and PAX6 (scale bars: 40 µm). **F**, UMAP visualization of the single cell RNA-seq data of CFS_2, CS_2 (p14) cell lines, and H9 ES parental cells (10504, 12603 and 8671 single cells respectively), displaying NMP markers. **G**, RT-qPCR quantification of the telomere length relative to a 100 bp-long region on chromosome 17 in CFS_1-3, CS_1-3 lines and undifferentiated H9 ES cells over 10-20 passages. **H**, Phase-contrast images showing the dissection of the tail bud from a human embryo (PCW5) and the morphology of cells maintained in CFS medium for 10 passages. Cells at passage 10 immunostained for SOX2 and T (BRA). **I,** RT-qPCR analysis of HOX gene expression in CFS lines (p6-7 and p13-14 respectively) derived from PCW4.5 and PCW5 human embryos and in a CFS line generated from ePSC-H9 cells.

Focusing on signaling factors that are active during ePSC posterior differentiation, and in the NSB (Figure S1D), we found that sustained modulation of Wnt/β-catenin, FGF, and TGF-β signaling inhibition by CHIR99021, FGF2, and SB-431542 (abbreviated C,F, and S), respectively, yielded long-term self-renewing cell lines named CFS and CS that stabilized over 10 passages (Figure 1B). Extended passaging up to p27 revealed that CFS cell lines maintained stable high expression of *T (BRA)* and *CDX2*, whereas CS lines showed elevated *PAX6* expression. Both cell types expressed *SOX2* at levels comparable to undifferentiated H9 ePSCs, and markedly downregulated *OCT4* and *NANOG,* confirming their post-pluripotent state (Figure 1C). CFS and CS cell colonies exhibited homogeneous morphology and mutually exclusive expression of NMP- and NT-associated markers, as confirmed by scRNA-seq and flow cytometry (Figures 1D–F and S1E). Our screen also yielded CF and CH cell lines, but these were excluded from further analysis due to unstable phenotype during passaging (Figure S1F).

Given the role of telomere maintenance in sustaining the proliferative capacity of stem cells,^4^ we quantified telomere length across 20 passages. Using an assay measuring the telomere-to-single-copy reference ratio, we found that telomere length was stably maintained in both CFS and CS cell lines, comparable to H9 ePSCs (Figure 1G). Accordingly, CFS cells were clonogenic and could be expanded from single cells while maintaining NMP-associated markers (Figure S1G).

To validate the permissiveness of human embryonic NMPs for long-term self-renewal, we isolated tail bud cells from human embryos and plated them in CFS medium. We found that PCW4.5 and PCW5 embryos gave rise to stable CFS lines that could be propagated for more than 10 passages (Figure 1H). Like ePSC-derived CFS cells, embryo-derived CFS colonies displayed co-expression of SOX2 and T (BRA) together with similar colony morphology, and elevated expression of posterior *HOX* genes compared to H9-derived CFS. We conclude that CFS and CS cell lines capture developmental windows permissive for long-term self-renewal during human axial development.

### Mapping self-renewing CFS and CS cell lines to human and primate NMPs and downstream dorsal neural tube progenitors

To precisely map relationships between human CFS and CS cell lines and embryonic cells, we performed a multiomic characterization combining scRNA-seq and proteomic profiling (Figure S2A-D). In CFS cells, marker-based analysis revealed enrichment of canonical NMP transcription factors T (BRA), CDX2, and SOX2, together with axial progenitor markers including ETV5, EVX1, NKX1-2, and RARG (Figure 2A–B). ^14,22,42^ CFS cells also markers of expressed caudal neural progenitor, including NKX6-1, SP8, and FOXB1, MNX1, MSX1, GBX2, together with the segmentation clock regulator HES7.^43,44^ Consistently, CFS cells expressed nearly the full anterior-to-posterior HOX repertoire across HOX PGA–D (Figure 2C). Importantly, this CDX2-HOX signature was stably maintained across passages (Figures 2C and S3A). Expression of the most posterior HOX genes of PG11 – 13 increased with passaging of PSC-derived CFS cells (Figure 2C), while expression of these HOX genes was strong already at low passage of human tail bud-derived CFS cells (Figure 1I). Moreover, CFS cells expressed stemness factors including MYCN, LIN28A/B, and SALL4 (Figure 2B).

**Figure 2:**
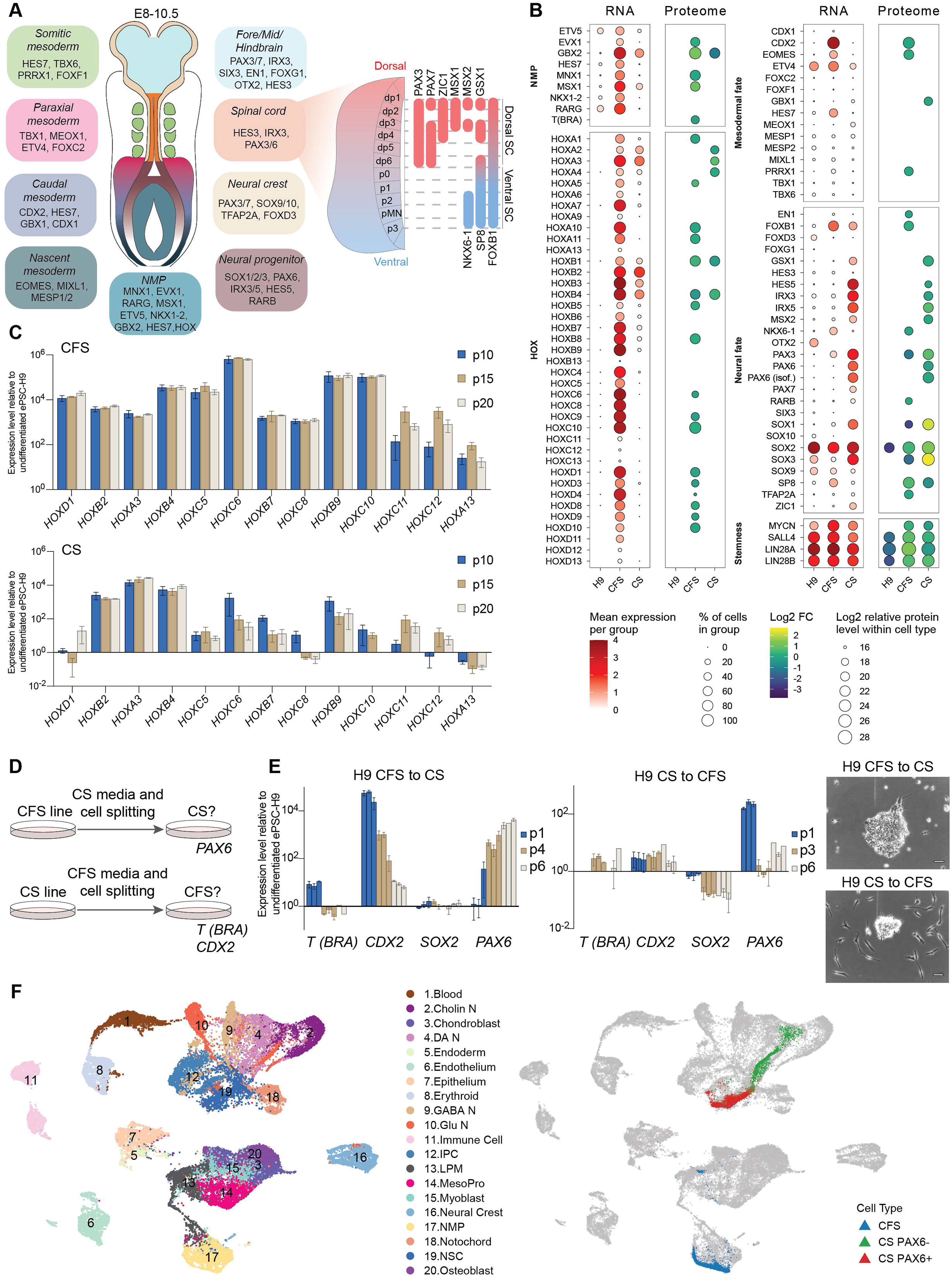
Embryonic annotation of human AxSC lines. **A**, On the left a model of an E8.5 mouse embryo illustrating the approximate locations of transcription factors expressed in NMPs and axial structures, including the neural tube and somitic mesoderm along the rostrocaudal axis, approximately mouse E9.5-E13.5, corresponding to human Carnegie stages 12-19.^14,22,43,73,74^ **B**, The expression of the markers categorized as per (**A**), is analyzed based on scRNA-seq data of human CFS_2, CS_2 and ePSC-H9 (10504, 12603 and 8671 single cells respectively) and proteome of three independently derived human CFS (1-3) and CS (1-3) lines, and two technical replicates of ePSC-H9 line. **C**, RT-qPCR analysis of three independently derived human CFS (1-3) and CS (1-3) for HOX genes (error bars represent SEM). **D**, A schematic representation of media swap experiments where CFS were transitioned to CS conditions, and reciprocally, CS were transitioned to CFS conditions. **E**, qRT-PCR analysis and cellular morphology during and following the reciprocal media swap experiments at passages 1-6 (depicted left and right). **F**, UMAP visualizations of the integration of scRNA-seq data of human CFS_2, CS_2 and ePSC-H9 (Figure 1F) into the human axis elongation dataset^9^ spanning PCW 3 to 12.

CS cells displayed expression of transcription factors that are associated with NT progenitor identity, including PAX6, SOX1-3, IRX3, IRX5, HES5 (Figure 2A–C). Additionally, neural patterning regulators such as NR2F1/2, and POU3F2/3, together with neuronal structural genes such as neurofilaments, were enriched (Figure S3A). The HOX expression profile of CS cells was more anterior than that of CFS cells, but cervical HOX genes were robustly expressed over dozens of passages (Figure 2C). This axial identity of CS cells was distinct from that of anterior neural progenitor cells (NPCs) generated from PSCs by dual SMAD inhibition^45^, which lacked the selective enrichment of HOX genes and posterior regulators such as MSX1/2 observed in CS cells (Figures S3B–D). Importantly, CS cells also exhibited enrichment of dorsal NT and neural crest transcription factors such as PAX7, PAX3, ZIC1, MSX2, and GSX1, together with stemness-associated factors shared with CFS cells (Figure 2B). Marker profiles were reproducible for CFS and CS lines derived from i/ePSC lines of distinct genetic backgrounds (Figure S3E–H).

The transcriptional programs of CS and CFS cells suggest a hierarchical model in which the CFS state is upstream to CS stem cells. To test this model, we transferred CFS cells to CS medium and vice versa (Figure 2D–E). CFS cell lines transitioned to CS morphology following 4-6 passages, downregulated *CDX2* and *T (BRA)*, and upregulated *PAX6*. No major changes in a panel of *HOX* or dorsoventral neural tube genes were observed between CS cells derived directly from H9 cells and those derived from CFS cells (Figure S3I). Conversely, CS cells transferred to CFS medium failed to upregulate *T (BRA)* and *CDX2* and did not downregulate *PAX6* expression. Instead, *SOX2* declined and the cells underwent spontaneous differentiation (Figure 2E).

To complement marker-based analyses, we mapped scRNA-seq profiles of CFS and CS cells onto reference datasets from gastrulating human and macaque embryos.^9,46^ CFS cells mapped to NMP populations in a PCW3 human reference, while PAX6^+^ CS cells mapped to neural progenitors (annotated as neural stem cells (NSCs) in this reference dataset, Figure 2F). Consistently, PAX6^−^ CS cells mapped along a trajectory extending from NSCs toward more differentiated neural lineages. Additionally, cosine similarity analysis showed that CFS cells exhibited the highest correspondence to PCW3 human embryos and E26 in cynomolgus macaques, whereas CS cells displayed comparable similarity to NSC populations across stages up to PCW12 (Figure S4A–B). Importantly, HOX gene expression pattern spanning anterior-posterior PG was comparable between CFS cells and human and primate NMPs at these stages (Figure S4C). Taken together, these results demonstrate that NMP- and downstream dorsal NT progenitor-like states can be captured as long-term self-renewing stem cell states while preserving their developmental identities and expressing stemness-associated factors.

### Differentiation of CFS and CS AxSCs into neural tube neuronal and glial lineages

Next, we analyzed the neural differentiation competence of CFS and CS states. We exchanged maintenance media for a cocktail of neurotrophic factors and analyzed differentiation through time-lapse imaging and time-resolved scRNA-seq (Figure 3A). Neuronal morphologies were observed earlier when starting from CS cells compared to CFS cells (3-4 days versus 7 days of differentiation, Figure 3B, Video S1). Consistently, differentiation of CFS cells passed through a CS-like gene expression state (Figures S5A–B). Furthermore, scRNA-seq indicated that by day 2 of differentiation approximately 80% of CS cells were in G_1_, compared with approximately 20% of CFS cells (Figure 3C). These results affirm the hierarchical positioning of CFS cells upstream of CS cells before they exit the cell cycle during further differentiation.

**Figure 3:**
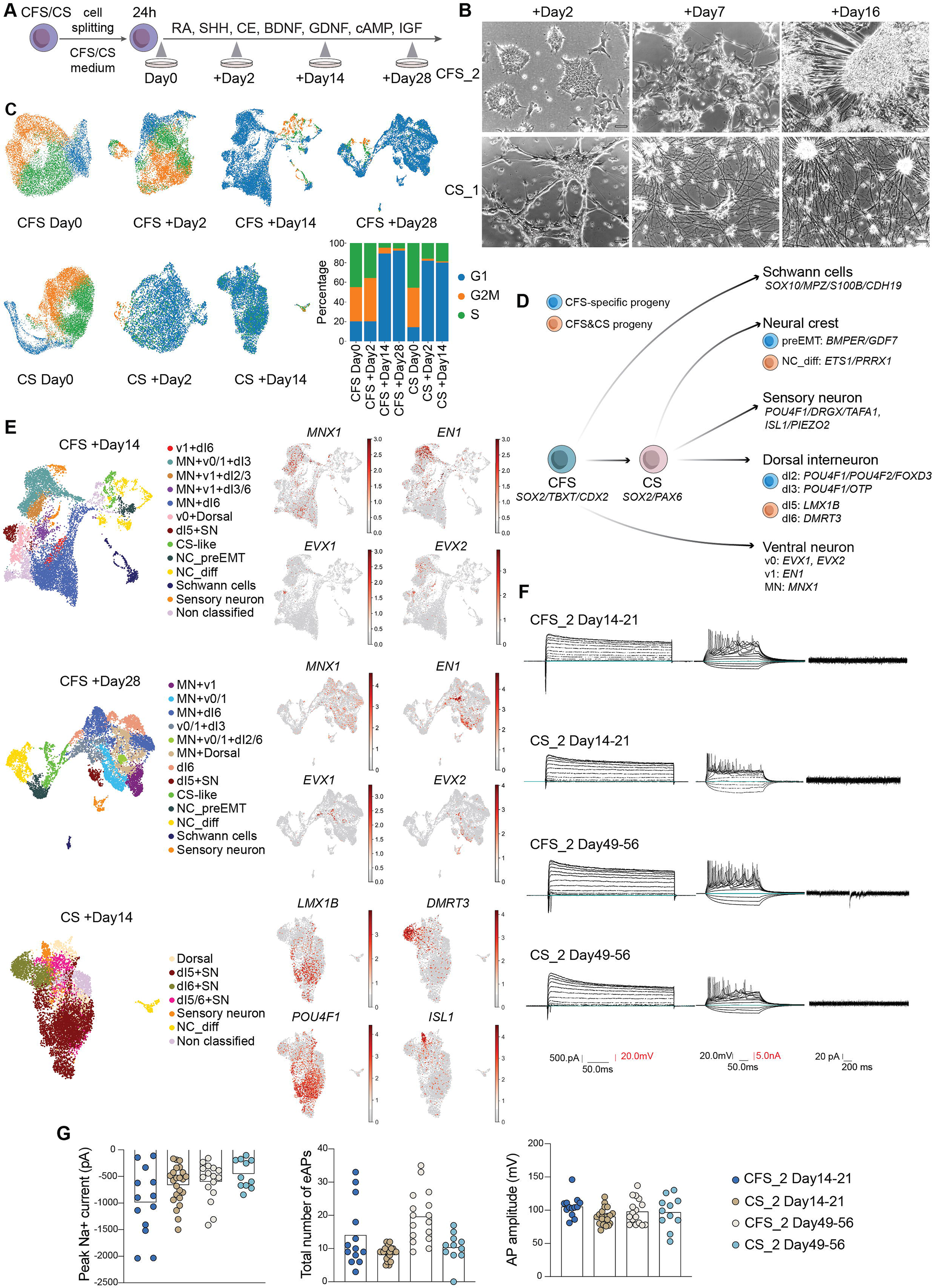
Functional neural differentiation of AxSCs. **A,B**, A schematic illustration of the neural differentiation protocol omitting NMP induction (A), and the corresponding phase contrast images at the respective time points (B) (scale bar: 50 µm). CFS_2 and CS_1 cell lines were dissociated and subject 10x chromium scRNA-seq at the indicated time points. **C**, UMAP analysis and quantification of G1, G2M and S cell cycle phases derived from the time course scRNA-seq data. **D**, An illustration depicting marker genes of the annotated cell types along the ventral – dorsal axis of the neural tube, and arrows connecting CFS and CS cell lines with the respective progeny (preEMT NC: pre-epithelial-to-mesenchymal transition neural crest cells, NC_diff: neural crest differentiation). **E**, Annotations of cell clusters in CFS Day14 (upper panel), CFS Day28 (middle panel) and CS DAY14 (lower panel) within the scRNA-seq data based on markers identified in **D** and Table S1. Clusters representing cells that display similar transcriptional signatures are labelled with the same coloring scheme across samples. Expression of ventral spinal cord neuron markers (*MNX1*, *EN1*, *EVX1*, *EVX2*) in CFS datasets, and dorsal spinal cord neuron markers (*LMX1B*, *DMRT3*, *POU4F1*, *ISL1*) in CS dataset are color labelled in corresponding UMAP plots (right panel). **F**, Representative current- and voltage-clamp data illustrating passive and active membrane properties of CFS_2 and CS_2 cells, at time points 2-3 weeks and 7-8 weeks into neuronal differentiation. Displayed are voltage clamp recordings (left column), current clamp recordings of evoked action potentials (AP, middle column) and spontaneous excitatory postsynaptic currents (right column). Currents in the left column were evoked in response to increasingly depolarizing voltage (steps of 20mV; 300ms). Fast-inward Na^+^ current (downwards) were followed by slow outward K^+^ currents (upwards). Voltages shown in the middle column were responses to both hyperpolarizing (downward) current injection and depolarizing (upward) current injection (steps of 20 pA; 300ms), showing multiple evoked APs in response to continued stimulation. The current trace in the right column reveals spontaneous excitatory postsynaptic currents during CFS cell differentiation, indicating spontaneous neural network activity with functional synapses. **G**, Summary of peak Na^+^ currents, AP peak amplitudes, and the total number of evoked APs per cell at each time point for measurements of 14-21 days CS, n=23; CFS, n=13; 49-56 days CS, n=11; CFS, n=16.

We then asked which specific neuronal cell types differentiated from CFS and CS states. Cell identities were annotated in scRNA-seq datasets according to markers expressed by ventral and dorsal human NT subdomains, including motor neurons, interneurons, neural crest progenitors, Schwann cells, and sensory neurons of the DRG (Table S1).^43,47^ Ventral neural tube identities were observed exclusively in CFS differentiation, with distinct clusters expressing V0-V1 interneuron markers, including *EVX1*-*2*, and *EN1*, as well as populations expressing the motor neuron marker *MNX1* (Figures 3E and S5C–D). In contrast, dorsal NT identities were shared between differentiated cultures derived from both CFS and CS states. These included dorsal interneuron (dI)2 and dI3-like identities expressing *POU4F1* (BRN3A), *POU4F2*, *FOXD3*, or *OTP*, which were predominantly generated from CFS cultures, as well as dI5- and dI6-like populations marked by expression of *LMX1B* and *DMRT3* expression, respectively, which were detected in differentiated cultures derived from both CFS and CS states (Figures 3E and S5D).

The NT commitment of CS cells was demonstrated in the higher abundance of two DRG sensory neuron subtypes, expressing *POU4F1* with either *DRGX*/*TAFA1* or *ISL1*/*PIEZO2*, relative to differentiated CFS cells (Figures 3D–E and S5D). In contrast, neural crest-derived Schwann cell markers *SOX10*, *MPZ*, *S100B* and *CDH19* were detected only in differentiated CFS clusters. Similarly, clusters expressing neural crest progenitor markers before delamination *GDF7* and *BMPER*, were observed exclusively in CFS differentiation (preEMT cluster). An additional G1/S enriched cluster expressing *ETS1* and *PRRX1*, was identified in both CFS and CS cultures, consistent with neural crest proliferation (NC_diff cluster; Figure 3E). Differentiation of selected identities from CS and CFS cells was further confirmed by qPCR and immunocytochemistry (Figure S6A–B). Together, these data support a hierarchical model in which CFS cells possess a broader developmental potential than CS cells, and transition through a CS-like state before generating dorsal NT fates (Figure 3D).

Analysis of HOX gene expression after 14 and 28 days of differentiation revealed broad maintenance of axial HOX identities, with only modest remodeling characterized by gradual attenuation of anterior HOX gene expression and a subtle increase in posterior HOX genes (Figure S6C–D). In contrast, CS cells after 14 days of differentiation underwent substantially greater progressive posteriorization, consistent with previous observations in NMP-derived neural progenitors.^33^ Importantly, these differentiated cells exhibited HOX profile that closely matched that of the human PCW3 neural tube.^9^

To determine the capacity of CFS and CS cells to generate functionally mature neurons, we performed whole-cell patch-clamp recordings. Voltage-clamp experiments at 2-3 and 7-8 weeks of differentiation revealed fast inward Na^+^ currents followed by slower outward K^+^ currents (Figures 3F and S7A). Both CFS- and CS-derived neurons generated reproducible action potentials upon depolarizing current injection (Figure 3F). Amplitudes approaching 100 mV within 2-3 weeks of indicated rapid functional maturation (Figure 3G). Comparable input resistance and resting membrane potential in the early and late differentiation stages further supported rapid electrophysiological maturation (Figure S7A). Additionally, neurons exhibited limited spontaneous network activity, consistent with the predominantly projection-neuron identity of peripheral neurons (Figures 3F). Similar results were obtained using CFS and CS lines generated from iPSCs, although maturation appeared slower than in H9 CFS lines (Figure S7B–C). To further assess functional synaptic connectivity, we cocultured human CFS-derived neurons with C2C12 mouse myotubes. These cocultures developed acetylcholine receptor (AChR) clusters adjacent to myotubes, indicative of neuromuscular junction formation (Figure S7D). Together, these results demonstrate that both CFS and CS are self-renewing undifferentiated progenitors that can be directed into mature neurons, neural crest and glia.

### Mesodermal potential of human CFS cells without tri-germ-layer teratoma formation

Expression of T (BRA), HES7, and CDX2 suggested that CFS cells possess mesodermal differentiation potential. To test this, we adapted a skeletal myocyte (SKM) differentiation protocol in which CFS cells were treated as paraxial mesoderm intermediates (Figure 4A).^48^ Indeed, differentiation of CFS cells, but not CS cells, led to a strong upregulation of SKM markers including *MYOD*, *CDH15* (M-cadherin), *MYOG*, and MYHC (myosin heavy chain) (Figure 4B–F). Reduced or absent expression of TBX6 and MSGN in CFS-derived multinucleated myotube-like cells was consistent with their early temporal window during SKM differentiation.

**Figure 4:**
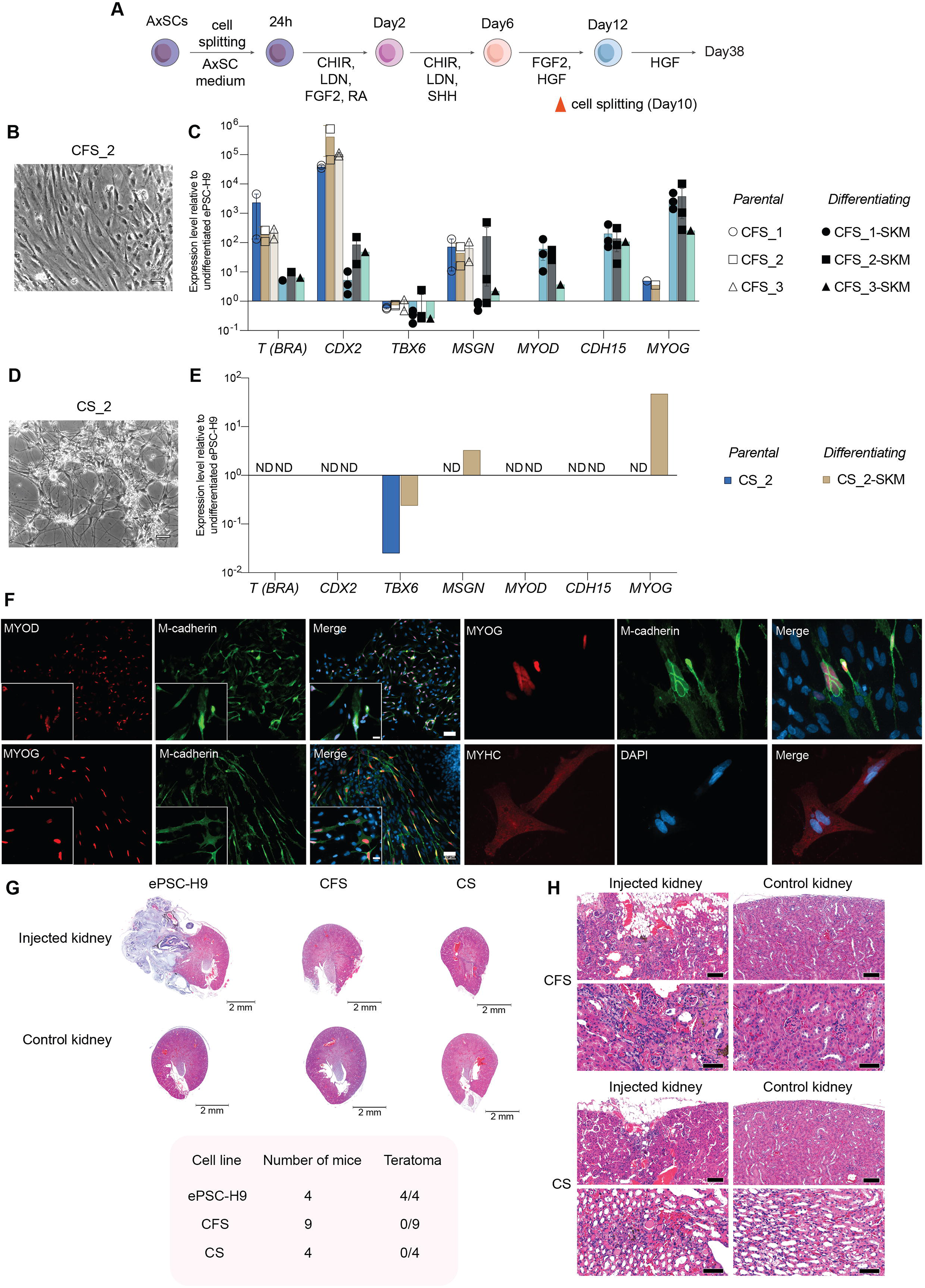
Skeletal muscle differentiation of CFS cells without teratoma formation. **A**, A schematic illustration of skeletal muscle differentiation omitting NMP induction. **B-E,** The phase contrast images of CFS (B) and CS (D) differentiation cultures at Day35-38 and Day35 respectively (scale bar: 50 µm), and corresponding RT-qPCR analysis (C, E) of NMP, somitic mesoderm, and skeletal muscle markers (SKM: skeletal muscle differentiation, ND: not determined, error bars represent SEM, scale bars: 50 µm and 20 µm for high and low magnification). **F,** Immunostaining and representative images of multinucleated cells in skeletal muscle differentiation of CFS cells at day 38 (scale bars: 50 µm and 20 µm for high and low magnification). **G-H**, Assessment of the teratoma formation potential of CFS and CS lines following injection to the kidney capsule of immunodeficient mice. Histological H&E images showing a tumor mass 4 weeks after the injection of control parental H9 ES cells (n=4), and tumor-free kidneys following H9 CFS and CS injection (n=9 and 4 respectively; scale bar: 2 mm (left) and 50 µm (right)).

Next, to assess broader developmental potentials, we injected CS (n=4 mice) and CFS (n=9 mice) cells under the kidney capsule of immunodeficient mice, using ePSCs as controls (n=4 mice). While teratomas comprising multiple tissue types formed in all ePSC-H9 injected animals (4/4), neither CFS (0/9) nor CS (0/4) grafts generated macroscopic teratomas (Figures 4G and S7E). Instead, grafts contained cells resembling cartilage and adipocytes, consistent with mesodermal and neural crest differentiation (Figure 4H). Together, these findings indicate that long-term self-renewing CFS cells, but not CS cells, retain paraxial mesoderm potential while both cell states lack pan-germ-layer developmental potential and avoid teratoma risk.

### Epigenetic remodeling during AxSC establishment reveals a developmental HOX timer

Having mapped the identity and differentiation potential of AxSCs, we next sought to interrogate the molecular mechanisms underlying their self-renewal. To identify genes that are essential for CFS maintenance, we performed a genome-wide CRISPR knockout screen targeting 18166 coding genes^49^ and analyzed the abundance of mutants at 6 and 16 days post-infection (Figure 5A). We quantified the levels of guide RNAs (gRNAs) within the initial and later cell populations and revealed that mutants of a total of 2,012 genes were significantly depleted across the two time points (Tables S2 and S5). Approximately half of these genes (1,028) were unique to the CFS state and not shared with the essential genes for human PSCs (Tables S2 and S5). Mutants depleted at day 6 were enriched for genes that were found to be associated with GO terms related to regionalization and the development of embryonic organ systems (Figure 5B). Thus, axial identity genes appear to be essential for CFS maintenance. Additional essential genes at this time-point mapped to the GO terms related to “epigenetic regulation of gene expression” and “DNA methylation”, indicating that CFS cells rely on a state-specific epigenetic configuration. At the 16-day time point, enriched GO terms were dominated by transcriptional and RNA-processing programs, suggesting that they are essential for sustaining the long-term fitness of the cells (Figure 5B). Consistent with the GO-term enrichment, network analysis across both time points grouped 1,553 essential genes into highly interconnected modules centered on transcriptional regulation, RNA processing, ribosome biogenesis, alongside nuclear-, epigenetic-, and metabolic-associated factors (Figure S8A and Table S2).

**Figure 5:**
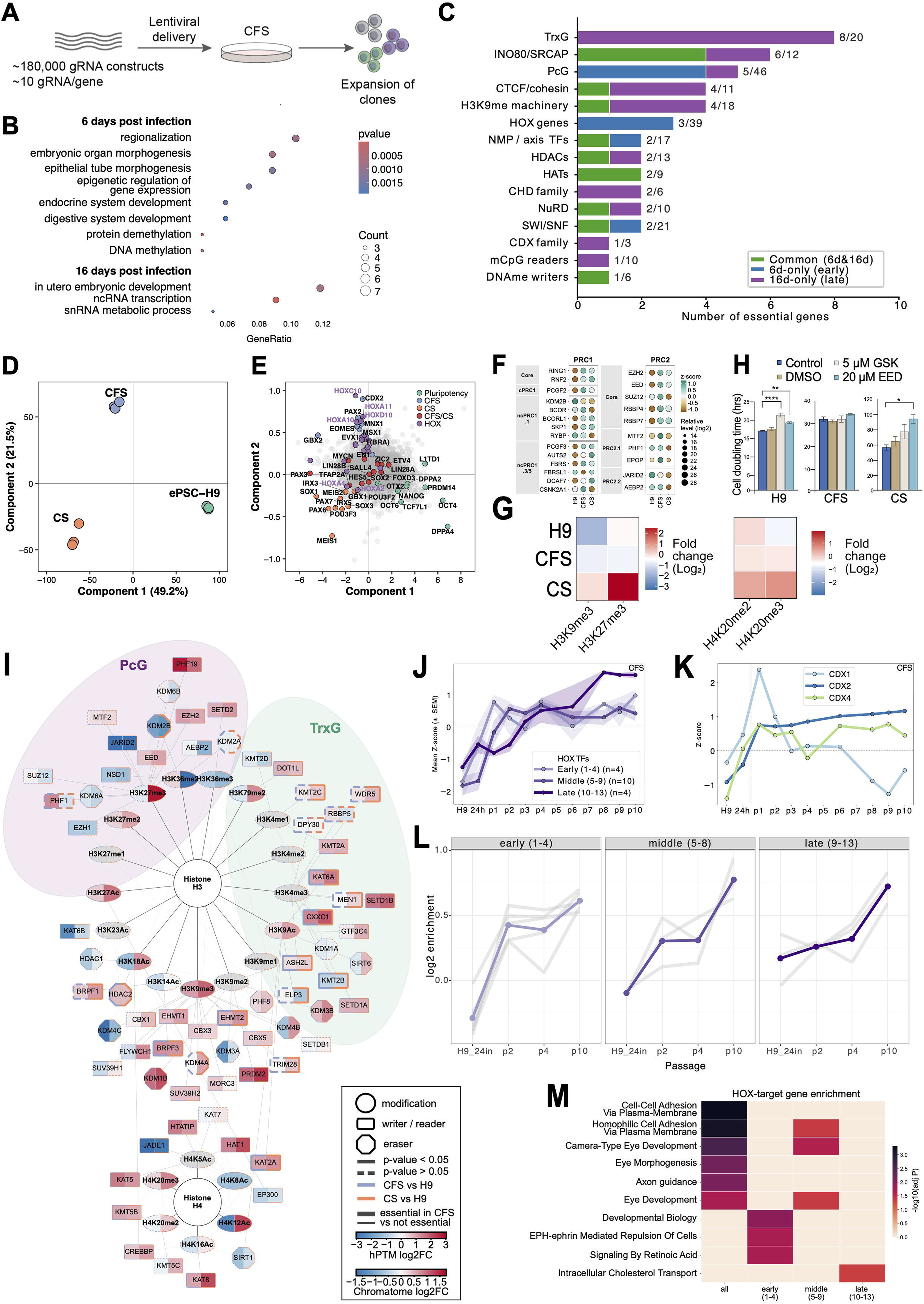
Chromatin and epigenetic regulation of AxSCs. **A**, A schematic illustration of the genome-wide CRISPR–Cas9 knockout (essentialome) screen in the human CFS_2 line. **B**, Gene Ontology enrichment of the essential genes at 6 days and 16 days post-infection; dot size reflects the gene count and shading the enrichment p-value. **C**, Number of essential genes per epigenetic and developmental category at day 6 and day 16 post-infection. Bar chart of CFS-essential genes from the genome-wide CRISPR essentialome screen, partitioned by manually curated functional groups. Bar height reports the number of category members scoring essential at each time point (Day6-unique n = 439, Day16-unique n = 1,074, common n = 499). Group definitions were validated against CORUM (v4.1) and EpiFactors (v2.1). **D**, Principal component analysis (PCA) based on chromatomes of ePSC-H9 (p61), CFS and CS AxSCs (p14-15) showing component 1 and 2 that together describe 70% of the variance. **E**, Individual loadings of PCA from (**D**) with markers coloured by category: pluripotency-associated, CFS-associated, CS-associated, shared CFS/CS, and HOX(-associated) proteins. **F**, Row Z-scored chromatin-bound abundance of PRC1 (left) and PRC2 (right) subunits across ePSC-H9, CFS and CS. **G**, Log2 Fold change of repressive histone H3 and H4 post-translational modifications after histone acid extraction and LC-MS/MS across ePSC-H9, CFS and CS. **H**, Cell doubling times upon treatment with PRC2 inhibitors GSK126 and EED226. DMSO treated and untreated cells were used as controls. **I**, Histone PTM and respective reader, writer and eraser protein network for CFS vs ePSC-H9 (purple lines) and CS vs ePSC-H9 (orange lines). Solid lines indicate a significant change while dashed lines indicate that a protein/PTM is not significantly more/less abundant compared to ePSC-H9. **J**, Chromatome trajectory of HOX transcription factors in the CFS time course, grouped by paralog position along the antero-posterior axis (early/anterior 1–4, n=4; middle 5– 9, n=10; late/posterior 10–13, n=4); mean row Z-score ± SEM across passages (H9, 24h CHIR pulse, p1–p10). **K**, Chromatome trajectory of the CDX paraHOX factors CDX1, CDX2 and CDX4 in the CFS time course; conventions as in (**J**). **L**, Enrichment of HOX motifs in hypomethylated regions across passages for CFS, with HOX proteins grouped by paralog position. Each grey line indicates a HOX protein, the colored line shows each group’s mean. Each point shows enrichment in the respective passage compared to H9. Included were HOX proteins whose motifs were found significantly enriched (p < 1e-3) in the hypomethylated (compared to H9) regions of passage p10. Derived from RRBS. **M**, GO analysis of genes associated with HOX motif-harboring hypomethylated regions for CFS, with HOX proteins grouped by paralog position. Hypomethylated regions of passage p10 were searched for motifs of the HOX genes shown in Figure 5L and annotated with the nearest gene. Panels J–M draw on the time-resolved chromatome and RRBS data (see Figure S9).

The strong representation of transcriptional and epigenetic regulators among essential CFS genes prompted us to investigate the epigenetic mechanisms underlying AxSC establishment and self-renewal. Notably, essential genes were highly enriched for Trithorax group factors (TrxG), but also included Polycomb group proteins (PRC), HDACs and HATs, together with components of the NuRD and SWI/SNF complexes (Figures 5C and S8B, Table S5). We thus performed a mass spectrometric analysis of chromatin-bound proteins (termed chromatome datasets; see Methods, Figure S2A-C, and Table S3) which captures functional regulatory states more directly than whole-cell proteomics (Figure S2E-G).^39^ PCA of the chromatome data revealed that PC1 (49.2% variance) separated ePSCs from CFS and CS states, whereas PC2 (21.5% variance) separated CFS from CS, indicating that chromatome differences between PSCs and both types of AxSCs are more pronounced than between CFS and CS states (Figure 5D and Table S5).

To examine molecular pathways that distinguish PSCs, CS, and CFS states, we next analyzed chromatome PC-loadings for the stable cell lines. Pluripotency factors, including OCT4, NANOG, DPPA4, and TCF7L1 (TCF3), were positively associated with PC1, suggesting that their downregulation enables the de-repression of lineage programs in AxSCs (Figure 5E). Transcription factors characteristic for NMPs, such as CDX2, and T (BRA) as well as HOX transcription factors were positively associated with PC2 and were specifically upregulated in CFS cells. In contrast, neural lineage regulators, including SOX1/3 and POU3F2/3, dorsal and neural crest–associated factors, TFAP2A and PAX3/6/7, together with retinoic acid–(RA) associated factors such as MEIS1/2, were uniquely associated with chromatin profiles of CS cells (Figure 5E and Table S5). These data also confirmed the chromatin association of stemness factors MYCN, LIN28A/B, and SALL4 in CFS cells (Figure 5E), with MYCN additionally identified as essential in the CRISPR screen (Figure S8A-B and Table S2).

We next focused on the PRC and TrxG machineries that read and write H3K27 and H3K4 methylation, respectively. Because activation of posterior HOX programs requires H3K27me3 removal^18^, we first asked how the PRC1/PRC2 complexes are configured across states (Figure 5F). Relative to PSCs, variant PRC1 components were enriched on chromatin in both CFS and CS cells. PRC2 components were most strongly enriched in CFS cells, while CS cells predominantly retained the core maintenance subunits EED, EZH2 and RBBP7 (Figure 5F and Table S5). Because the enrichment of PRC2 in the HOX-active CFS state was unexpected, we analyzed the PRC2 catalytic output in CFS cells. Profiling of the global histone PTMs by mass spectrometry (Figure 5G and S8D–G) revealed that H3K27me3 was lowest in CFS and highest in CS cells. This was accompanied by reduced H4 acetylation (H4K5/K8/K12/K16) in both AxSC states relative to PSCs together with modest gains of repressive H4K20me2/3 and H3K9me3 (Figures 5G and S8D–G,Table S5). Thus, despite comparable or higher PRC2 occupancy, CFS cells maintain low H3K27me3 HOX active state, whereas CS cells accumulate repressive chromatin features. To test the functional consequence, we pharmacologically inhibited the PRC2 subunits EZH2 and EED by GSK126 and EED226. This selectively impaired proliferation of CS but not CFS cells (Figure 5H), indicating that H3K27me3-dependent repression is required for CS self-renewal but not for CFS cells (Figure 5I and Table S5).

To identify mechanisms that potentially render CFS cells independent of PRC2 catalytic activity we next examined the temporal remodeling of PcG-TrxG balance during CFS and CS establishment. We performed a chromatome time course across ten passages of CFS and CS derivation from PSCs. Additionally, given the essentiality of DNMT1 (Figure S8B), we performed time matched reduced-representation bisulfite sequencing (RRBS) to track DNA methylation dynamics (Figure S9A, Table S4). The chromatome analysis detected 7,854–9,127 protein groups per sample, resulting in the deepest chromatin proteomes reported to date, enabling quantification of low-abundance chromatin factors (Figure S9B). It revealed that the cells progressed from the 24 h CHIR99021-posteriorized transient state toward stabilized CFS and CS states along PC1 (39.1% variance), with transient shifts along PC2 (12.8%; Figure S9C). In this, CFS and CS shared their PC1/PC2 trajectories but segregated along PC3 (7.7%) from passage 2 onward (Figure S9D), reinforcing their developmental relationship. Analysis of CFS and CS state-specific markers confirmed their identity and revealed passage-associated posteriorization evident by an increase in chromatin association of HOX factors, and accompanied by an early peak of CDX1 whereas the increase of CDX2 was gradual (Figures 5J–K and S9E). Importantly, while the chromatin association of H3K27me3 demethylases KDM6A/B were initially reduced during derivation, they regained elevated chromatin association in established CFS but not CS cells (Figure S9F). Thus, mechanistically, the dynamics of these demethylase potentially contribute to lower H3K27me3 in CFS despite higher PRC2 occupancy, and a more constrained chromatin state in CS cells which exhibited higher H3K27me3 abundance and PRC2 inhibition sensitivity.

Analyzing DNA-methylation factors for the chromatome data revealed increased chromatin association of DNMT1, UHRF1 and PCLAF with reciprocal loss of DNMT3A/B, alongside reduced TET1 and modest gains of TET2/3, yielding more balanced TET1/2/3 occupancy in the stabilized states (Figure S9G). This indicated a shift from *de novo* towards maintenance of DNA methylation during CFS and CS derivation.

To link these regulator dynamics to site-specific DNA methylation, we analyzed the time-resolved RRBS data. Regions demethylated from undifferentiated PSCs to stabilized AxSC states were enriched for HOX motifs, indicating progressive demethylation of HOX-target loci (Figure S9H). When resolved by HOX group, anterior targets (paralogs 1–4) demethylated first, followed by middle (5–8) and posterior (9–13) targets (Figures 5L and S9J), mirroring the step-wise posteriorization revealed by chromatin association of HOX factors (Figure 5J). Likewise, the sequential chromatin association increase of CDX1 followed by CDX2 was matched the temporally ordered hypomethylation of CDX motifs (Figure S9I). Conversely, motifs for the pluripotency regulators OCT4 and OCT4::SOX2 underwent step-wise hypermethylation during CFS posteriorization, consistent with progressive silencing of the pluripotency enhancer network upon exit from pluripotency (Figure S9L).^50^ Finally, HOX-motif-containing hypomethylated regions mapped to promoters of NMP/differentiation genes (Figure 5M).

Together, these observations indicate that sequential HOX activation is associated with a temporal CDX relay that precedes ordered HOX chromatin engagement and progressive demethylation of HOX-target loci. More broadly, these data support a model in which AxSC capture recapitulates a developmental CDX–HOX DNA-demethylation timer that operates during human axial development, coordinated by PRC-TrxG activity, transcription factor networks, and resulting in histone PTMs that promote progressive activation of posterior HOX programs. Consistent with this model, human embryo tail-derived CFS cells, derived from the most posterior NMP CNH state, stably maintain both anterior and posterior HOX expression during long-term self-renewal (Figure 1I).

### Metabolic reprogramming establishes distinct OXPHOS–glycolytic states during AxSC capture

A class of essential genes identified by the CRISPR screen encoded components of mitochondrial energy metabolism, suggesting that oxidative phosphorylation is a core requirement for AxSC maintenance (Figure S8C and Table S2). We therefore asked how metabolic profiles of AxSCs differ from those of PSCs, and whether further metabolic specialization occurs between CFS and CS states. Unsupervised clustering of differentially expressed proteins identified two clusters which were upregulated in CFS and CS cells and that were enriched for proteins associated with the TCA cycle, acetyl-CoA metabolism, and mitochondrial biogenesis (clusters 3 and 4, Figure 6A, Table S6). To determine how these protein changes were reflected at the metabolic level, we performed bioenergetic profiling using Seahorse assays (Figure 6B). While PSCs and CS cells showed similar levels of basal glycolysis and glycolytic capacity, CFS cells displayed a small but significant reduction in basal glycolysis compared to PSCs, and lower glycolytic capacity than CS cells (Figure 6C). Consistent with these functional data, glycolytic enzymes were enriched in CS relative to CFS cells (Figure 6D). Basal respiration, ATP production and maximal respiration, in contrast, were elevated in both CFS and CS cells compared to PSCs, with CFS cells showing higher maximal respiration than CS cells (Figure 6C). Accordingly, key mitochondrial metabolic proteins were upregulated in both CFS and CS cells relative to PSCs, including the pyruvate dehydrogenase (PDH) complex components PDHA1 and PDHB, the TCA cycle enzymes CS, OGDH, and SDHA as well as ACLY, which links mitochondrial citrate metabolism to acetyl-CoA production (Figure 6B, E). These results suggest that both CFS and CS cells rely more strongly on oxidative phosphorylation (OXPHOS) than PSCs, with CS cells maintaining a mixed OXPHOS–glycolytic metabolism.

**Figure 6:**
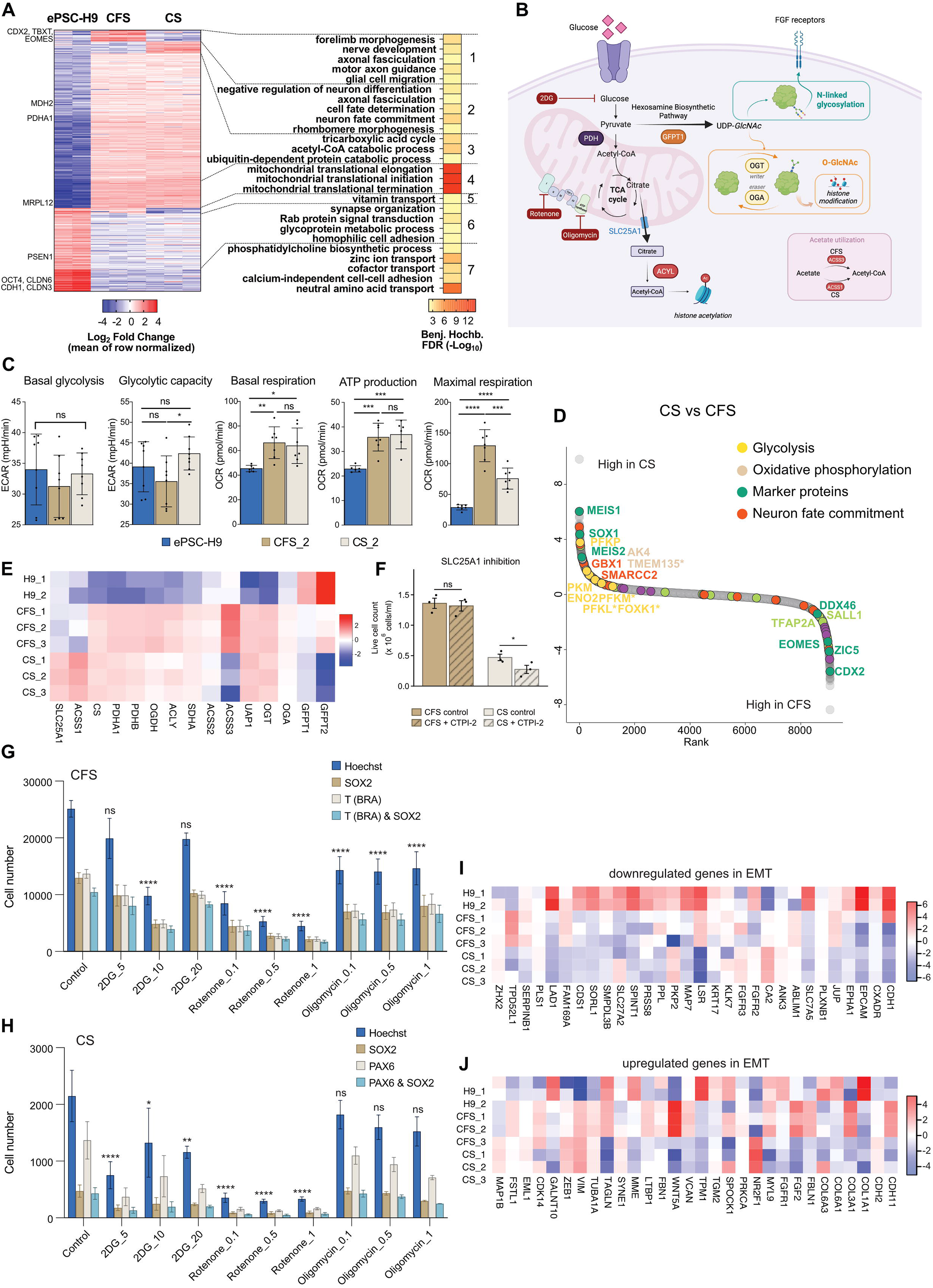
Metabolic reprogramming of human AxSCs. **A**, A heatmap displaying seven clusters of differentially expressed proteins based on a global proteome analysis of CFS_1-3, CS_1-3 and ES cells H9. Selected subset of the GO terms corresponding to the respective clusters are showed. **B**, Schematic overview of the hexosamine biosynthetic pathway, glycolysis, and oxidative phosphorylation with corresponding inhibitors. **C**, Figure comparing the extracellular acidification rate (ECAR) and oxygen consumption rate (OCR) to pinpoint metabolic differences and energy production pathways in CFS_2, CS_2 and H9 ES cells. ECAR and OCR rates indicate glycolytic metabolism, and mitochondrial oxidative phosphorylation metabolism together with non-mitochondrial respiration, respectively. ECAR and OCR are normalized by cell number. Error bars represent SEM; unpaired t-tests. ns, not significant; *, p<0.05; **, p<0.005; ****, p<0.0001. **D**, Protein rank based on the relative log2 fold change with respect to global proteomes of CFS_1-3 versus CS_1-3. The annotation of proteins is based on CFS and CS markers and GO term categories. **E,** A heatmap displaying the expression of key genes involved in the hexosamine biosynthetic pathway, glycolysis and oxidative phosphorylation. **F**, Quantification of live CFS and CS cell numbers at passage 2 following continuous treatment with 30 μM CTPI-2. **G-H,** The number of unstained cells, SOX2-stained cells, T (BRA)-stained cells, and SOX2 & T (BRA) double-stained CFS cells, and SOX2-stained cells, PAX6-stained cells, and SOX2 & PAX6 double-stained CS cells measured following treatments with 2DG to inhibit glycolysis, and rotenone and oligomycin to inhibit oxidative phosphorylation. Statistical analysis compares unstained cell counts across different treatments. Error bars represent SEM; ns, not significant; *, p<0.05; **, p<0.005; ****, p<0.0001. **H-I**, Heatmaps displaying expression of upregulated and downregulated genes during EMT process in global proteome of CFS_1-3, CS_1-3 and ES cells H9.

Metabolic activity can influence epigenetic and signaling pathways by supplying precursor molecules for posttranslational protein modifications. In particular, mitochondrial citrate metabolism provides acetyl-CoA for histone acetylation (Figure 6B).^37,51^ To determine whether mitochondrial metabolism supports functions beyond ATP production, we compared the effects of inhibiting complex I with Rotenone and ATP synthase with Oligomycin. Both CFS and CS cells were sensitive to Rotenone, whereas Oligomycin had a more limited effect (Figure 6F–G), suggesting that mitochondrial metabolism supports biosynthetic functions in addition to ATP production. Consistent with this interpretation, CS cells exhibited elevated histone H4 acetylation (Figure S8E, G). This was accompanied by increased expression of the mitochondrial citrate transporter SLC25A1 (Figure 6E), suggesting enhanced citrate export to the cytoplasm and expansion of the acetyl-CoA pool available for histone H4 acetylation relative to CFS cells. Accordingly, inhibition of this transporter by CTPI-2 lowered the proliferation of CS cells but not CFS cells (Figure 6F).

Tailbud elongation in mouse and chick embryos, and cell fate decisions in mouse gastruloids are sensitive to glycolysis inhibition.^52–56^ Glycolytic activity has also been proposed to play a role in human cells, although the metabolism was not directly analyzed.^57^ A potential molecular mechanism for glycolysis sensitivity is the hexosamine biosynthetic pathway (HBP) that routes glycolytic intermediates toward UDP-GlcNAc production, which in turn promotes FGF signaling through protein glycosylation (Figure 6B).^58–60^ When examining the expression of HBP pathway components and downstream enzymes, we found that the rate-limiting factors GFPT1 and GFPT2 were enriched in PSCs and CFS cells, which rely on FGF signaling for self-renewal (Figure 6E). Moreover, UAP1 and OGT, which catalyze flux through O-GlcNAc cycling, a pathway that promotes cell delamination,^61^ were enriched in CFS and CS cells, while OGA that promotes O-GlcNAc cycling, was comparably expressed in all states. In accordance, inhibition of glycolysis by 2DG impaired the survival but not stage specific markers of both cell types (Figure 6G-H). Finally, examination of epithelial-to-mesenchymal transition-(EMT) associated factors in protein clusters 6-7 that were downregulated in CFS and CS cells (Figure 6A) revealed reduction of epithelial adhesion molecules, including CLDN3, CLDN6, and CDH1, but not an increase of canonical transcription factors SNAI1/2 and TWIST1 (Figure 6I-J, Video S2). Together, these findings suggest that glycolysis fulfills distinct stage-specific functions during AxSC establishment, transitioning from supporting FGF-dependent stem cell maintenance to promoting O-GlcNAc– associated neural crest competence and delamination.

### Derivation of mouse CFS cells and their axial chimeric potential

We next asked if post-implantation AxSC lines could be derived from i/ePSCs of other mammals, focusing on mouse EpiSCs. Because mEpiSCs express primitive streak/tail bud markers like *T (Bra)* under primed culture conditions,^62,63^ we omitted the CHIR99021-induced posteriorization step and directly screened candidate media (Figure 7A). Starting with the CFS media formulation that was successful for human PSCs and tail bud embryos, we varied concentrations of CHIR99021, FGF2, SB-431542, and the BMP signaling inhibitor DMH1 (Figure S10A-B). We also tested FGF8 as a potential self-renewal factor^64^ and mouse embryonic fibroblast (MEF) feeders compared to Matrigel coating (Figures 7A, S10A–B). We obtained stable lines self-renewing for 10 or more passages in the same media used for human CFS (huCFS) cells with MEFs, as well as with lowered CHIR99021 and FGF2 concentrations (3 µM instead of 5 µM and 50 instead of 100 ng/ml, respectively). Presence of 10 µM SB-431542 was essential for mouse CFS (mCFS) maintenance, whereas DMH1 was dispensable, and FGF8 could not replace FGF2 (Figure S10A-B). Stable lines were obtained with Matrigel coating only with concentrations of 3 µM CHIR99021, 50 ng/ml FGF2 and 10 µM SB-431542. In all conditions that supported long-term self-renewal, cell lines expressed *T (Bra)*, *Cdx2*, and *Sox2*, but not *Oct4*, *Nanog*, and *Pax6*, similar to huCFS lines (Figures 7B and S10A-D). No self-renewing mCS-like cell lines were obtained.

**Figure 7:**
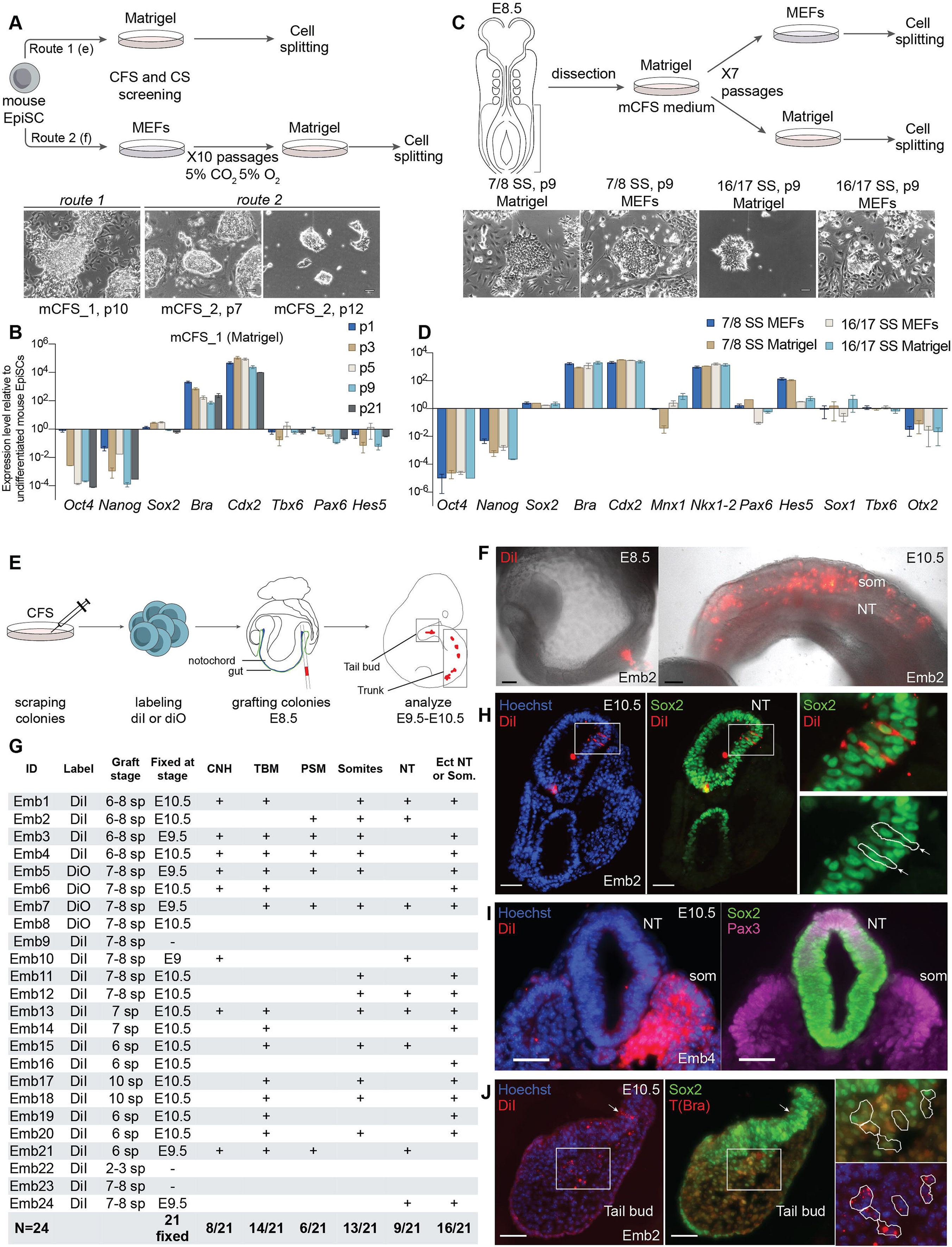
Derivation of chimera competent mouse AxSCs from PSCs and embryos. **A**, A schematic representation of a strategy for protocol screening for the derivation of mouse posterior stem cells from EpiSCs (p38-p43, E14.1 mESC line). Cells were passaged by single cell splitting to Matrigel (route 1, Figure S10A) or inactivated MEFs followed by transfer to Matrigel at p10 (route 2, Figure S10B). Phase contrast images of mouse CFS cell lines successfully derived on both Matrigel (mCFS_1) and MEFs-Matrigel (mCFS_2) are shown. **B**, RT-qPCR analysis of a representative mCFS_1 line using pluripotency markers, NMP and paraxial mesoderm development associated genes (error bars represent SEM). **C**, A schematic illustration detailing the strategy for dissection of the CLE region of E8.5 mouse embryos and grouping according to somite stage 7/8 and 16/17. Cells were dissociated and plated on Matrigel in mouse CFS medium as mCFS_1 line. After p7, cells were passaged either on Matrigel or MEFs. Phase contrast images showing the distinct colony morphologies of CFS cells derived from 7/8 and 16/17 somite stage (SS) embryos (scale bars: 50 µM). **D**, RT-qPCR analysis of mouse CFS lines (p11) derived from 7/8 and 16/17 SS embryos, using pluripotency markers, NMP and paraxial mesoderm development associated genes (error bars represent SEM). **E**, An illustration depicting the microcapillary ting of fluorescently labeled (DiI or DiO) undifferentiated mouse CFS cells (Figures 7A, B) into E8.5 mouse embryos in the node-streak border (NSB) region, and regions where fluorescently labeled cells were detected at the 24h or 48h follow-up time points. **F**, Representative images showing the transplantation site of mouse CFS cells labeled with DiI (E8.5) and incorporation into somites (som) and neural tube (NT) at the E10.5 follow-up. **G**, A table summarizing the number of embryos transplanted at E8.5, and observed DiI or DiO signal in axial structures, as follows, somites (som), neural tube (NT), presomitic mesoderm (PSM), chordo-neural hinge (CNH) and tailbud mesoderm (TBM) (Emb: embryo, Ect: ectopic, sp: somite pairs). Additionally, the notochord was labelled in 2 embryos, and labelled putative neural crest cells were observed in an additional 2 embryos. In 1 embryo a clump of labelled cells was present in the hindgut lumen. **H**, A transverse section of embryo 2 showing DiI labelled cells (red) in the neural tube, and Sox2 (green) immunostaining of the same section, and an overlay image. The white outline shows the position of DiI-labelled cells, noting that the Sox2 signal in the grafted cells was at a similar level to the endogenous host cells. **I**, A transverse section of embryo 4 showing DiI label in one of the somites. Maximum intensity projections of z-stack images taken from the same section with double-immunostaining of Sox2 (green) and Pax3 (magenta). **J**, A transverse section of the tail bud of embryo 2 showing DiI labelled cells in the epithelium (arrowhead) and mesenchyme (lower panel, right). Maximum intensity projections of z-stack images taken from the same section double-immunostained for Sox2 (green) and Brachyury T (red). DiI-labelled cells are outlined in white, noting that the signal of Sox2 and T were similar to neighboring endogenous unlabeled host cells.

Motivated by the successful derivation of huCFS cells from human embryonic tail tissue (Figure 1H), we attempted to establish mCFS cell lines from the stem zone of 7/8 and 16/17 somite-stage mouse embryos in the presence or absence of MEFs (Figure 7C). mCFS lines were obtained from both somite stages, exhibiting characteristic NMP-like genes, with 7/8-somite derived lines displaying a more pronounced epithelial morphology (Figure 7C–D).

Mouse embryonic NMPs self-renew and differentiate upon grafting in cultured E8.5 embryos.^10,65^ We therefore used this assay to assess the multilineage potential of mCFS cells *in vivo*. EpiSC-derived mCFS cells were labeled with lipophilic dyes (DiI or DiO) and grafted into the NSB/CLE region of E8.5 mouse embryos (n=24) (Figure 7E). At both 24h and 48h (E9.5 and E10.5, n=21 embryos), graft-derived cells contributed to the presomitic mesoderm (6/21 embryos), somites (13/21 embryos) and NT (9/21 embryos) (Figure 7F–G). Multilineage contribution was evident by presence of Sox2⁺ DiI⁺ cells in the neural tube, and of Pax3⁺ or Meox1⁺ Sox2⁻ DiI⁺ cells in somites (Figures 7G–I and S10E). At the later tailbud stage (E10.5), DiI⁺ Sox2⁺ Brachyury⁺ cells were observed intermingled with endogenous NMPs in the CNH, suggesting that grafted mCFS cells remained in the late NMP niche (8/21 embryos) (Figure 7J). Moreover, in 16 embryos, we identified CFS cell-derived ectopic neural tube or somite structures (Figure 7G, S10F). These results suggest that long-term self-renewing mCFS cells possess a similar multilineage competence as embryonic NMPs. In addition, our inability to establish stable mCS lines points to an evolutionary divergence in the organization of posterior progenitor populations between mouse and human.

### Transcriptomic landscape reveals a hybrid AxSC state in mCFS cells

To define the basis for divergence of posterior progenitors between human and mouse, we performed scRNA-seq of mCFS lines derived from EpiSCs and from somite-stage 7/8 and 16/17 embryos (Figure 8A). EpiSC- and embryo-derived lines expressed *Sox2* and *Cdx2* homogeneously, and were largely negative for *Pax6* and *Mnx1* expression. *T* (*Bra*) expression was heterogeneous and somewhat more pronounced in embryo-compared to EpiSC-derived lines (Figure 8A). Mapping onto a time-resolved atlas of mouse gastrulation revealed highest transcriptional similarity to 7.5-8.5 mouse NMPs (Figures 8B–C and S11A). All three mCFS lines exhibited similar *Hox* expression profiles extending up to PG11, recapitulating and extending beyond those observed in embryonic mouse NMPs at E8.5 (Figure S11B). This indicates close correspondence of the self-renewing mCFS cells to axial progenitors *in vivo*. Additionally, similar to human CFS and CS states, mCFS cells expressed stemness factors MYC, SALL4, and LIN28A/B (Figure 8D).

**Figure 8:**
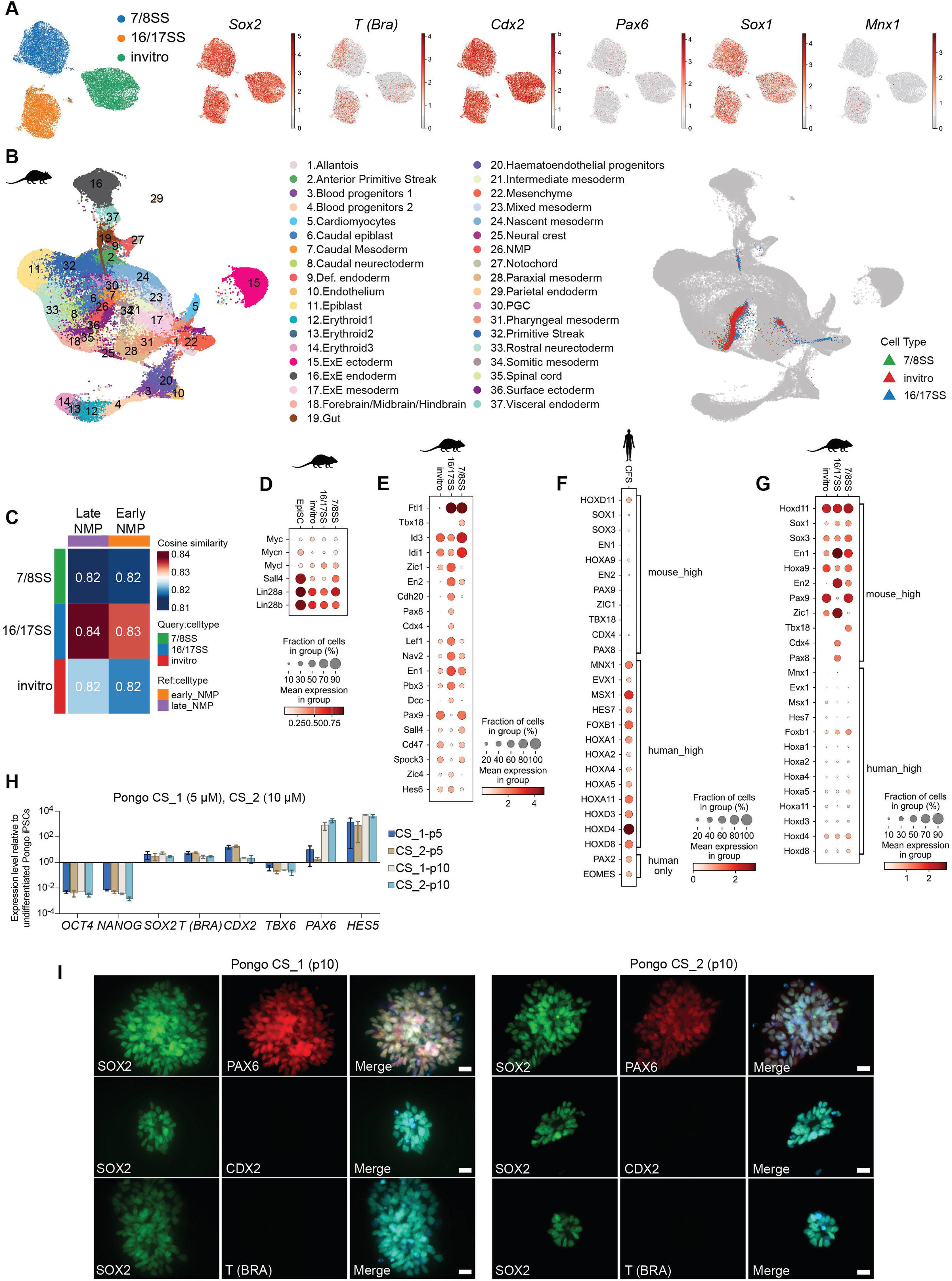
Embryonic annotation of a single state of mouse AxSCs. **A**, UMAP visualization of the single cell RNA-seq data of mouse CFS cell lines including *in vitro* derived (mCFS_1), and 7/8 SS and 16/17 SS (Matrigel) embryo derived lines. UMAPs displaying the expression of NMP and neural progenitor associated genes. **B,** UMAP visualization of the integration of scRNA-seq data of *in vitro* mCFS_1, and embryo derived 7/8 SS (Matrigel) and 16/17 SS (Matrigel) lines into the mouse gastrulation dataset^73^ spanning E7.5 to E8.5. **C**, A heatmap showing similarity of the mouse CFS lines to early and late NMPs identified by Pijuan-Sala et al^73^. **D,** A dotplot displaying expression of stemness-related genes in mouse CFS lines and mouse EpiSCs.^75^ **E**, A dotplot displaying expression of genes selected from DEG analysis. **F-G**, A dotplot showing the expression of genes that vary in human CFS_2 (F) and mouse CFS (G) datasets. **H-I**, RT-qPCR (H) and representative immunofluorescence images (I) used for characterization of Pongo CS_1-2 lines derived from Sumatran orangutan (*Pongo abelii*) female iPS cells (p: passage number, error bars represent SEM, and scale bars: 20 µm).

However, comparison between human and mouse CFS cells revealed substantial differences. Mouse CFS cells expressed the early neural transcription factors *Sox1* and *Sox3*, which in human are enriched in the CS state (Figure 8A, E–G). Moreover, they expressed *En1* and *En2*, transcription factors associated with posterior neural tube and hindbrain patterning (Figure 8G). In contrast, the mesodermal segmentation clock gene *Hes7*, as well as neural progenitor and patterning transcription factors, *Msx1*, *Mnx1*, *Evx1*, and *Foxb1*, were depleted (Figure 8A, E–G). Consistent with this immature neural gene expression signature, the second highest transcriptional correspondence of mCFS cells upon reference mapping was to embryonic ectoderm clusters, including spinal cord, and forebrain, midbrain and hindbrain (FMH) progenitors (Figure S11A). Together, these findings suggest that mCFS cells occupy a transcriptionally hybrid state showing features of both human CFS and CS cells but with a less pronounced expression of patterning genes. Further evidence for the resemblance to human CFS cells came from the expression of metabolic genes, which showed significant scores for glycolysis- and OXPHOS-associated genes (Figure S11C).

Finally, to test if a separate program for self-renewal of NT progenitors existed in other primate species, we applied CS culture conditions to iPSCs of the Sumatran orangutan (*Pongo abelii*). This resulted in the derivation of cell lines phenotypically similar to human CS lines (Figure 8H-I). This corroborates the idea that inability to derive mouse CS cells, and the neural priming of mouse CFS cells, reflects a primate-murine evolutionary and developmental differences.

## DISCUSSION

Here, we establish AxSCs as a new class of developmental stem cells positioned between pluripotent and tissue-specific stem cells. While embedded within progressive axial patterning programs, they achieve long-term self-renewal while maintaining multilineage developmental competence. Our findings further raise the possibility that additional transient axial progenitor populations, including notochord progenitors, may also be captured as stable self-renewing stem cell states.^11^

Mechanistically, capture of AxSCs rests on suspending a developmental timer that starts running once PSCs leave their pre-implantation state. Molecularly, this timer is driven by a relay of TrxG-dependent H3K27me3 demethylation and a CDX-HOX regulatory chain leading to site-specific DNA demethylation. These events are integrated with a global reprograming of chromatin and metabolic pathways. We propose that during human embryonic development, this timer prepares cells for assuming progressively more posterior identities in axial elongation, although direct validation of this model warrants further investigation.

In contrast to cells in the embryo, which proceed to specialized neural and mesodermal identities, the signaling environments that we identify pauses this developmental timer and thus stabilizes human AxSCs as post-pluripotency stem cells. We note that CS and CFS cells differ in their HOX expression profile, and that human tail-derived CFS cells express more posterior HOX identities than PSC-derived lines. Thus, depending on signaling environment and starting cells, multiple stages of the axial developmental timer can be suspended in self-renewing stem cell states.

The mechanisms sustaining AxSC self-renewal remain incompletely understood. Our data suggest that AxSCs combine lineage-specific developmental programs with a stemness circuitry that is present in pluripotent cells, including SOX2, MYC family transcription factors, LIN28A/B, and SALL4.^49,66,67^ In addition, sustained telomere maintenance during long-term culture likely provides a complementary mechanism supporting the extensive proliferative capacity of AxSCs. AxSC self-renewal may also depend on as-yet-unidentified autocrine signaling mechanisms that emerge upon AxSC capture. AxSC self-renewal as well as their specific developmental states may further be supported by the bioenergetic remodeling associated with their derivation that feeds into epigenetic and signaling systems. Increased OXPHOS in both CFS an CS cells, for example, will lead to higher mitochondria acetyl-CoA levels. Stronger expression of citrate export enzymes in CS compared to CFS cells may then lead to an increase in cytosolic Acetyl-CoA, increased histone H4 acetylation, and a narrower differentiation potential. In parallel, expression of rate limiting enzymes for the HBP decreased from PSCs over CFS to CS cells, whereas expression of O-GlcNAc cycling enzymes progressively tended to increase. While the HBP pathway shunts of glucose-derived metabolites to glycosylation pathways that promotes FGF signaling, O-GlcNAcylation promotes cell delamination.^60^ Progressive shifts in the activity of these different metabolic pathways may therefore help stabilizing specific developmental identities through promoting distinct sets of protein post-translational modifications.

In humans, we derived two distinct self-renewing post-implantation stem cell states: CFS cells, corresponding broadly to NMP-like identities, and CS cells, corresponding to dorsal neural tube progenitor-like identities. These states extend previously described transient human NMP and neural differentiation models by enabling long-term self-renewal.^32^ While neural stem cells (NSCs) constitute CNS progenitors that persist into adulthood,^68^ CS cells represent an earlier developmental state that retains competence to generate neural crest derivatives, including dorsal root ganglion (DRG) neurons. Mechanistically, although CFS and CS cells are transcriptionally closely related, they are epigenetically and functionally divergent. Histone PTMs and PRC2 inhibitor sensitivity indicate that CS cells adopt a more restricted chromatin state, whereas CFS cells maintain a permissive epigenetic configuration that preserves neuromesoderm plasticity.

In contrast to the two AxSC states captured with human cells, the same derivation approach led to the stabilization of only a single state from mouse PSCs and embryos. This state combines transcriptional features of both human CFS and CS cells while exhibiting incomplete activation of lineage-specification and patterning programs. Although we cannot exclude the possibility that discrete states with human-like CS and CFS properties can be captured from mouse cells under alternative conditions, a species-specific organization of posterior progenitor domains offers an alternative attractive explanation for the existence of a single hybrid AxSC state in the mouse. Such divergent organization has been described between chick and mouse embryogenesis, where mouse NMPs occupy the growing caudal edge of the neural tube, whereas NMP and neural tube progenitors have been described as distinct populations in chick embryos.^69^ Our ability to capture two AxSC states with distinct developmental potencies might indicate that the organization of posterior progenitors in human embryos resembles that of the chick more than that of the mouse.

The long-term self-renewal, scalability and cellular homogeneity of AxSCs represent key advantages compared to transient i/ePSC-derived NMPs and neural tube progenitors. Providing more direct and efficient access to defined lineages, AxSCs are poised to become a versatile platform for investigating mechanisms of posterior body formation in human, including body segmentation, tail formation, and tail regression.^70^ From a translational perspective, AxSCs provide scalable access to spinal cord, neural crest, and DRG derivatives for disease modeling, drug discovery, and studies of drug–cell interactions. Thanks to their long-term self-renewal and lack of teratoma formation following transplantation, AxSC-derived products may offer a safer regenerative medicine strategy than PSC-derived counterparts for disorders affecting the peripheral nervous system such as Hirschsprung disease.^71,72^

**Figure S1: A-B**, Schematic representation of the bulk RNA-seq (**A**) data analysis and representative genes (**B**) following CHIR99021 treatment and doxycycline activation of β-catenin ΔN90 integrated in human H9 ES cells to pinpoint the temporal point of NMP induction. Note the upregulation of primitive streak *T* (*BRA*) and posterior embryo *CDX2* transcription factors at 24h, followed by the downregulation of *SOX2*. Additionally, there is a steep rise in mesoderm and endoderm-associated transcription factors and morphogens, including *TBX6*, *FGF17*, *BMP4*, and *GATA6* between 24-48h. **C**, A heatmap displaying upregulated genes and developmental associations, i.e. ectoderm, mesoderm, neural tube and neural crest, analysed using Enrichr tool and JENSEN tissue expression database. Expression was normalized to untreated undifferentiated cells (n=2). **D**, An analysis of the over-represented signaling pathways in the differentiation time course. Activated pathways at 24h were the basis for screening pathway agonists and antagonists of Wnt/β-catenin, FGF, TGF-β, and BMP pathways. **E**, Representative FACS plots showing CFS (left) and CS (right) cells analyzed for SOX2 and T expression. **F,** The characterization of human CH and CF cell lines, derived by 5 µM CHIR99021 (CH), or 5 µM CHIR99021 and 100 ng/ml FGF2 (CF) via RT-qPCR using pluripotency markers, NMP and neural development associated genes (error bars represent SEM). **G,** RT-qPCR analysis of CFS cells and two isolated single clones from bulk CFS culture for pluripotency markers, NMP and mesodermal development associated genes (error bars represent SEM).

**Figure S2: A**, Chromatome samples were prepared by Chromatin Aggregation Capture (ChAC) followed by data-independent acquisition (DIA). **B**, Total number of identified protein groups across DIA search strategies using two analysis tools (Spectronaut 16 and DIA-NN 1.8), each with match-between-runs enabled; chromatomes were searched either alone or together with the proteome datasets, and with additional DDA- and/or DIA-based spectral libraries. The final analysis used a hybrid (DDA + DIA library) strategy. Bars are shown for the chromatome and proteome of ePSC-H9, CS and CFS. **C**, Pairwise R² correlations of chromatomes and proteomes across ePSC-H9, CFS_1–3 and CS_1–3. **D**, PCA based on the proteomes of ePSCs and AxSCs; components 1 (78.6%) and 2 (10.4%) together describe ∼89% of the variance. **E**, Number of significant changes between the respective proteomes and chromatomes (Student’s t-test, p-value < 0.05, |log2 fold change| ≥ 1) for each pairwise comparison. **F**, UpSet plot of significant changes in relative chromatin binding (after protein-level normalization of the chromatome) for CFS vs ePSC-H9, CS vs ePSC-H9 and CS vs CFS, together with the unchanged set. **G**, Relative chromatin binding (log2 fold change) of selected marker proteins grouped as pluripotency markers (OCT4, DPPA4), shared chromatin factors (T/BRA), CFS markers (CDX2, HOXA3, HOXA5, HOXA11, HOXC6, HOXD3, HOXD4, HOXD9, HOXD10) and CS markers.

**Figure S3: A**, The top 50 differentially expressed genes in the scRNA-seq data of CFS_2, CS_2, and ePSCs-H9, as described in Figure 1F. Markers that provide additional insights regarding CFS cell lines include MLLT3, a hematopoietic regulator and downstream effector of T(BRA)^76^, SPRY4 a promoter of ESC stemness^77,78^, and SPRY2 which is associated to with both mesodermal and neural development.^79^ Markers that provide additional insights regarding CS cell lines include, PRTG, which is suggested to suppress of neural differentiation in early mouse embryos^80^, ZFHX4 which is upregulated during neural commitment^81^, ZFHX3 and ZIC2 both involved in the regulation of both muscle and neural development^82–85^ and CRABP1 which is a target of RA receptors. **B**, Schematics of the dual SMAD inhibition differentiation protocol of neural progenitor cells (NPCs). **C**, RT-qPCR characterization of NPCs using pluripotency and NPC markers. **D**, A dotplot of the single cell RNA-seq data of CS_2 (Figure 1F), and NPC (p3) lines displaying dorsoventral spinal cord markers and HOX genes. **E**, Derivation and characterization of CFS and CS cell lines from hiPSC line HMGU1.1-3 and hESC HUES6 with markers as in Figure 1C (gene expression normalized with the respective parental undifferentiated cells). **F**, Venn diagrams representing the total count of the genes expressed in >50% of the cells in human ES H9, HUES6, and iPS cell HMGU1 derived CFS (p15-18) and CS (p12-14) lines based on scRNA-seq (CFS H9: 3215, CFS HUES6: 2660, CFS HMGU1: 7059, CS H9: 6777, CS HUES6: 833, CS HMGU1: 8757 single cells). **G-H**, The expression of the markers categorized as per Figure 2A, B in the scRNA-seq data of human ES H9, HUES6, and iPS cell HMGU1 derived CFS and CS cell lines (Figure S3F). **I**, RT-qPCR analysis of CS cells derived from H9 and CFS cells for dorso-ventral neural tube and HOX genes (error bars represent SEM).

**Figure S4: A**, UMAP visualization of the integration of scRNA-seq data of human CFS_2 and CS_2 lines (Figure 1F) into the monkey gastrulation dataset^46^ spanning E20 to E29. **B**, Heatmaps displaying similarity of the human CFS and CS lines to the cell types in monkey^46^ and human^9^ gastrulation datasets. **C**, Violin plots displaying the expression of HOX genes in human CFS and CS cells, and the cells selected from monkey and human gastrulation datasets.

**Figure S5: A-C**, Dotplots showing the expression of undifferentiated CS markers in clusters from CFS and CS neural differentiation (Figure 3E). In the same manner, dotplots representing ventral spinal cord neuron markers in differentiated CS cells. **D**, Dotplots showcasing the (co)expression of marker genes corresponding to the cell clusters identified in Figure 3D, E. **E**, Dotplots displaying the expression of stage-specific markers of neural crest development in clusters from CFS and CS neural differentiation (del.: delamination, mes. trans.: mesenchymal transition, NC id.: neural crest identity).

**Figure S6: A**, Replications of the neural differentiation of additional CFS and CS cell lines. RT-qPCR analysis of *ISL1*, *MNX1* and *POU4F1* at differentiation Day28 (Figure 3A) showing H9 ES cell derived CFS lines 1-3 and CS lines 1 and 2 (Figure 1C) (error bars represent SEM). **B**, Confirmation of the sensory and motor neuron differentiation by immunostaining and imaging. Day28 cells were stained with BRN3A, MNX1, ISL1, and TUBB3 antibodies (scale bar: 50 µM). **C-D,** Dot plots showing HOX gene expression across clusters identified in day 14 and day 28 differentiated CFS cultures (Figure 3E), day 14 differentiated CS cultures (Figure 3E), and undifferentiated CFS and CS cultures (Figure 1F).

**Figure S7: A**, A summary of the peak K^+^ currents, cell input resistance, and resting membrane potential at each time point for measurements of 14-21 days CS, n=23; CFS, n=13; 49-56 days CS, n=11; CFS, n=16). **B**, Representative current- and voltage-clamp data illustrating the passive and active membrane properties of HMGU1 CFS and CS cells at the indicated times of differentiation. Displayed are voltage clamp recordings (left column), current clamp recordings of evoked action potentials (AP, middle column) and spontaneous excitatory postsynaptic currents (right column). Currents shown in the left column were evoked in response to increasingly depolarizing voltage (steps of 20mV; 300ms). The fast-inward Na^+^ current (downwards) was followed by slow outward K^+^ currents (upwards). Voltages shown in the middle column were in response to both hyperpolarizing (downward) current injection and depolarizing (upward) current injection (steps of 20 pA; 300ms), showing multiple evoked APs in response to continued stimulation. The current trace in the right column reveals spontaneous excitatory postsynaptic currents during CFS differentiation, indicating spontaneous neural network activity with functional synapses. **C**, A summary of peak Na+, peak K+ currents, and the total number of evoked APs at each time point for measurements of HMGU1 14-21 day CS cells, n=23; CFS cells, n=6; 49-56 days CFS cells, n=28. **D**, Immunofluorescence analysis of neuromuscular junction formation by α-bungarotoxin staining of day 21 neuronally differentiated CFS cells that were cocultured with C2C12 myoblasts for additional 12 days. **E,** Histological hematoxylin and eosin images showing a tumor mass displaying trilineage features of teratomas following injection of undifferentiated H9 ES cells in the kidney capsule of immunodeficient mice (scale bars: 50 µM).

**Figure S8: A,** STRING protein–protein interaction network of the 1,553 essential genes from the CFS essentialome that interact at a combined score ≥ 400 (16,301 edges); communities were detected with the Louvain algorithm (resolution 8.0). Node size is proportional to the absolute CRISPR essentiality score; nodes outlined in black are CFS-unique essential genes (n = 1,028) and nodes without an outline are also essential in hESCs (n = 984); colours denote gene-level functional meta-categories (epigenetic modifiers, transcription factors, OXPHOS, metabolic enzymes and the indicated subcellular/complex categories). Of the 2,012 essentialome genes, 1,552 fell into communities of ≥ 3 connected proteins. CDX2 was additionally highlighted. **B**, Zoom of the epigenetic-modifier and transcription-factor sub-network (e.g. PRC1/PRC2, TrxG, CHD family, NuRD, SWI/SNF (BAF), HATs, HDACs, H3K9 writers/readers/erasers, mCpG readers, NMP/axis TFs, HOX genes, DNMT1, MYCN), with the dominant complex or enrichment term labelled. **C**, Zoom of the metabolic and OXPHOS sub-network (glycolysis, OXPHOS electron-transport chain and their regulators TFAM, PPARGC1B, GABPA/B, NRF1 and YY1; AMPK signalling). **D**, Boxplots of relative intensities (log2) of the identified histone H3 PTMs across ePSC-H9, CS and CFS; each replicate is plotted as an individual point (quadruplicates per line). **E**, As in (**D**) for the histone H4 PTMs. **F**, Heatmap of the log2 fold changes of the histone H3 PTMs in ePSC-H9, CFS and CS. **G**, As in (**F**) for the histone H4 PTMs.

**Figure S9: A**, Experimental design: human ePSC-H9 received a 24h CHIR pulse and were then independently expanded as CFS and CS over ten passages (p1–p10); matched samples for chromatome and RRBS were collected at the starting culture (H9), after 24h and at every passage. **B**, Identified chromatome protein groups per sample (DIA-NN, non-imputed; range ∼7,854– 9,127), colored by sample identity (ePSC-H9/24h, CFS, CS); CFS p5 and CS p6 were excluded from downstream analyses. **C**, PCA of the 18 batch-corrected, KNN-imputed, row Z-scored chromatome samples, PC1 (39.1%) vs PC2 (12.8%). **D**, PC1 (39.1%) vs PC3 (7.7%). **E**, Row Z-scored chromatome abundance of differentiation markers across the 18 samples (columns in experimental order; ePSC/24h, CFS p1–p10, CS p1–p10; rows hierarchically clustered), color-coded by category (pluripotency, stemness, CFS, CS). **F**, Row Z-scored chromatin abundance of the H3K27 demethylases KDM6A and KDM6B. **G**, Row Z-scored chromatin abundance of the DNA-methylation machinery: DNMTs (DNMT1, DNMT3A, DNMT3B), recruiters/readers (UHRF1, PCLAF) and TET demethylases (TET1, TET2, TET3). **H**, Heatmap of motif enrichment in differentially methylated regions (DMRs) of CFS passage p10 (compared to H9), with DMRs grouped according to demethylation size. The dark green bin holds the most hypermethylated DMRs, the dark brown bin the most hypomethylated DMRs. Shown are motifs with significant enrichment (p < 0.01) in at least one bin. The heatmap indicates log2 enrichment in each bin compared to all other bins (one-sided Fisher’s exact test with Benjamini-Hochberg correction). **I**, Motif-enrichment trajectory for the CDX motifs (CDX1, CDX2, CDX4) across passages. Each point shows enrichment in the respective passage compared to H9. **J**, Enrichment of HOX motifs in hypomethylated regions across passages for CFS. Each point shows enrichment in the respective passage compared to H9. Included were HOX proteins whose motifs were found significantly enriched (p < 1e-3) in the hypomethylated (compared to H9) regions of passage p10. **K**, Correlation of HOX/CDX motif occurrence in DMRs of CFS passage p10 (compared to H9) shown as a heatmap. Strong correlation between motifs indicates that their enrichment probably stems from one underlying motif, making them potential false positives. Included were HOX and CDX proteins whose motifs were found significantly enriched (p < 1e-3) in the hypomethylated (compared to H9) regions of passage p10. Derived from RRBS. **L**, Motif-enrichment trajectories for transcription-factor motifs in hypermethylated regions across passages for CFS. Each point shows enrichment in the respective passage compared to H9.

**Figure S10: A-B**, Screening included small molecules and growth factors pertaining to the derivation of mouse CFS and CS derivation using human CFS and CS condition with the addition of FGF8. Green rectangles represent successful derivations of cell lines cultured with cell splitting of >10 passages, labeled 1 through 5. **C**, Representative immunostaining images of mCFS_1 (p21) for Sox2, Brachyury (T ortholog), and Cdx2 (scale bar: 20 µm). **D**, RT-qPCR analysis of mouse CFS lines for pluripotency markers, NMP (including early NMP marker Nkx1-2 and late NMP marker Mnx1) and paraxial mesoderm development associated genes corresponding to mCFS_2-5 lines related to Figure 7A-B (relative expression normalized to parental cells (n=4). **E**, A transverse section of embryo 2 (Figure 7F) showing DiI-labelled cells in the neural tube (NT) and somite (som). Immunostaining images of Sox2 (green) and Meox1 (red) are shown below. Labeled cells correctly express Sox2 and Meox1 in NT and somites respectively. **F**, Representative images showing the transplantation site of mouse CFS cells labeled with DiO at E8.5 (i) and incorporation into presomitic mesoderm (PSM), som, NT, chordo-neural hinge (CNH) and tailbud mesoderm (TBM) at E9.5 (ii). Embryos sectioned at the indicated site at E10.5 (iii), and immunostained for Sox2 (red) and Hoechst (blue). Cells arranged in auxiliary neural tube structures were derived from mouse CFS cells (green: DiO labeled) (iv-xii).

**Figure S11: A,** A heatmap displaying similarity of the three mouse CFS lines to the cell types in mouse gastrulation dataset. **B**, Violin plots displaying expression of HOX genes in mouse CFS cells and the cells selected from mouse gastrulation dataset. **C,** Glycolysis and oxidative phosphorylation scores computed based on the expression of genes associated with the respective metabolic pathways^75^ in the scRNA sequencing of mouse CFS lines (Figure 8A).

## KEY RESOURCES TABLE

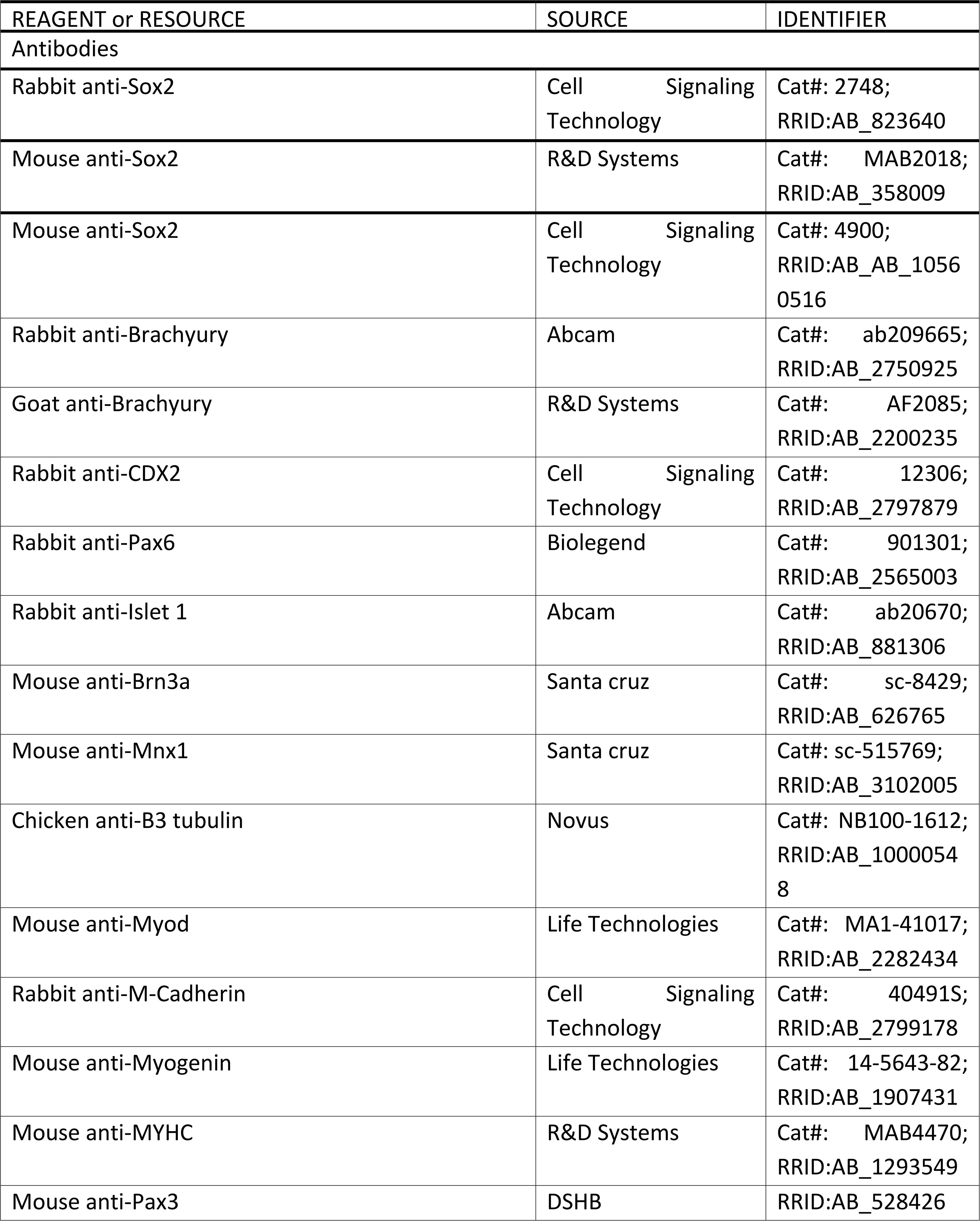
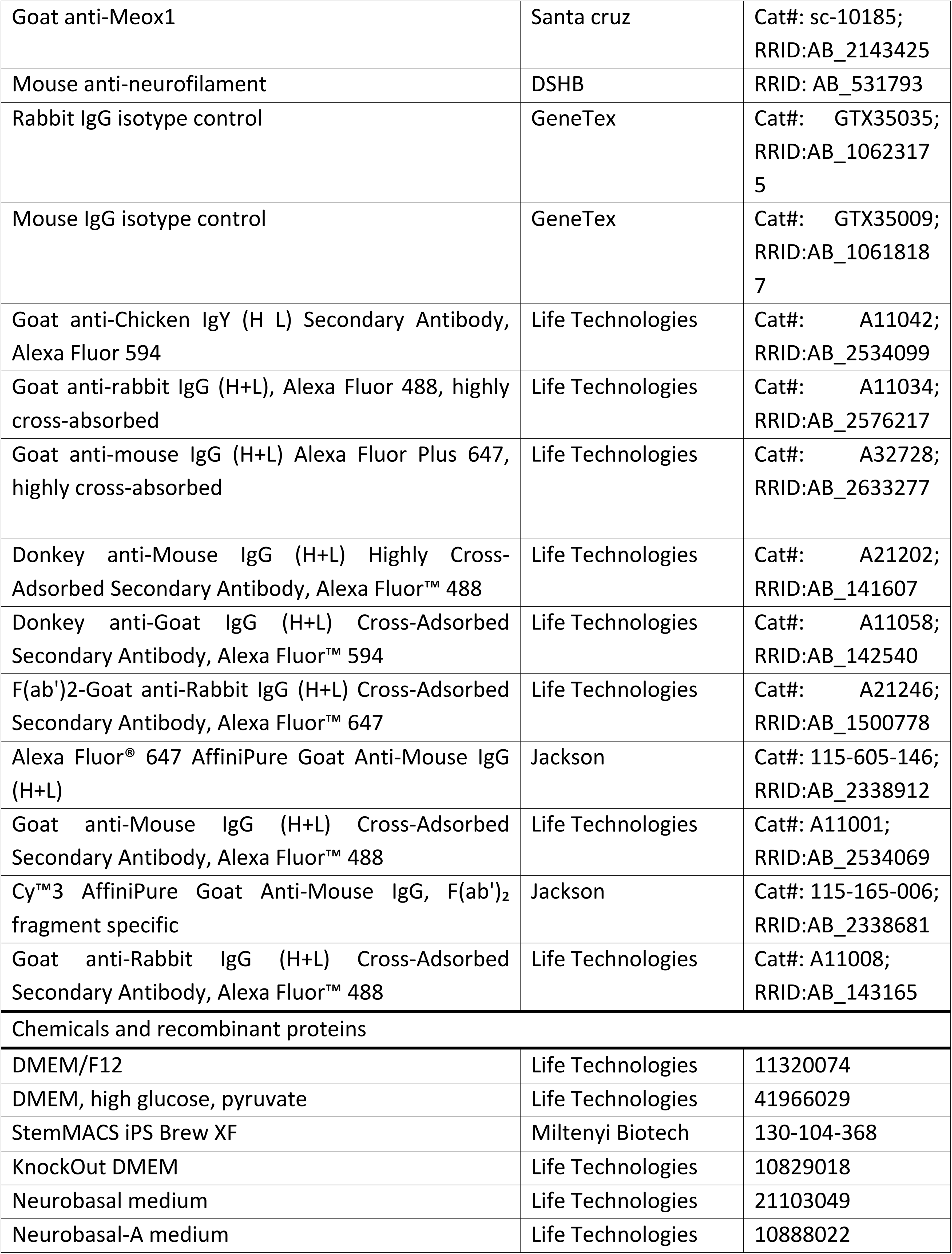
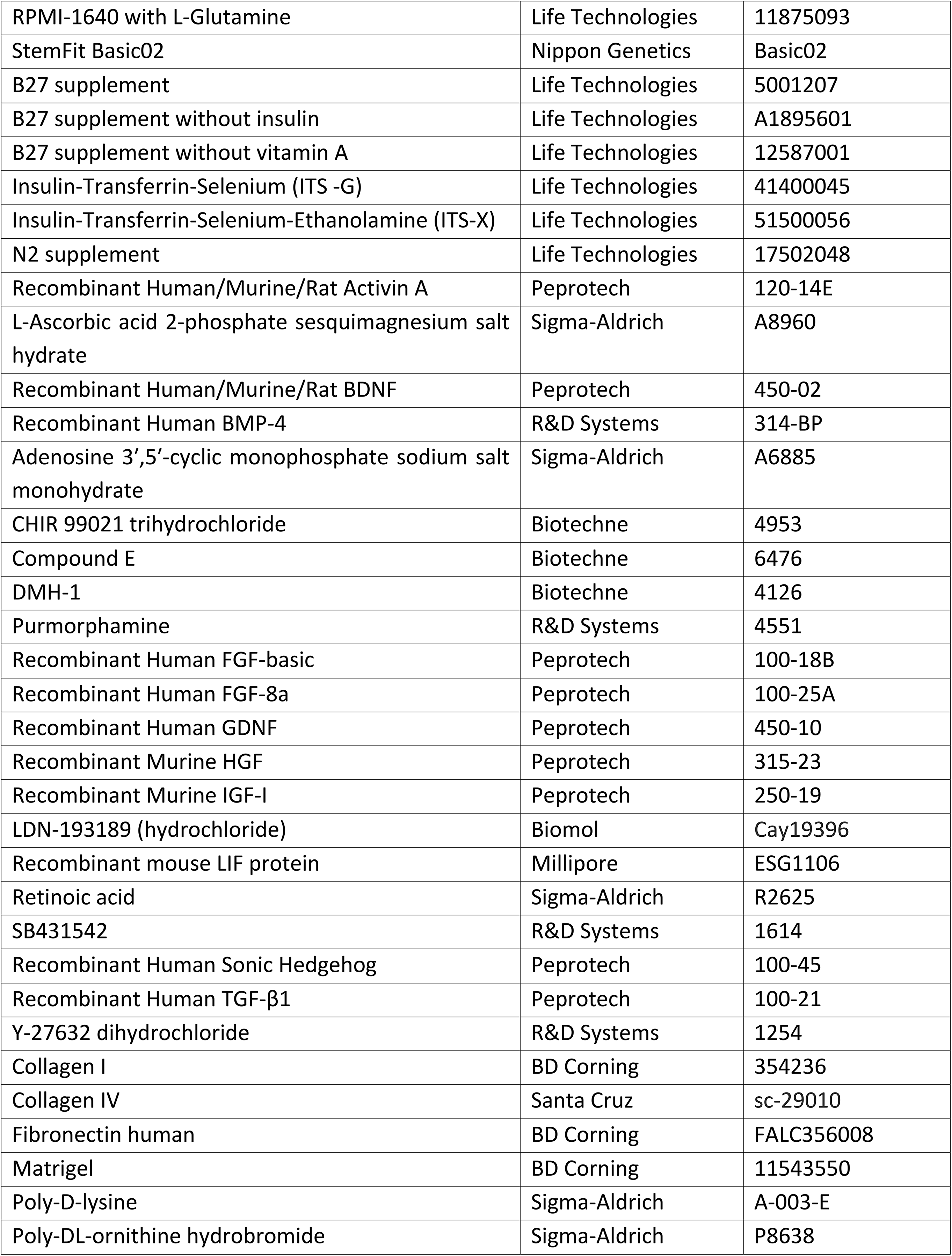
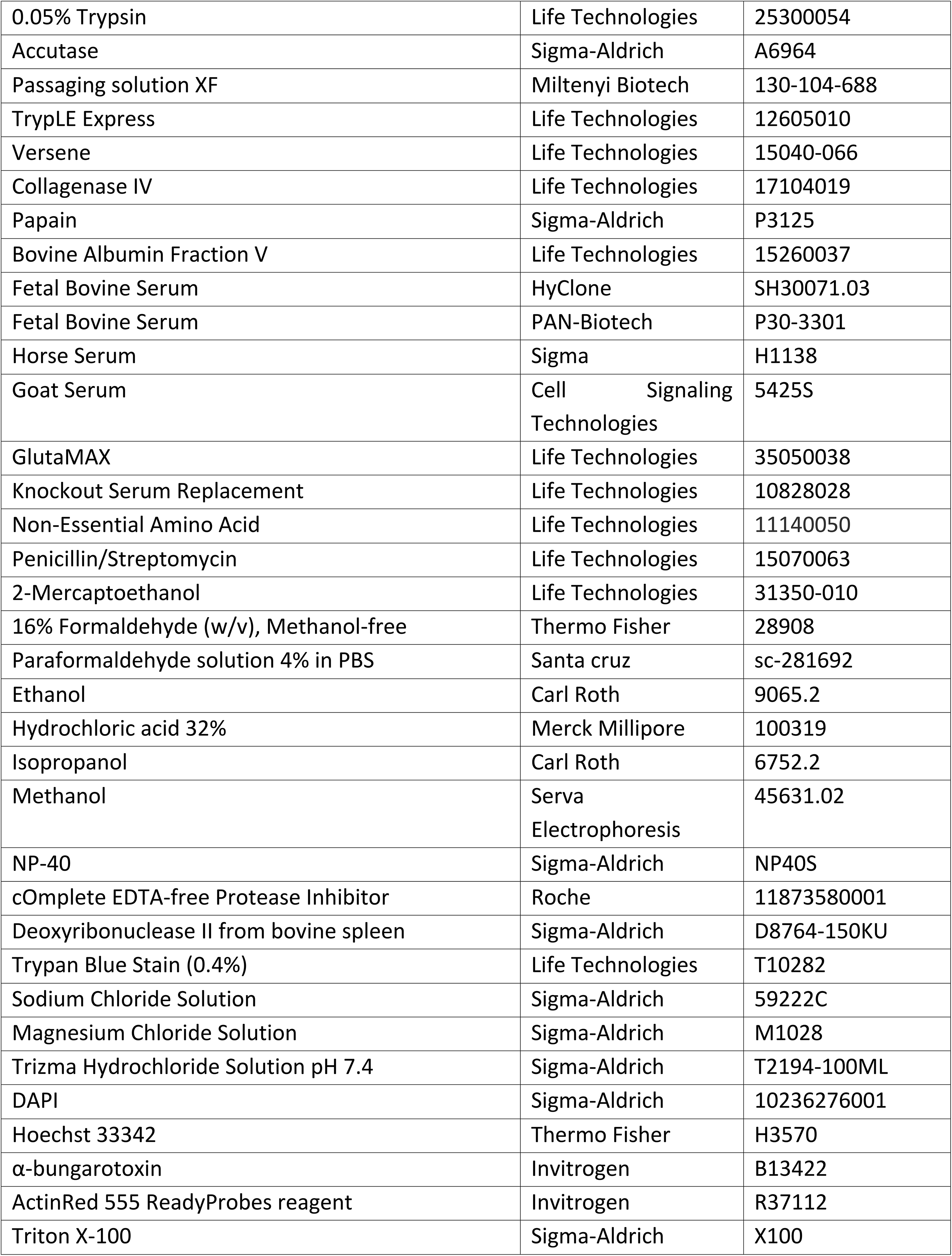
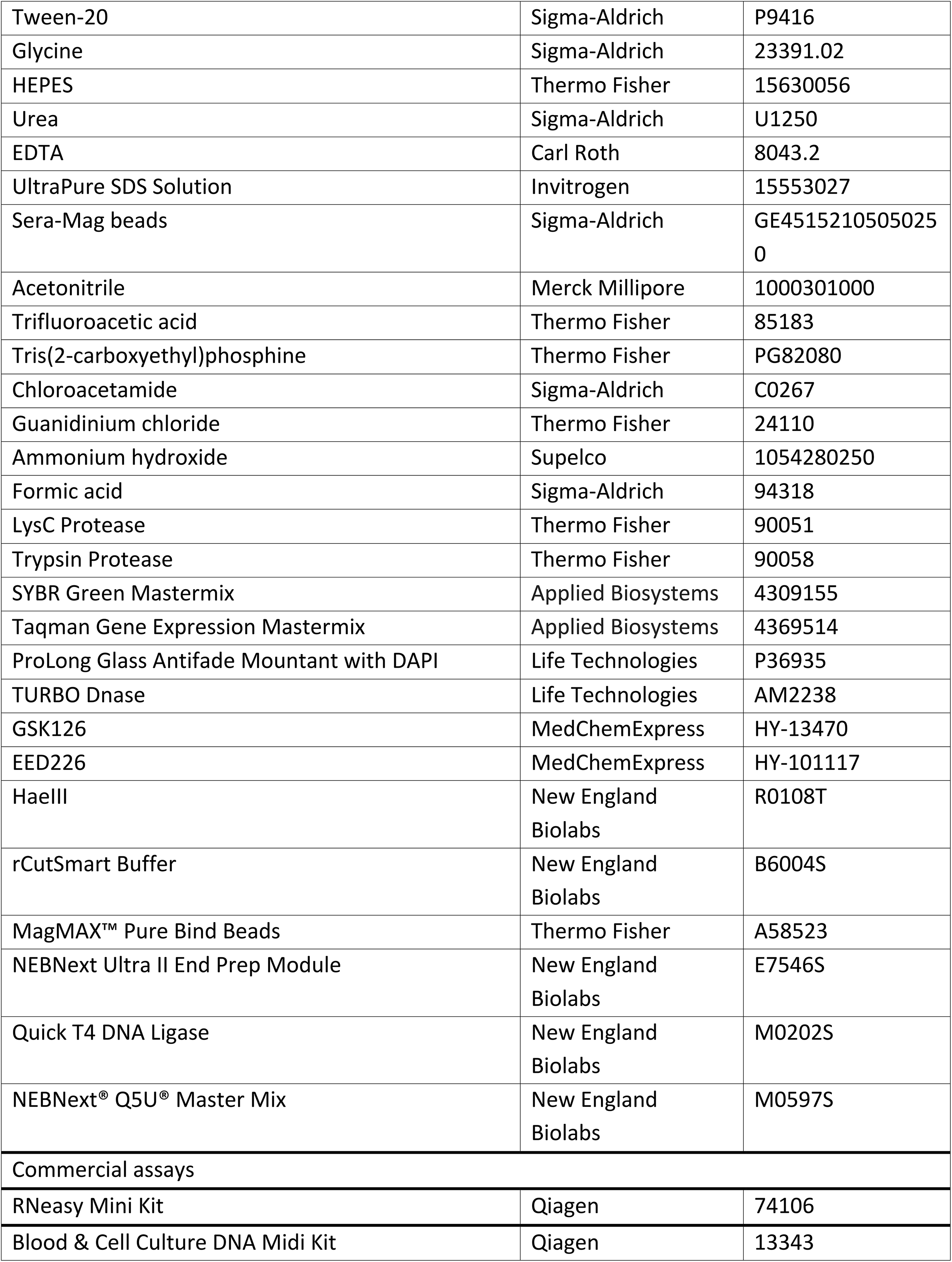
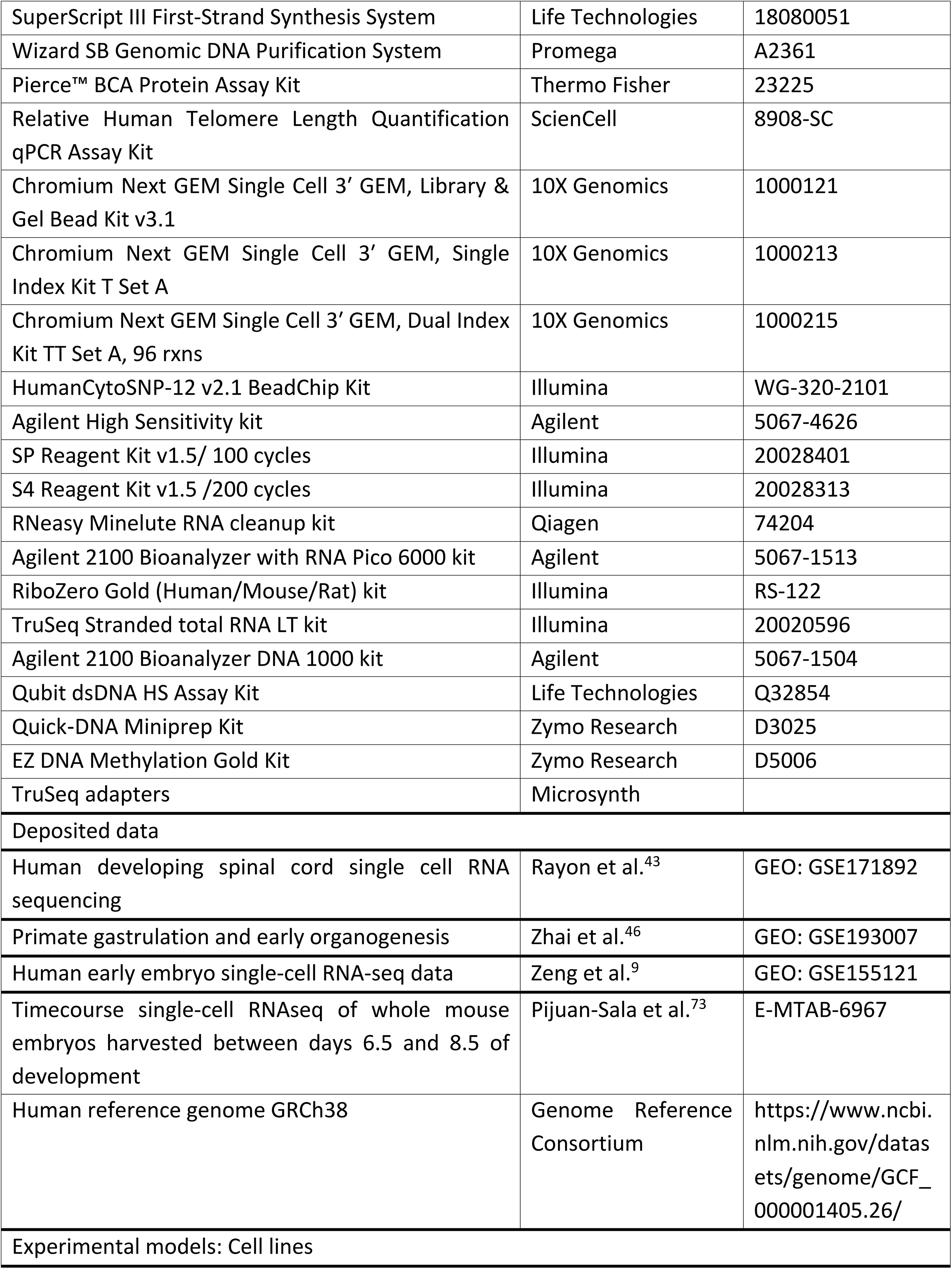
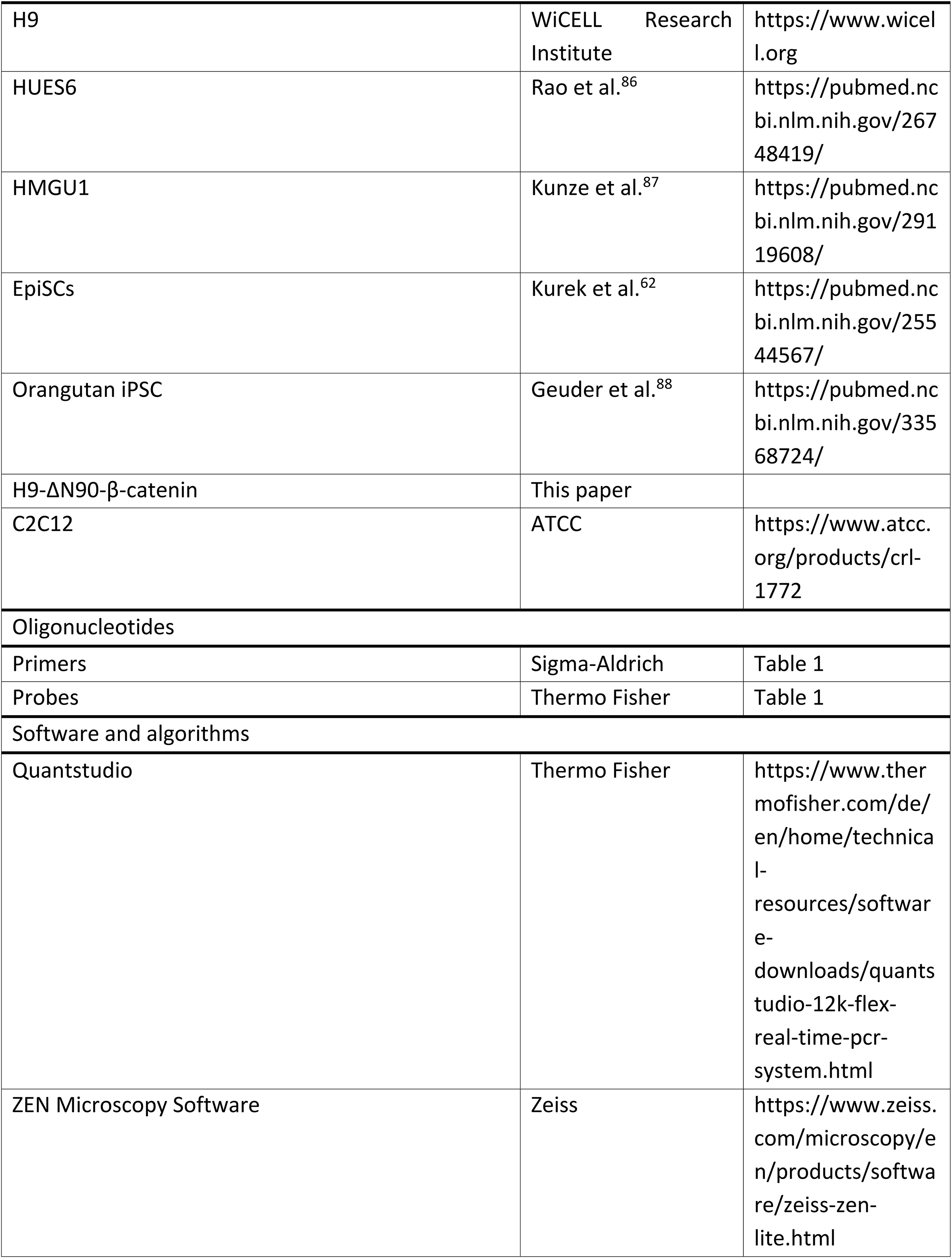
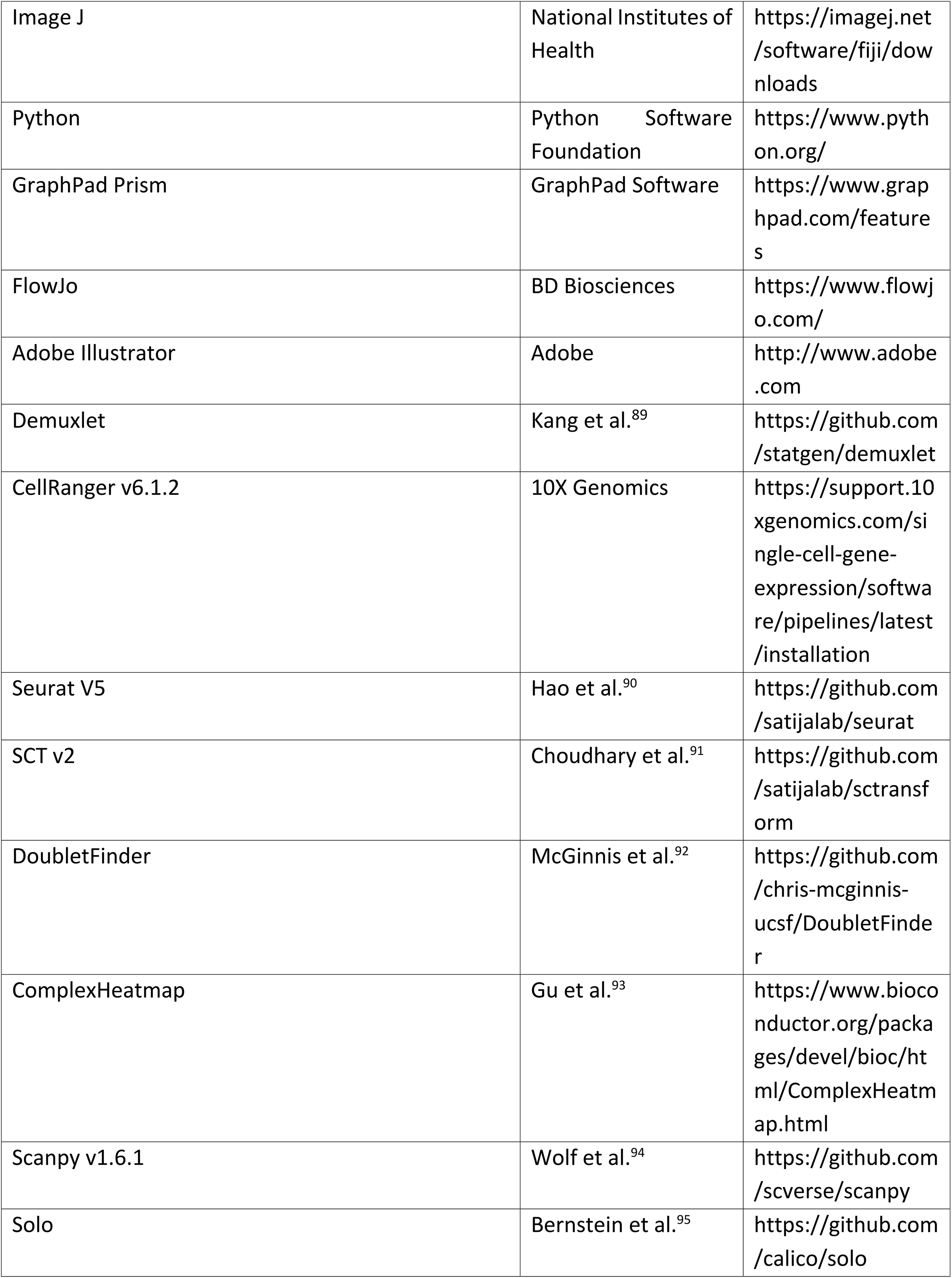
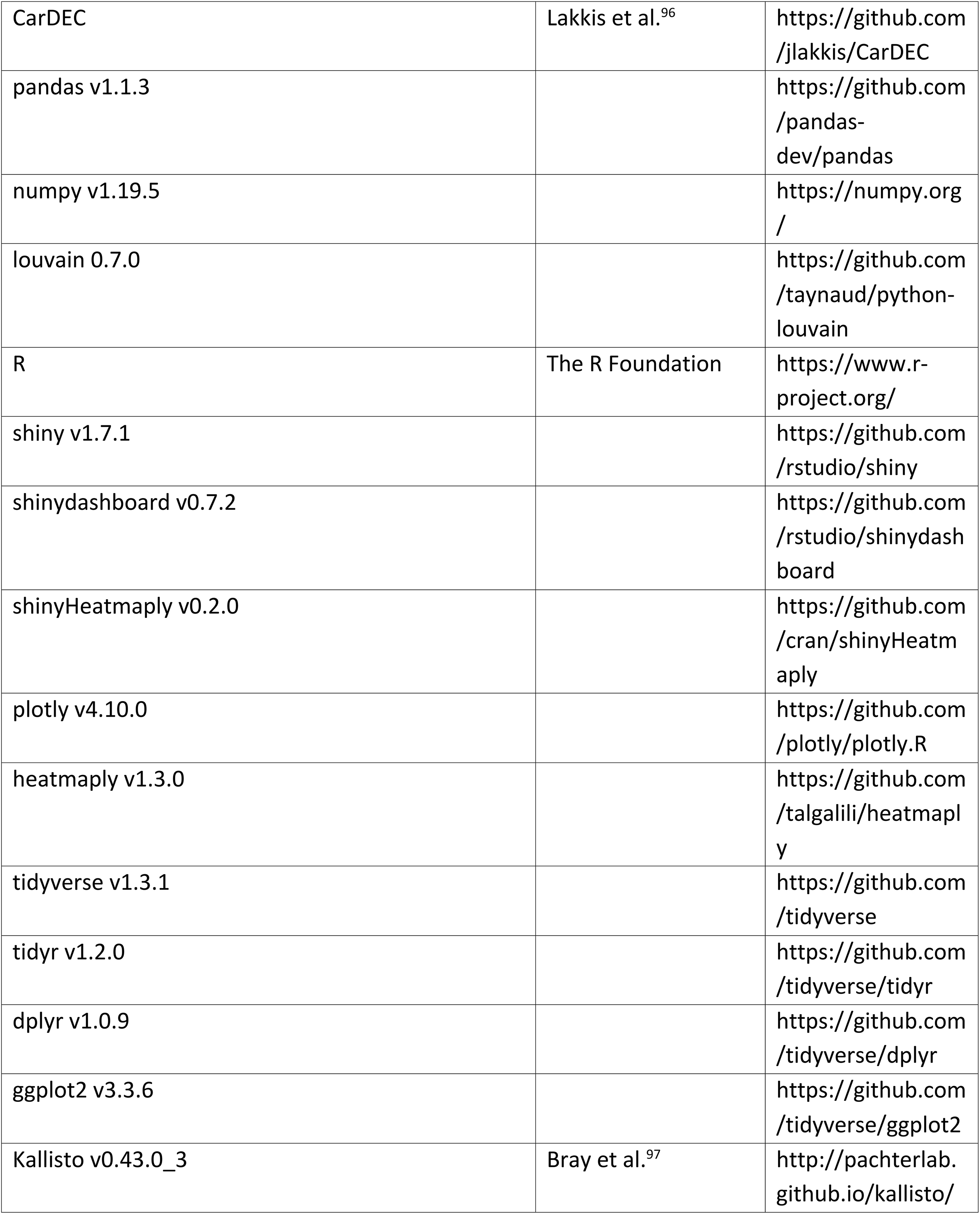
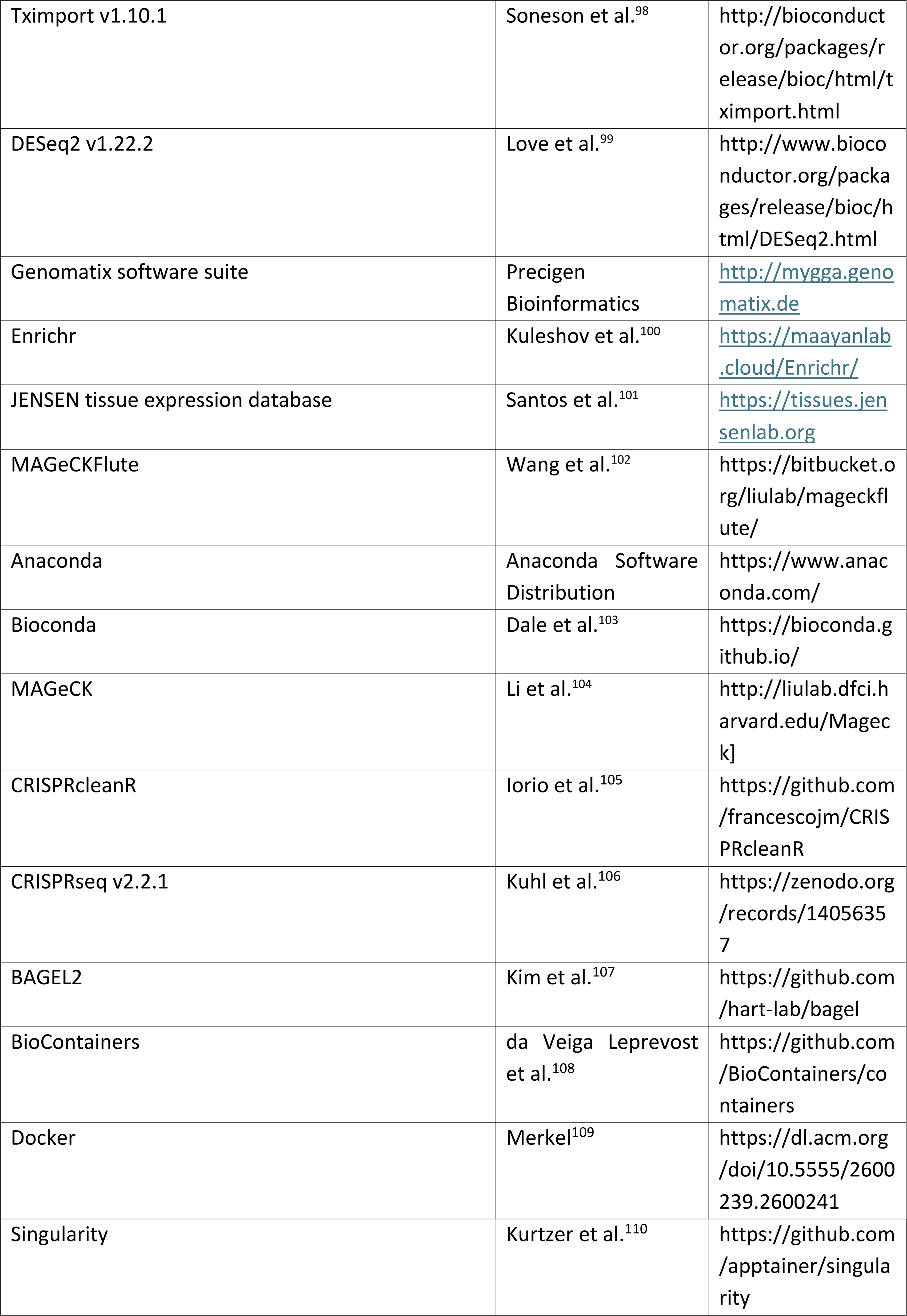

## AUTHOR CONTRIBUTIONS

D.K., E.U., E.R., D.S., M.Dr. designed the study and M.Dr. supervised the study. D.K. established human AxSCs lines; performed screening for mouse and orangutan AxSCs culture conditions and differentiation, immunocytochemistry, RT-qPCR experiments; prepared the proteome, chromatome, essentialome and scRNA-sequencing samples and prepared the figures. E.U. prepared, measured and analyzed the proteome and chromatome samples under supervision of H.L. and M.M, prepared the related figures and programmed the web application. E.R. established additional human AxSC lines, assisted with cell line establishment and differentiation experiments, experimental design and analysis of scRNA-sequencing samples. D.S. and A.H.A performed screening for human AxSCs, established the human HMGU1 AxSCs, generated the β-catenin hESC line and performed RNA-sequencing. A.L. and V.W. performed *in vivo* injection and analysis of the mouse AxSCs and contributed to the writing of the manuscript. A.B.C. and R.W. established the CFS cell line from a human embryo and performed the RT-qPCR experiment. S.H. and M.Da. performed electrophysiology experiments under supervision of P.K. A.Y. conducted lentiviral delivery for essential gene analysis, A.Bo. processed the sequencing data, and A.M.A. participated in data analysis. K.W. performed the time-course reduced representation bisulfite sequencing experiments and data analysis under the supervision of P.G with E.U. X.Y. performed integration of scRNA-sequencing datasets with publicly available gastrulation datasets under supervision of Z.X. and H.W. M.Dr. perfomed teratoma assays and A.B and J.B. performed histopathological analysis. M.S. generated libraries for scRNA-sequencing under supervision of H.L. K.A. performed preprocessing of the scRNA-sequencing samples and assisted with downstream analysis. B.T. generated the time-lapse videos and performed with B.S. analysis of neuromuscular junction. N.S. performed the SLC25A1 inhibitor assays. C.S. advised on manuscript structure and revisions. R.d.V. and M.H. performed Seahorse experiment. Y.W. performed flow cytometry. D.K. and M.Dr. wrote the manuscript.

## ACKNOWLEDGEMENTS

This work was partially funded by the Lingling Wiyadharma Fonds ter bevordering van de beoefening der Natuurwetenschappen, Stichting Painless, the John Templeton foundation grant 62572 and VolkswagenStiftung under grant A130140. We would like to thank the German Academic Exchange Service (DAAD) for supporting DK. We acknowledge the technical support of Core Facility Genomics at Helmholtz Munich. We thank Dr. Inti Alberto De la Rosa-Velázquez for help with sequencing, Dr. Peter Lichtner for help with genotyping, Dr. Thomas Walzthoeni and Xavier Pastor Hostench for help with data processing, Moritz Thomas and Dr. Carsten Marr for help with data analysis. We would like to thank Dr. Thure Adler, Dr. Anna Pertek and Dr. Friederike Matheus for help with the animal experiments as well as Dr. Silvia Schirge for help with embryo dissections. We would like to acknowledge Core Facility Statistical Consulting at Helmholtz Munich for their guidance on statistical analysis, particularly Ronan Le Gleut and Dr. Kainat Khowaja. We thank CRC 1309 (ID: 325871075) and the BMBF Cluster4Future program (Cluster for Nucleic Acid Therapeutics Munich, CNATM, ID: 03ZU1201AA). We are very grateful to individuals for donating tissue. We thank Richard Anderson and Norma A. Forson for obtaining consent sourcing and providing human samples.

## DATA AVAILABILITY

Proteome and chromatome data have been deposited in ProteomeXchange with identifier PXD046087, can also be explored at the following integrative website https://axialstemcells.shinyapps.io/atlas/. Single cell RNA sequencing data is available on NCBI Gene Expression Omnibus under accession number GSE247215. The publicly available bulk RNA sequencing dataset on NCBI Gene Expression Omnibus with accession number GSE130381 was used in this study.

## METHOD DETAILS

### Maintenance of pluripotent stem cells

All cell culture of all species was maintained at 37°C with 5% CO2 and 5% O2 on Matrigel (1:100, BD Corning) coated plates unless otherwise specified. After passaging cells, 10 µM Y-27632 (R&D Systems) was always added to the medium for the first 24 hours. Medium was refreshed daily, unless otherwise specified. All cell culture components were purchased from Life Technologies unless specified otherwise.

Human pluripotent stem cells (hPSCs), including H9 (WiCell Research Institute), HUES6^86^, and HMGU#1^87^, were cultured using StemMACS iPS-Brew XF medium (Miltenyi Biotec). Cells were split using StemMACS Passaging Solution XF (Miltenyi Biotec) at a ratio of 1:10. Experiments involving human stem cells were performed in accordance with permissions obtained from the Robert-Koch-Institute (AZ: 3.04.02/0151 to MD to and AZ 3.04.02/0172 to CS).

E14 mouse embryonic stem cells (mESCs)-based epiblast stem cells (EpiSCs)^62^ were cultured in 1:1 DMEM/F12 and Neurobasal medium, supplemented with 0.5X N2, 0.5X B27, 0.033% BSA, 50 µM β-mercaptoethanol, 1X GlutaMAX, 1X penicillin/streptomycin, 20 ng/ml Activin A (Peprotech), 12 ng/ml FGF2 (Peprotech), and 2 µM IWP2 (Santa Cruz). The cells were split using 1X TrypLE Express at a ratio of 1:20-1:40.

Pongo abelii Sumatran orangutan induced pluripotent stem cells (iPSCs)^88^ were cultured in StemFit Basic02 medium (Nippon Genetics) supplemented with 1X penicillin/streptomycin and 100 ng/ml FGF2. The cells were regularly split using Versene at a ratio of 1:10.

### Maintenance of C2C12 myoblast cells

C2C12 myoblast cells (ATCC) were cultured in DMEM with 4,5 g/L D-glucose, 4 mM L-glutamine and 1mM pyruvate (Thermo Fisher) containing 10% fetal bovine serum (FBS) and 0.5% penicillin/streptomycin. Cells were cultured 37°C with 5% CO2. Medium was replenished every 2 days. C2C12 cells were passaged to uncoated culture dishes when cells reach 70-80% confluency using 1X TrypLE Express. Myoblasts differentiation was induced at 90% confluency by changing the medium to DMEM supplemented with 2% horse serum (HS) and 0.5% penicillin/streptomycin. Differentiation medium was refreshed every other day.

### Generation ΔN90 β-catenin hESC line

To generate tetracycline-inducible overexpression of constitutively active β-catenin (with a deletion of the first 90 amino acids from the N-terminus), H9 cells were transfected with the PB-GFP-P2A-ΔNβCAT plasmid and a Piggybac transposase-coding plasmid using the P3 primary cell 4D nucleofector kit (Lonza). After 48 hours, the cells were transferred to a 10-cm dish and subjected to selection with 50 µg/ml Hygromycin B for a duration of 2 weeks. Subsequently, the polyclonal stable line was maintained following the same protocol as the parental H9 line, but with the addition of 25 µg/ml Hygromycin B.

### Induction of WNT signaling and time course analysis

Cells were split using Accutase (Sigma) at a density of 2.5×10^5^ cells per well in mTesR1 medium. After 24 hours, the medium was replaced with differentiation medium composed of RPMI-1640 with L-Glutamine and 1X B27 without insulin, supplemented with 10 µM CHIR99021 (Tocris). For beta-catenin overexpression the CHIR99021 was substituted with 1 µg/ml doxycycline (Clontech).

### Derivation of human axial stem cells

H9 and HUES6 PSCs were adapted to single cell splitting through passaging with TrypLE Express at 37°C for 6 minutes, for a minimum of 3 passages. Subsequently, PSCs (between passage 55-60) were plated at a density of 71.5×10^3^ cells/cm^2^ in StemMACS iPS-Brew XF. After 24 hours the cells were washed with PBS and incubated for 24 hours in RPMI-1640 supplemented with 1X B27 without insulin and 10 µM CHIR99021. Next, the cells were dissociated using TrypLE Express for 5 minutes at 37°C and centrifuged at 300 g for 3 minutes. The pellet was resuspended either in human CFS medium, (RPMI-1640, 1X NEAA, 1X B27 without vitamin A, 100 ng/ml FGF2, 5 µM CHIR99021, and 10 µM SB43152), or in human CS medium (RPMI-1640, 1X NEAA, 1X B27 without vitamin A, 5 µM CHIR99021, and 10 µM SB43152). For the first 7 passages, the cells were split twice a week at a ratio between 1:5-1:20 using TrypLE Express for 4-5 minutes for CFS and 4 minutes for CS at 37°C. After passage 7, both cell lines were split at a ratio between 1:10-1:20, with CS cells being split once a week and CFS cells being split twice a week, on average.

HMGU1 AxSCs were derived using a similar method with minor differences. The cells were split at a ratio of 1:20 - 1:30 in either human CFS or CS maintenance medium. Routine splitting was performed at a 1:10-1:20 dilution using either StemMACS Passaging Solution XF or Versene.

### Derivation of mouse axial stem cells

EpiSCs were split using TrypLE Express and plated at a dilution of 1:20-1:40 directly in mouse CFS medium (RPMI-1640, 1X NEAA, 1X B27 without vitamin A, 50 ng/ml FGF2, 3 µM CHIR99021, and 10 µM SB43152). The cells were continuously maintained on either a Matrigel-coated plate (1:100) or on mitotically inactivated fibroblast feeders seeded at a density of 1×10^5^ cells/cm^2^ until passage 10. After that, they were maintained on Matrigel-coated plates (1:100). The cells were split two or three times per week by incubating them with TrypLE Express at 37°C for 4 minutes and then seeded at a dilution of 1:20-1:40.

Stem zone regions of Foxa2VenusFusion x mTmG litters were dissected at E8.5. Mice were kept and experiments performed at the central facilities at Helmholtz Munich German Research Center of Environmental Health in accordance with the German animal welfare legislation and acknowledged guidelines of the Society of Laboratory Animals (GV-SOLAS) and of the Federation of Laboratory Animal Science Associations (FELASA). Mice are kept under SPF conditions in animal rooms with light cycle of 12/12 hours, temperature of 20-24°C and humidity of 45-65%. Mice receive sterile filtered water and standard diet for rodents ad libitum. For embryo generation mice at the age of ≥ 8 weeks were used. Embryonic tissues were centrifuged at 400 g for 4 minutes. The supernatant was discarded, and the pellet was dissolved in 0.25% Trypsin, then incubated at 37°C for 15 minutes. After incubation, FBS was added, followed by centrifugation at 400 g for 4 minutes. The pellet was then resuspended in mouse CFS medium and seeded on Matrigel-coated plates. For the first three passages 1X penicillin/streptomycin was added to the medium. The cells were split twice a week by incubating them with TrypLE Express at 37°C for 4 minutes, and then seeded at a dilution of 1:20-1:40. After passage 7, the cells were plated on feeder cells or Matrigel-coated plates.

### Derivation of orangutan axial stem cells

Sumatran orangutan iPSCs were treated for 24 hours with an induction medium comprising RPMI-1640, 1X B27 without insulin, and either 5 or 10 µM CHIR99021. Next, the cells were dissociated using StemMACS Passaging Solution XF and resuspended in human CS medium as described earlier. The cells were split at a dilution of 1:10-1:20 once or twice a week, using StemMACS Passaging Solution XF.

### Derivation of human neural progenitor cells

H9 cells were rinsed with PBS twice and incubated with Collagenase IV (2 mg/ml) for 30 minutes at 37°C followed by plating 1:1 onto low-attachment 6-well plate in a medium consisting of DMEM/F12, 20% KSR, 1% GlutaMAX, 1% NEAA, 1 μM DMH-1, 10 μM SB431542, 3 μM CHIR99021, 0.5 μM Purmorphamine (PMA), 10 μM Y-27632. The next 2 days, fresh medium was applied without Y-27632. On Day3 and Day4, the medium was replaced with 1:1 DMEM/F12 and Neurobasal-A medium, 1:100 B27 without vitamin A, 1:200 N2, 1% GlutaMAX, 1 μM DMH-1, 10 μM SB431542, 3 μM CHIR99021, 0.5 μM PMA. On Day5, the medium was changed to 1:1 DMEM/F12 and Neurobasal-A medium, 1:100 B27 without vitamin A, 1:200 N2, 1% GlutaMAX, 1 μM DMH-1, 50 μg/ml Ascorbic acid (AA). On Day6, the Day5 medium with 5 ng/ml FGF2 was added. On Day7, cells were plated on a Matrigel-coated (1:100) 6-well plate at 1:1 ratio in a medium consisting of 1:1 DMEM/F12 and Neurobasal-A medium, 1:100 B27 without vitamin A, 1:200 N2, 1% GlutaMAX, 1 μM DMH-1, 50 μg/ml AA, 20 ng/ml FGF2 (NPC medium). NPC medium was changed daily until Day14 and then every other day. On Day14 and Day21, cells were washed with PBS and treated with Collagenase IV (2mg/ml) at 37°C for 15 minutes and TrypLE Express in 37°C for 4 minutes respectively. Cells were plated on Matrigel-coated (1:100) plates at a ratio between 1:4-1:6. After Day21, NPCs were passaged using TrypLE Express when the confluency reached >70%. Cells were maintained at 37°C, 5% CO2 and 5% O2.

### Processing of human embryonic tissue and derivation of human CFS axial stem cells

Fresh human embryos were obtained immediately after elective termination of pregnancy either from the Royal Infirmary of Edinburgh (RAA/FT study, approved by the Lothian Research Ethics Committee, Reference: 08/S1101/1). All patients donating tissue had given informed written consent for the use of human embryonic tissues in research prior to tissues being obtained and anonymized. The developmental stage of human embryos was determined by the CS system ^111,112^.

Dissections were performed under a tissue culture hood using a low power stereomicroscope (Leica MZ8) in sterile dissection buffer (Dulbecco’s phosphate buffered saline (PBS) with Mg2+ and Ca2+ ions (Sigma-Aldrich) containing 7% fetal calf serum (FCS) (Thermo Fisher Scientific or APS), 100 IU/mL penicillin and 100 μg/mL streptomycin (Invitrogen)). Further sub-dissection of the tail bud region is described in the Results section. Images of human tissue were taken using a Leica MZ FLIII with DFC7000T camera (Leica). The tail bud tissue was dissociated to small clumps in 35mm tissue culture treated dishes. For this, a drop of 800 μL of TrypLE was used and incubated at RT for 10 min with gentle pipetting to dissociate the tissue. The dissociation was stopped by pipetting the tissue/cell clumps into human CFS media 2 times before platting into standard human CFS culture conditions (this paper). Growing colonies were monitored daily and a few were manually picked for expansion before characterisation.

### Neural differentiation from human axial stem cells

Plates were coated with 20 µg/ml Poly-D-lysine (Sigma) and 20 µg/ml Poly-ornithine (Sigma) in PBS and incubated overnight at 37°C. The following day, the coating solution was replaced with 20 µg/ml Collagen I (BD Corning) and 20 µg/ml Fibronectin (BD Corning) in PBS and incubated overnight at 37°C. The next day, the coating solution was substituted with 10 µg/ml Collagen IV (Santa cruz) in 0.05M HCl (Merck) and incubated for 2-4 hours at 37°C, followed by 2 washes with PBS. For immunostaining experiments, cells were directly differentiated on either coverslips or chamber slides coated as described above.

Human CFS and CS lines were dissociated using TrypLE Express and plated on the pre-coated plates in the respective medium, at a density of 28.5×10^3^ cells/cm^2^ for CFS and 71.5×10^3^ cells/cm^2^ for CS. The next day, the medium was replaced with differentiation medium consisting of a 1:1 mixture of DMEM/F12 and Neurobasal-A medium, 1X B27, 1X N2, 0.1 µM Retinoic acid (RA) (Sigma), 100 ng/ml SHH (Peprotech), 100 µM cAMP (Sigma), 10 ng/ml GDNF (Peprotech), 10 ng/ml BDNF (Peprotech), 10 ng/ml IGF (Peprotech), and 0.1 µM Compound E (Biotechne). The medium was changed every other day.

For myotube-neuronal co-culture, 1.25×10^4^ cells/cm2 of CFS cells were plated on Matrigel coated dishes in CFS medium. After 24h, medium was changed to neuronal differentiation medium. Differentiated neuron clusters were harvested after 18 days of differentiation by 10 min incubation with TrypLE Express. Cells were replated on top of three-day old myotubes, which were differentiated from myoblasts separately. Myoblasts were cultured in DMEM with 10% FBS. Cells were grown until 80-90% confluency after which medium was changed to DMEM with 2% HS. After replating the neural cells on top of the myotubes medium was changed to co-culture medium consisting of DMEM with 2% HS, 10 ng/ml BDNF, 10 ng/ml GDNF.

### Skeletal muscle differentiation from human axial stem cells

Human CFS and CS lines were dissociated using TrypLE Express and seeded on Matrigel (incubated for >4 hours). The cells were seeded in their respective medium at a density between 1.5-2.5×10^4^ cells/cm^2^. The following day, the medium was changed to DMEM/F12 supplemented with 1X NEAA, 1X ITS-G, 3 µM CHIR99021, 0.5 µM LDN193189 (Biomol), 5 ng/ml FGF2, and 10 nM RA. On Day 2, FGF2 and RA were replaced with 34 ng/ml SHH. On Day 6, the medium was changed to DMEM/F12 with 1X NEAA, 1X ITS-G, 10 ng/ml FGF2, and 10 ng/ml HGF. On Day 10, the cells were split by washing with PBS and incubating with TrypLE Express for 6-7 minutes at 37°C. The cells were then harvested, centrifuged at 400 g for 4 minutes, and the supernatant was removed. The pellet was resuspended in DMEM/F12 with 1X NEAA, 1X ITS-G, 10 ng/ml FGF2, and 10 ng/ml HGF (Peprotech) and plated at a density of 2-2.8×10^4^ cells/cm^2^. From Day 12 onwards, the medium was changed to DMEM/F12 with 1X NEAA, 1X ITS-X, and 10 ng/ml HGF. Fresh medium was applied every other day during the differentiation process.

### Quantitative RT-PCR

RNA extraction was conducted using the RNeasy Mini Kit from Qiagen following the manufacturer’s instructions. The extracted RNA samples were then reverse-transcribed using the SuperScript III First-Strand Synthesis System (Life Technologies), according to the manufacturer’s instructions. RT-qPCR was performed in 384-well plates using either the Power SYBR Green PCR Mastermix or the Taqman Gene Expression Assay Mastermix (Applied Biosystems), with a total volume of 10 µl. The reaction was carried out under the following conditions: for Power SYBR Green Mastermix, 2 minutes at 50°C, 10 minutes at 95°C, 40 cycles of 15 seconds at 95°C and 1 minute at 60°C; and for Taqman Gene Expression Assay Mastermix, 2 minutes at 50°C, 10 minutes at 95°C, 40 cycles of 15 seconds at 95°C and 1 minute at 60°C. The primers and probes used (Thermo) are listed in Table 1. For the analysis of orangutan AxSCs, human primers were used, except for PAX6. The Delta-Delta Ct method was employed to calculate the relative expression levels, which were normalized to the GAPDH housekeeping gene. The data were visualized using GraphPad Prism 8.0.

**Table 1:**
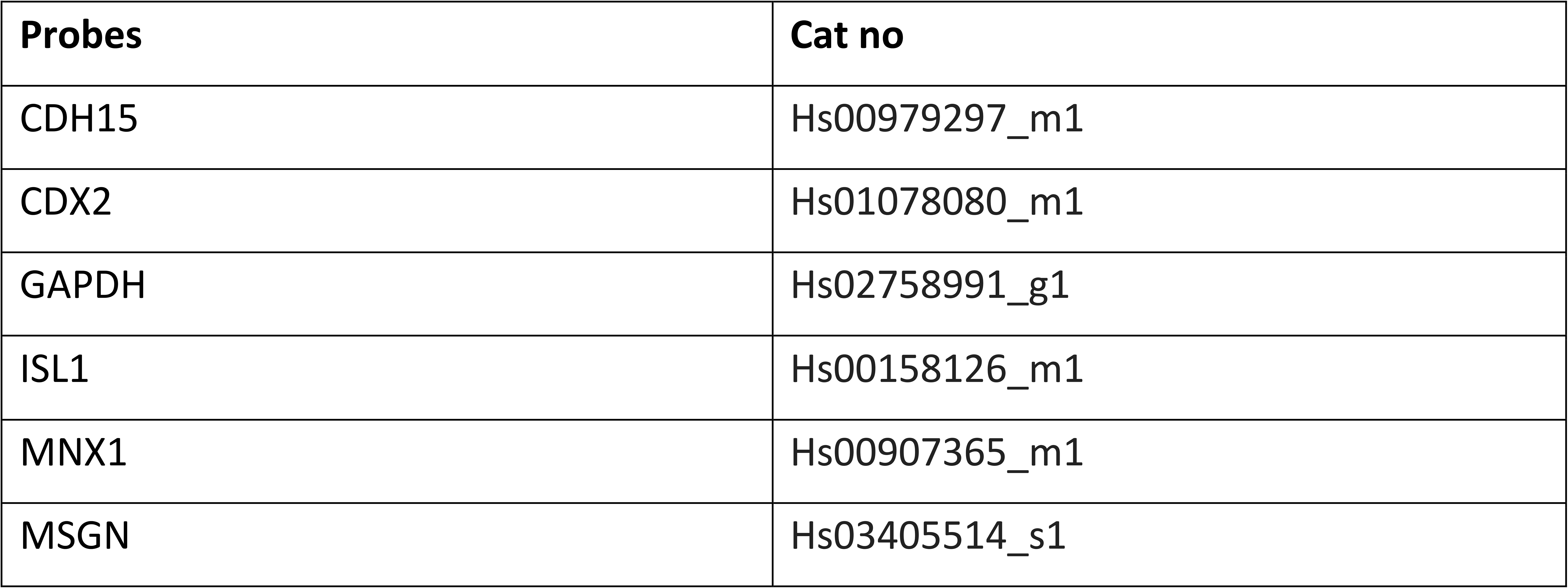
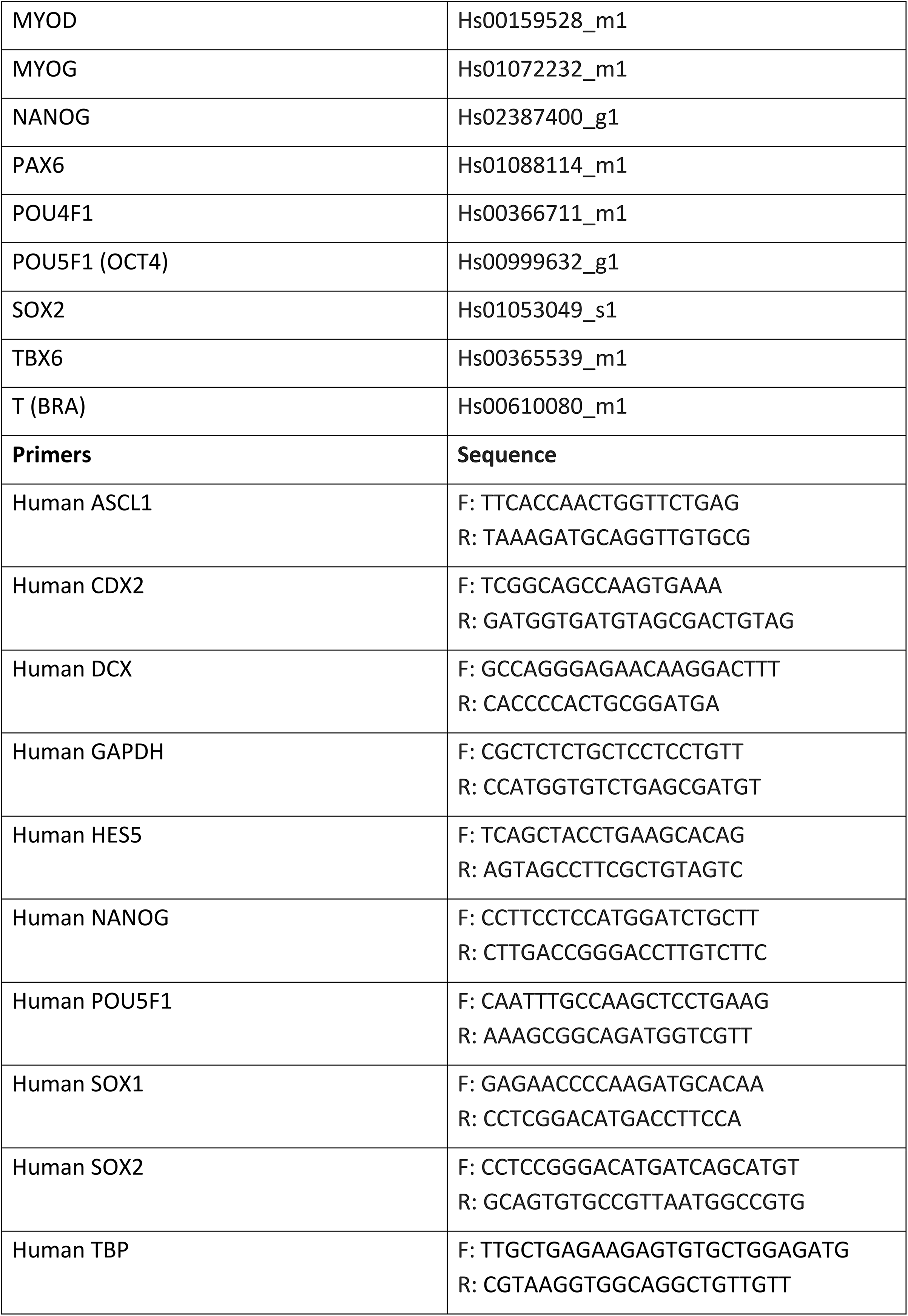
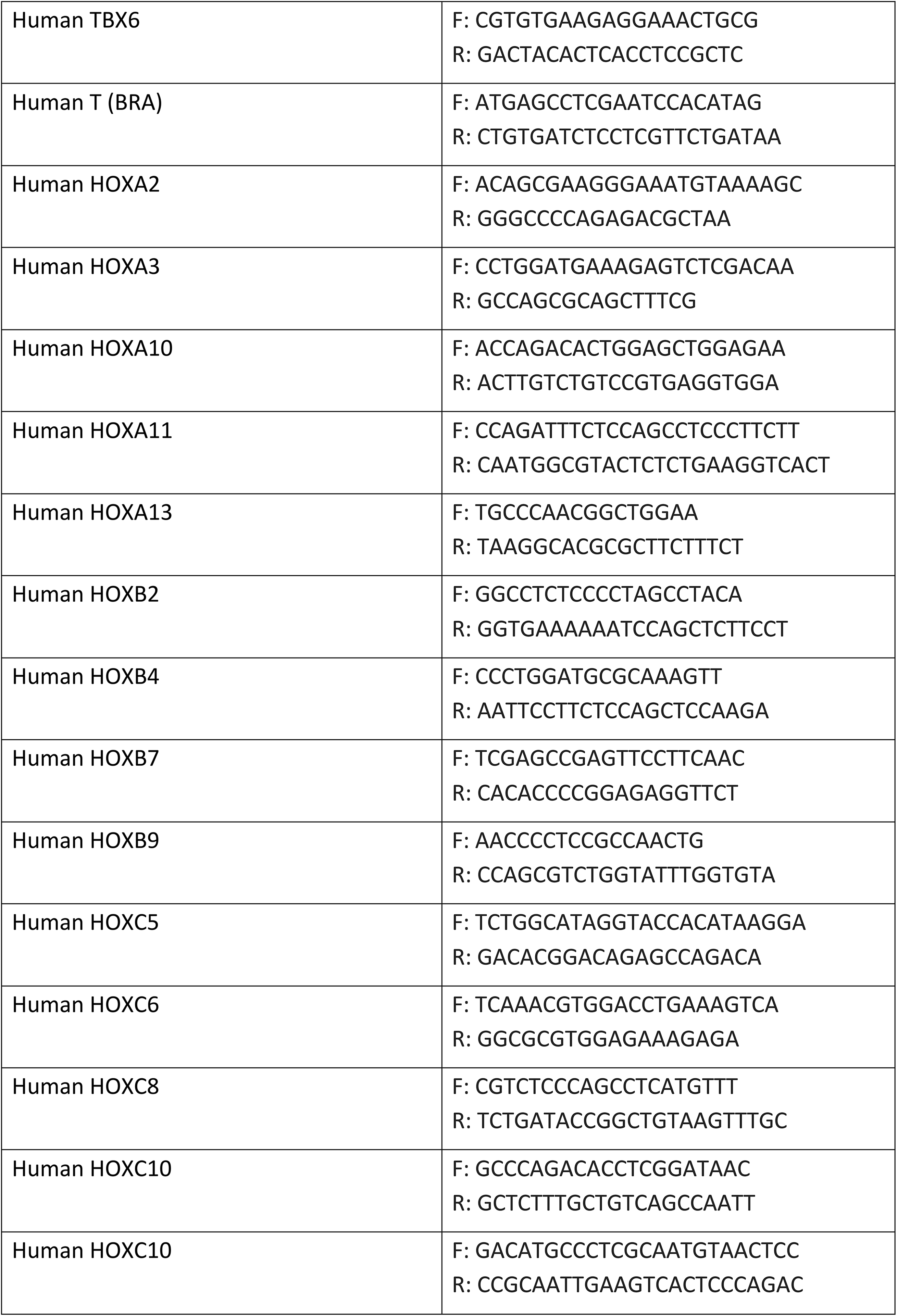
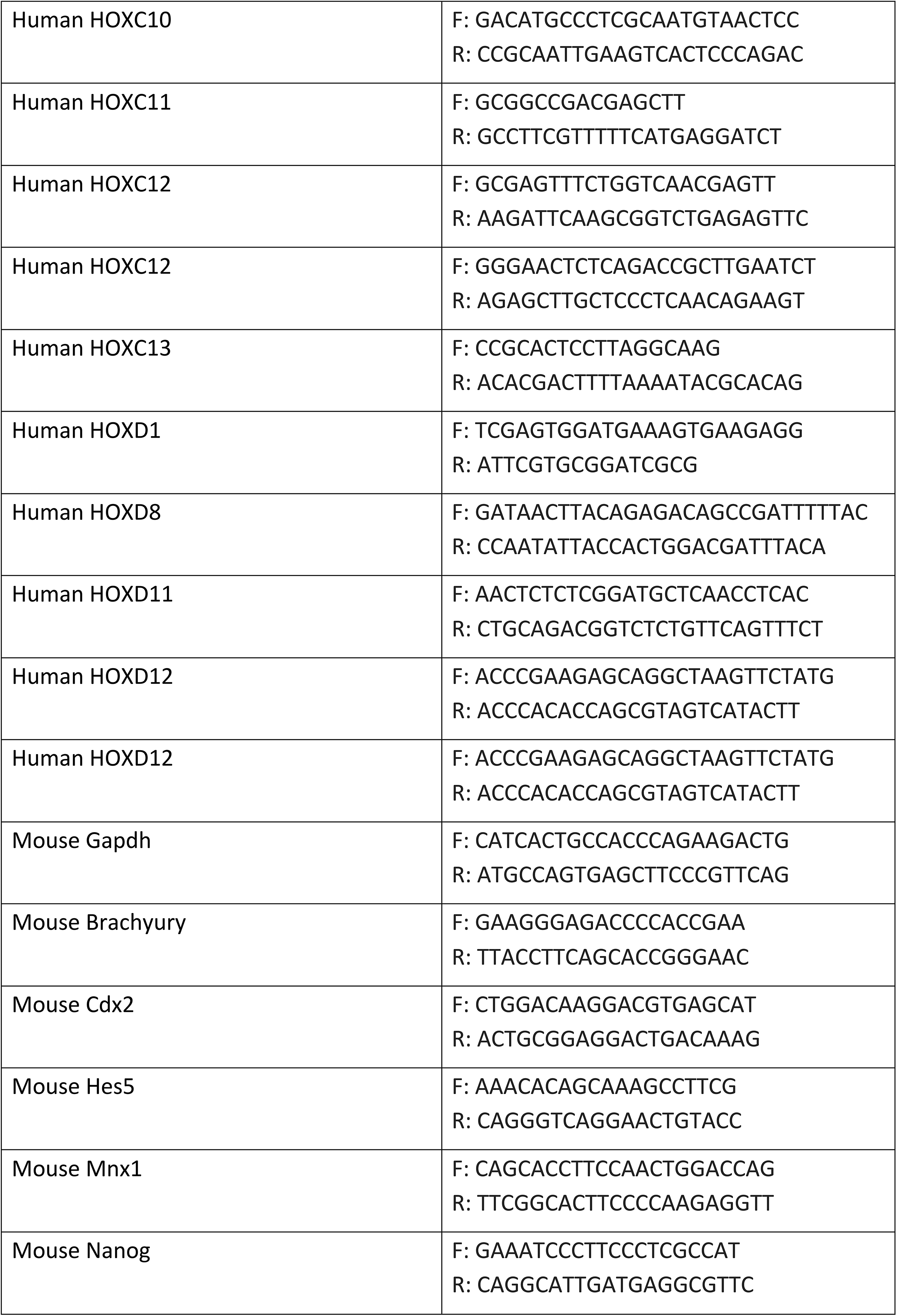
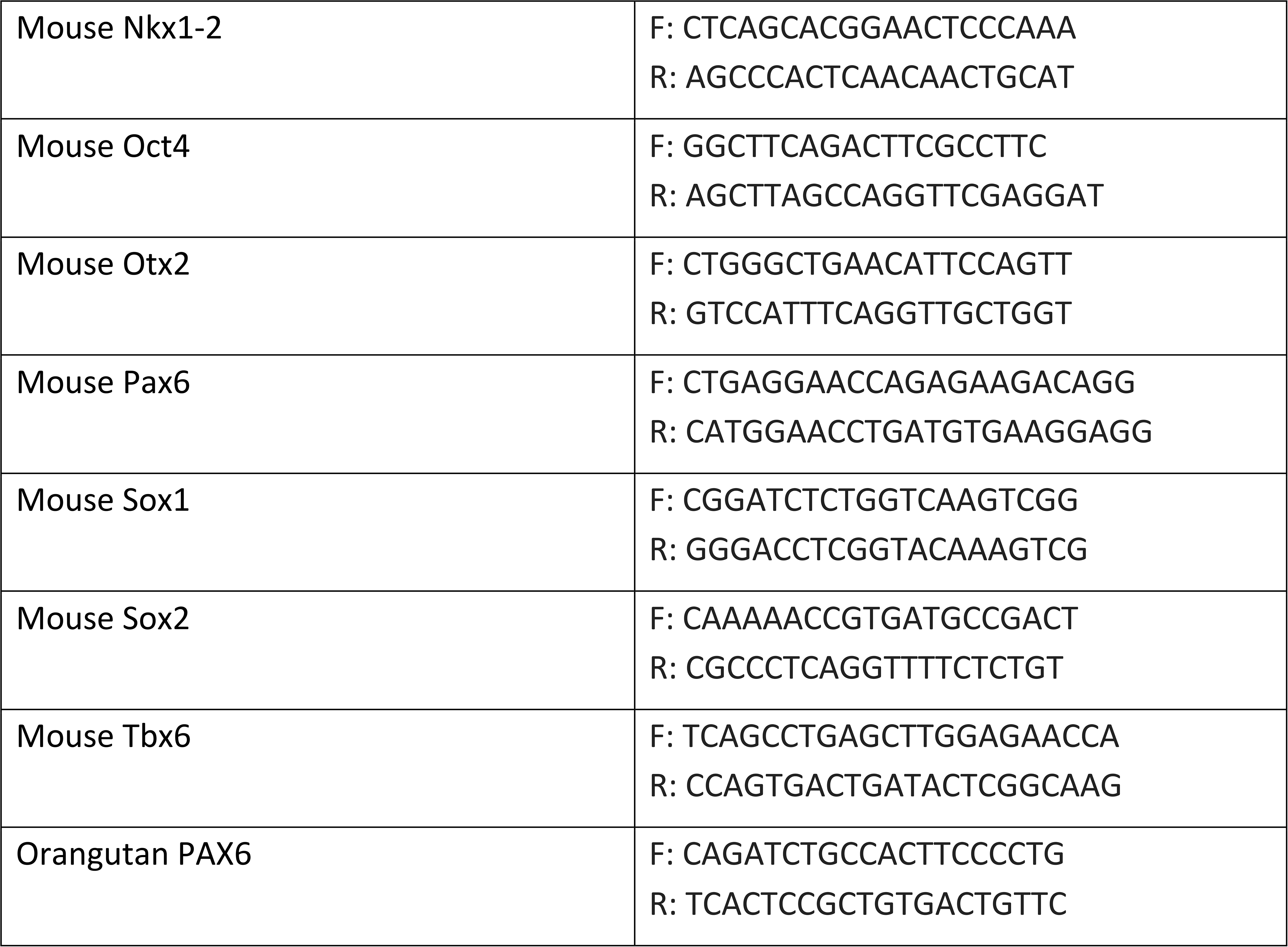
List of probes and primers.

### Immunocytochemistry

For the goat anti-Brachyury antibody (R&D Systems, Table 2), the following protocol was employed. Cells cultured on either coverslips or chamber slides were washed with 0.1% BSA in PBS and fixed with 4% paraformaldehyde (Thermo) for 20 minutes at room temperature (RT). After two washes with 0.1% BSA in PBS, permeabilization was carried out using 0.2% Triton X-100 (Sigma) in PBS for 10 minutes. Samples were blocked with 5% BSA in PBS for 1 hour at RT and incubated overnight at 4°C with primary antibodies diluted in 0.5% BSA/1% Tween-20 (Sigma) in PBS. Following two washes with 1% Tween-20 in PBS for 5 minutes each, the cells were treated with secondary antibodies diluted 1:750 in 0.5% BSA/1% Tween-20 in PBS for 3 hours at RT. After two additional washes with 1% Tween-20 in PBS for 5 minutes each and one wash for 10 minutes, coverslips or chamber slides were mounted using Prolong Gold Antifade and incubated overnight at RT. For other antibodies, cells cultured on either coverslips or chamber slides were washed twice with PBS, followed by fixation with 4% formaldehyde in PBS for 15 minutes at RT. After washing with PBS, cells were permeabilized with 0.2% Triton X-100 in PBS for 5 minutes at RT. Subsequently, the cells were blocked with 3% BSA for 30 minutes at RT and incubated overnight at 4°C with primary antibodies diluted in 0.1% Triton X-100/3% BSA in PBS. The next day, cells were washed three times with PBS for 5 minutes each and incubated with secondary antibodies diluted 1:1000 in 0.1% Triton X/3% BSA in PBS for 1 hour at RT. After washing with PBS three times for 10 minutes each, coverslips or chamber slides were mounted using Prolong Gold Antifade and incubated overnight at RT. The primary and secondary antibodies used are listed in Table 2. Imaging was performed using an Axio Observer Z1 microscope.

**Table 2:** List of antibodies.

| Primary antibodies | Cat no | Manufacturer | Dilution |
| --- | --- | --- | --- |
| Rabbit anti-Sox2 | 2748 | Cell Signaling Technology | 1:400 |
| Mouse anti-Sox2 | MAB2018 | R&D Systems | 1:100 |
| Mouse anti-Sox2 | 4900 | Cell Signaling Technology | 1:400 |
| Rabbit anti-Brachyury | ab209665 | Abcam | 1:100 for immunocytochemistry<br>1:400 for FACS |
| Goat anti-Brachyury | AF2085 | R&D Systems | 1:100 |
| Rabbit anti-CDX2 | 12306 | Cell Signaling Technology | 1:200 |
| Rabbit anti-Pax6 | 901301 | Biolegend | 1:100 |
| Rabbit anti-Islet 1 | ab20670 | Abcam | 1:200 |
| Mouse anti-Brn3a | sc-8429 | Santa cruz | 1:100 |
| Mouse anti-Mnx1 | sc-515769 | Santa cruz | 1:100 |
| Chicken anti-B3 tubulin | NB100-1612 | Novus | 1:100 |
| Mouse anti-Myod | MA1-41017 | Life Technologies | 1:100 |
| Rabbit anti-M-Cadherin | 40491S | Cell Signaling Technology | 1:100 |
| Mouse anti-Myogenin | 14-5643-82 | Life Technologies | 1:100 |
| Mouse anti-MYHC | MAB4470 | R&D Systems | 1:100 |
| Mouse anti-Pax3 | AB_528426 | DSHB | 1:20 |
| Goat anti-Meox1 | sc-10185 | Santa cruz | 1:50 |
| Mouse anti-neurofilament | AB_531793 | DSHB | 2 µg/ml |
| Rabbit IgG isotype control | GTX35035 | GeneTex | 1 µg/ml |
| Mouse IgG isotype control | GTX35009 | GeneTex | 1 µg/ml |
| **Secondary antibodies** | **Cat no** | **Manufacturer** | **Dilution** |
| Goat anti-Chicken IgY (H L) Secondary Antibody, Alexa Fluor 594 | A11042 | Life Technologies | 1:1000 |
| Goat anti-rabbit IgG (H+L), Alexa Fluor 488, highly cross-absorbed | A11034 | Life Technologies | 1:1000 |
| Goat anti-mouse IgG (H+L) Alexa Fluor Plus 647, highly cross-absorbed | A32728 | Life Technologies | 1:1000 |
| Donkey anti-Mouse IgG (H+L) Highly Cross-Adsorbed Secondary Antibody, Alexa Fluor™ 488 | A21202 | Life Technologies | 1:1000 |
| Donkey anti-Goat IgG (H+L) Cross-Adsorbed Secondary Antibody, Alexa Fluor™ 594 | A11058 | Life Technologies | 1:1000 |
| F(ab')<sub>2</sub>-Goat anti-Rabbit IgG (H+L) Cross-Adsorbed Secondary Antibody, Alexa Fluor™ 647 | A21246 | Life Technologies | 1:1000 |
| Alexa Fluor® 647 AffiniPure Goat Anti-Mouse IgG (H+L) | 115-605-146 | Jackson | 1:500 for immunocytochemistry<br>1:1000 for FACS |
| Goat anti-Mouse IgG (H+L) Cross-Adsorbed Secondary Antibody, Alexa Fluor™ 488 | A11001 | Life Technologies | 1:500 |
| Cy™3 AffiniPure Goat Anti-Mouse IgG, F(ab')<sub>2</sub> fragment specific | 115-165-006 | Jackson | 1:500 |
| Goat anti-Rabbit IgG (H+L) Cross-Adsorbed Secondary Antibody, Alexa Fluor™ 488 | A11008 | Life Technologies | 1:1000 |

### Immunocytochemistry for neuromuscular junction analysis

Culture medium was removed, and cells were washed once with 1X PBS. Cells were fixed with 2% paraformaldehyde (Santa cruz) in PBS for 20 min on ice. Cells were washed once with PBS and subsequently permeabilized and blocked with 0.1% Triton X-100 (Sigma) and 1% goat serum (Cell Signaling Technologies) in PBS for 15 min on RT. ActinRed 555 ReadyProbes reagent (Invitrogen) was used for fluorescent staining of actin. Primary (mouse anti-neurofilament (DSHB)) and secondary antibodies (Alexa Fluor® 647 AffiniPure Goat Anti-Mouse IgG (Jackson), Goat anti-Mouse IgG (H+L) Cross-Adsorbed Secondary Antibody (Life Technologies), Alexa Fluor™ 488, Cy™3 AffiniPure Goat Anti-Mouse IgG, F(ab’)₂ fragment specific (Jackson)) were diluted in 0.1% Triton X-100 and 1% goat serum in PBS (Table 2). Primary antibodies were incubated overnight at 4°C, and were washed off after washing three times with PBS (3×10min). Cells were incubated with the secondary antibodies and 2 μg/ml α-bungarotoxin (Invitrogen) for 2 hours at RT. Secondary antibodies were washed off by 3×10 min washing with PBS. Nuclei were stained with Hoechst (1:10000) (Invitrogen) for 10 min in PBS. Cells were imaged using a Nikon Eclipse T*i* confocal microscope.

### Fluorescence Activated Cell Sorting (FACS)

Cells were washed with PBS and dissociated for 5 min at 37°C using TrypLE. Cells were then collected, centrifuged at 300 g for 3 min at RT, resuspended in 1 ml cold 4% PFA for 15 min at RT and then centrifuged at 300 g for 3 min to remove the supernatant. Cells were washed with PBS for three times. Cells were re-suspended in 1 ml of permeabilization buffer (0.2% Triton X-100 - 0.5% BSA in PBS) and incubated for 10 min at RT. Cells were spun again in the same conditions, re-suspended and incubated for 30 min at RT in 0.5% BSA in PBS and then passed through a 40 µm cell strainer. For immunostaining, 10^5^ cells in a total volume of 100 µl were dispensed into each well of 96-well, U-shaped plates. Primary antibodies (mouse anti-SOX2 (Cell Signaling Technology) and rabbit anti-Brachyury (Abcam)) were added to each well. Plates were incubated at RT for 1 hour and then centrifuged for 5 min at 500 g (4°C). For secondary staining, cells were re-suspended in 100 µl 0.5% BSA in PBS containing 1:1000 diluted goat anti-mouse Alex647 (Jackson) and goat anti-rabbit Alex488 (Life Technologies) antibodies for 45 min. Cells were then centrifuged, washed with 200 µl PBS and centrifuged. Finally, the cells were re-suspended in 100 µl FACS buffer containing 0.1% FBS in PBS. Flow cytometry was performed on CytoFlex (Beckman Coulter) using 488 nm and 647 nm lasers. Data analysis was performed using FlowJo.

### Genomic DNA extraction and telomere length assay

DNA extraction was carried out using the Wizard SB Genomic DNA Purification System kit with a minor modification. Prior to elution, to the 250 µl nuclease-free water, 2 µl of RNAse A solution was added, followed by a 10-minute incubation at RT. For the telomere length assay, a total reaction volume of 20 µl was prepared consisting of either 2 µl of telomere primers or 2 µl of SCR primers from the Relative Human Telomere Length Quantification qPCR kit (ScienCell), 1 µl of genomic DNA template (5 ng/µl), 10 µl of Power SYBR Green PCR Mastermix, and 7 µl of nuclease-free water. The reaction was run under the following cycling conditions: 2 minutes at 50°C, 10 minutes at 95°C, followed by 40 cycles of 15 seconds at 95°C and 1 minute at 60°C. The relative telomere length, normalized to the telomere-to-single-copy reference ratio, were calculated using the Delta-Delta Ct method.

### RNA sequencing

#### 1. Sample preparation and sequencing

3 µg of total RNA were treated with TURBO DNase (Life Technologies) and purified using RNeasy Minelute RNA cleanup kit (Qiagen). RNA quality was assessed using microcapillary electrophoresis on Agilent 2100 Bioanalyzer with RNA Pico 6000 kit (Agilent) and only RNA with RIN values >8 was processed further. Per RNA-seq library, 1 µg of DNAse-treated RNA was treated with RiboZero Gold (Human/Mouse/Rat) kit (Illumina) to remove rRNAs, followed by RNA cleanup using the RNeasy Minelute RNA cleanup kit. Sequencing libraries were prepared from equal quantities of rRNA-depleted RNA using TruSeq Stranded total RNA LT kit (Illumina) according to manufacturer’s instructions using 11 cycles of enrichment PCR. The quality of the libraries was assessed using Agilent 2100 Bioanalyzer with the DNA 1000 kit (Agilent). Library concentration was measured using Qubit dsDNA HS Assay Kit (Life Technologies). Multiplexing of libraries was performed according to manufacturer’s instructions. Multiplexed libraries were sequenced using a NextSeq 500 (Illumina) to generate 75-nt single-end reads. Sequencing depth was 20–40 Mio reads per library.

#### 2. Processing and downstream analysis

The reads were pseudo-aligned to human transcriptome (Ensembl version GRChg38.86) using *kallisto (version 0.43.0_3).*^97^ Resulting estimated transcript-level abundances were aggregated into counts per gene and exported using Tximport (*version 1.10.1*) pipeline.^98^ Differential gene expression analysis was performed using DESeq2 package (*version 1.22.2*)^99^ in R by comparing gene expression at each stimulation time point to undifferentiated parental cell line. Normalized counts were extracted from DESeq2 object. PCA plot was constructed from rlog-transformed raw counts.

Analysis of the over-represented pathways was performed using Genomatix Pathway System (GePS) tool from the Genomatix software suite (http://www.genomatix.de) by providing significantly up-regulated genes from each time point and stimulation as input. Tissue and cell type associations were analysed using Enrichr tool^100^ and JENSEN tissue expression database (https://tissues.jensenlab.org). Gene lists associated with selected cell types were extracted and their normalized expression values (from DESeq2 analysis) were used to construct heatmap of the Figure S1C.

### Single cell RNA sequencing

#### 1. Preparation of single cell suspensions

All parental lines, including H9 and AxSCs derived from H9, HMGU1, and HUES6, as well as Day2 neural differentiation samples, were dissociated into single cells using TrypLE Express. The harvested cells were then centrifuged at 300 g for 3 minutes, and the resulting pellet was resuspended in a solution of 0.04% BSA in PBS. The cell density was adjusted to 1000 cells/µl. For Day 14 and Day 28 CFS-derived cells, as well as Day 28 CS-derived cells, dissociation was carried out using 0.05% Trypsin. After centrifugation at 300 g for 3 minutes, the pellet was resuspended in a solution of 0.04% BSA in PBS. The cells were then passed through 40 µm strainers, and the cell density was adjusted to 1000 cells/µl. For Day 14 CS-derived cells, we adapted a protocol that uses Accutase and Papain as dissociation agents.^113^ The cells were first rinsed with PBS and then incubated at 37°C for 25 minutes in a dissociation buffer composed of 50% Accutase (Sigma) and 10% Papain (Sigma) in PBS. Without removing the dissociation buffer, a wash buffer containing DMEM/F12, 33.3 µg/ml Dnase II (Sigma), and 10 µM Y-27632 was added. The resulting cell suspension was filtered through a 70 µm strainer and centrifuged at 300 g for 3 minutes. The pellet was then resuspended in neural differentiation medium and filtered once again using 40 µm strainers.

#### 2. Library preparation and sequencing

##### 2.1. Single cell RNA sequencing of parental human AxSC lines

Live cells were counted using trypan blue stain (Life Technologies) with the Countess 3 automated cell counter (Thermo Fisher Scientific). Samples with viability above 85% were used to prepare cell suspensions containing 1000 cells/µL, which were subsequently used for the single cell library preparation. The leftover cells were pelleted via centrifugation and collected for DNA isolation for genotyping purposes with HumanCytoSNP-12 v2.1 BeadChip Kit (Illumina). The single cell library preparation was done using Chromium Controller instrument, Chromium Next GEM Single Cell 3ʹ GEM, Library & Gel Bead Kit v3.1 (10X Genomics) and the Single Index Kit T Set A (10X Genomics). The final sample loaded for the 10X library prep contained a mixture of 7000 cells/cell line for a total of 21000 cells. Quality control for the library was performed using Agilent High Sensitivity kit (Agilent) following manufacturer instructions. The final pools were sequenced on NovaSeq 6000 platform using SP Reagent Kit v1.5/ 100 cycles (Illumina) with 28×8×91 configuration run leading to 40000 reads/cell.

##### 2.2. Single cell RNA sequencing of neural differentiation time course from human AxSC lines

Counting was performed with Trypan blue for live – dead cell discrimination using Countess 3 automated counter. Samples with viability above 81% were used to prepare cell suspensions containing 500-1000 cells/µL. The single cell library preparation was performed using Chromium Controller instrument, Chromium Next GEM Single Cell 3ʹ GEM, Library & Gel Bead Kit v3.1 (10X Genomics) and Dual Index Kit TT Set A, 96 rxns (10X Genomics). Quality control for the library was performed using Agilent High Sensitivity kit (Agilent) following manufacturer instructions. The final pool contained 16 samples each with 10000 cells per time point and they were sequenced on NovaSeq 6000 platform using NovaSeq 6000 S4 Reagent Kit v1.5 /200 cycles (Illumina) with 28×10×10×90 configuration run leading to 70000 reads/cell.

##### 2.3. Single cell RNA sequencing of mouse AxSC lines

Counting was performed with Trypan blue for live-dead cell discrimination using Countess 3 automated cell counter. Samples with viability above 90% were used to prepare cell suspensions containing 1000 cell/µL. The single cell library preparation and the respective quality control was performed similarly as the neural differentiation time course in 2.2. The final pool was sequenced on the NovaSeqX Plus using 10B Reagent Kit (100 Cycle) with 28×10×10×90 configuration run leading to 25000 reads/cell.

#### 3. Processing of the datasets

For the dataset that includes human parental AxSC lines from H9, HUES6, HMGU1 PSCs, SNP sequencing was performed using Illumina GSA v.3 for cells of each cell line. Next, demuxlet^89^ was used to confidently assign each barcode to a specific cell line. Briefly, demuxlet harnesses genetic variation to identify the genetic origin of individual cells in a mixed sample. Ambiguous cells that could not be confidently matched to one of the three cell lines were removed.

For time course differentiation datasets, sequencing reads were annotated using CellRanger software (v 6.1.2, 10X Genomics) provided with the custom reference (GRCh38 assembly, NCBI annotation of December 2021). The subsequent analysis of the datasets was performed in the Scanpy^94^ environment. Low quality cells (mitochondrial genes > 20%, ribosomal genes > 10% and parameters in Table) were removed. The following thresholds were applied for cell quality filtering: for the day 2 to day 28 datasets, a minimum of 2,000 genes and a total of 80,000 counts; for day 0 CS, a minimum of 800 genes and a total of 60,000 counts; and for day 0 CFS, a minimum of 1,000 genes and a total of 60,000 counts. Additionally, cell doublets were predicted with Solo software^95^ and filtered out. Genes, present in less than 20 cells were removed. Gene counts were normalized to 10^5^ reads per cell and log transformed.

The single cell RNA sequencing data from mouse cell lines data were processed using 10x Genomics Cell Ranger v7.1.0, with the reference genome GRCm39 (Ensembl release 111) and default parameters.

#### 4. Downstream analysis

Downstream analysis was performed using the Scanpy (version 1.6.1), Pandas (version 1.1.3), Numpy (version 1.19.5), and Louvain (version 0.7.0) packages. The datasets were visualized using UMAP embedding with the pl.umap() function. Differential expression analysis for H9, CFS, and CS datasets was conducted using the tl.rank_genes_groups() function with default parameters. The list of differentially expressed genes (DEGs) was extracted using the get.rank_genes_groups() function, setting ’reference’ as "rest" and a log2 fold change threshold of 2. The top 50 DEG genes were visualized using the pl.dotplot() function. Genes associated with axial development, neural/mesodermal lineages (Fig. 3a), were summarized based on published scRNA sequencing studies conducted on *in vivo* cells.^14,22,73,74^ To assess the expression of dorsoventral spinal cord progenitor genes, the study by Rayon et al.^43^ was used as a benchmark.

For the time course neural differentiation experiment, each dataset was analyzed individually. Cell cycle phases were identified using the tl.score_genes_cell_cycle() function with default parameters, and the percentage of cells per phase was calculated using the value_counts() function. Clustering for each dataset was performed using the tl.louvain() function with a ’resolution’ set to 2. Marker genes that map to each domain of spinal cord neurons identified by Rayon et al.^43^, as well as the accompanying database (https://shiny.crick.ac.uk/scviewer/neuraltube/), and marker genes for neural crest development identified by Soldatov et al.^47^ were utilized for cluster annotation and summarized in Table S1. To evaluate specific cell types, single genes known to be specific to only one domain in the database mentioned above were used for analysis. For cell populations that could not be identified by a single gene, the coexpression of several genes was assessed. Clusters were manually merged and annotated based on the expression of marker genes in the datasets. Clusters representing the same cellular population across datasets were shown in the same color.

#### 5. Integration of datasets from multiple species

##### 5.1. Quality Control and Dimensionality Reduction Clustering

We utilized Seurat V5^90^ to import the counts matrix processed by Cellranger. DoubletFinder^92^, with its default parameters for 10X data, was employed to predict doublets, and cells with medium to high doublet probabilities were removed. Subsequently, we retained cells that met the criteria of nFeatures ≥ 500, percent.mt < 5, and nCount < 30,000. Additionally, mitochondrial and ribosomal genes were excluded from the counts matrix to prevent their influence on downstream dimensionality reduction and clustering results. Then the processed gene expression profile matrix was then normalized, and we experimented with batch correction methods including Harmony^114^, Seurat^115^, and SCT v2^91^. We compared these results with those obtained without batch correction, where Seurat was directly used for nonlinear dimensionality reduction via PCA and UMAP. Finally, we achieved the most optimal results using Seurat’s batch correction method. Based on this, we employed Seurat’s FindClusters function to determine cell types for each cluster. This was achieved by repeatedly analyzing classic markers at different resolutions and taking into account the spatial positions of these clusters.

##### 5.2. Cross-dataset/species comparative analysis

Using the MapQuery function from Seurat^90^, we mapped our sequencing results onto various reference datasets to obtain their ref.UMAP relative to these references. We then used a custom ggplot function to project them onto the same UMAP plane, visualizing their degree of similarity. For cross-species datasets, we first downloaded ortholog tables for human, monkey, and mouse genes from the Ensembl BioMart page (http://www.ensembl.org/biomart/martview) and converted all species’ genes to their human orthologs. Subsequently, we mapped our data onto datasets of other species using the same methodology.

We employed a custom ComplexHeatmap^93^ function to generate similarity heatmaps based on the confusion matrix between predicted cell types and our annotated cell types. For highly ranked Map Percent values within the matrix, specific numbers were annotated. For cross-species gene comparison, we utilized the heatmap and violin plot functions from the R package SCP v0.5.6 (https://github.com/zhanghao-njmu/SCP) to depict the expression of classic signaling genes, EMT genes, and others for each dataset. These were then combined into a single heatmap.

### Chromatome and proteome analysis

#### 1. Nuclei isolation and sample preparation for chromatome analysis

AxSCs and H9 cells were dissociated into single cells using TrypLE Express. The cells were incubated for 4-6 minutes at 37°C. The resulting cell suspensions were centrifuged at 300 g for 3 minutes, and the cell number was counted. For each derivation of CS and H9, 5×10^6^ cells were used, while for each derivation of CFS cells, 3×10^6^ cells were used. The cells were washed twice with PBS by centrifugation at 2300 g for 5 minutes at RT. The supernatant was discarded, and the cell pellet was resuspended in 1 ml of ice-cold lysis buffer (3 mM MgCl2, 10 mM NaCl, 10 mM Tris pH 7.4, 1% NP-40, and freshly added 1X cOmplete protease inhibitor (Sigma)). To ensure thorough dissolution of NP-40, the lysis buffer was pre-incubated on an orbital shaker at mild agitation at 4°C before being used for resuspending the cell pellet. The pellet was homogenized by pipetting up and down and then incubated on ice for 20 minutes. The suspension was centrifuged at 2300 g for 5 minutes at 4°C, and the supernatant was discarded. The pellet was resuspended and incubated in 3.3 ml of PBS with 1% methanol-free formaldehyde for 10 minutes on a rotating wheel at mild agitation at RT. The reaction was quenched by incubating the suspension with 125 mM Glycine for an additional 5 minutes on a rotating wheel. The nuclei suspension was then centrifuged at 2300 g for 5 minutes at 4°C and washed twice with ice-cold PBS. After another centrifugation step at 2300 g for 5 minutes at 4°C, the supernatant was discarded, and the pellets were snap-frozen in liquid nitrogen for 15 seconds and then stored at -80°C. Chromatin was released from crosslinked nuclei by dissolving 300 μl of SDS buffer (50 mM HEPES pH 7.4, 10 mM EDTA pH 8.0, 4% UltraPure™ SDS Solution from Invitrogen, along with a newly added 1X cOmplete™ EDTA-free Protease Inhibitor Cocktail) using gentle pipetting. This mixture was left to incubate at RT for 10 minutes before adding 900 μl of freshly prepared Urea buffer (10 mM HEPES pH 7.4, 1 mM EDTA pH 8.0, 8 M urea from Sigma). The solution was then carefully inverted seven times before being centrifuged at RT for 30 min at 20000 g. The supernatant was removed, taking care not to disturb the pellet. Two additional wash steps were performed (one wash with SDS and Urea, and one wash with only SDS). The final pellet was then dissolved in 300 μl of Sonication buffer (10 mM HEPES pH 7.4, 2 mM MgCl2, with a freshly added 1× cOmplete™ EDTA-free Protease Inhibitor Cocktail). The chromatin specimens were sonicated using a Bioruptor® Plus at 4°C for 15 cycles (30 s on, 30 s off). The protein concentration was determined using the Pierce™ BCA Protein Assay Kit.

Subsequently, Protein Aggregation Capture (PAC) was performed. In this step, Sera-Mag™ beads (1 mg from Sigma) were washed three times with 70% acetonitrile for every 100 μg of chromatin solution. After the final wash, 300 μl of the chromatin solution corresponding to 100 μg was added to the beads, followed by 700 μl of 100% acetonitrile. The chromatome-bead mixtures were then vortexed and left to rest on a bench for 10 minutes. The samples were vortexed again and placed into a magnetic rack. The samples were then washed with 700 μl of 100% acetonitrile, followed by 1 ml of 95% acetonitrile, and finally with 1 ml of 70% ethanol. The remaining ethanol was allowed to evaporate, and the beads were resuspended in 400 μl of 50 mM HEPES pH 8.5, supplemented with freshly prepared 5 mM TCEP and 5.5 mM CAA. The samples were then left to incubate at RT for half an hour. Protease digestion was initiated by adding LysC (protease to protein ratio of 1:200) and Trypsin (1:100) and allowing the mixture to incubate overnight at 37°C under constant agitation at 1100 rpm. From this point forward, the samples were handled in the same way as the total proteome samples.

#### 2. Sample preparation for total proteome analysis

All proteomic experiments were conducted in triplicate. AxSCs and H9 cells were dissociated into single cells using TrypLE Express. The cells were incubated for 4-6 minutes at 37°C. After incubation, the cell suspensions were centrifuged at 300 g for 3 minutes, and the cell number was determined. For each derivation of CS and H9, 2×10^6^ cells were used, while for each derivation of CFS cells, 1.5×10^6^ cells were used. These cells were then centrifuged at 300 g for 3 minutes. The supernatant was removed, and the cell pellets were snap-frozen in liquid nitrogen for 15 seconds, followed by storage at -80°C. Frozen cells were dissolved in 200 μl of lysis buffer (containing 6 M guanidinium chloride, 100 mM Tris-HCl with a pH of 8.5, and 2 mM DTT) and subjected to heating for 10 minutes at 99°C with a constant agitation rate of 1400 rpm. The sonication of samples was then performed at 4°C using a Bioruptor® Plus sonication device (Diagenode) at high-intensity settings, with 30 seconds on/off intervals for 15 rounds. If the sample viscosity was adequately reduced, protein concentrations were determined; if not, sonication was repeated. Protein concentrations were assessed using the Pierce™ BCA Protein Assay Kit (Thermo Fisher Scientific) in accordance with the manufacturer’s instructions. After incubating for at least 20 minutes with 40 mM chloroacetamide, 30 μg of each proteome sample was diluted in a 30 μl lysis buffer supplemented with chloroacetamide and DTT. These samples were further diluted in 270 μl of digestion buffer (containing 10% acetonitrile, 25 mM Tris-HCl at pH 8.5, 0.6 μg Trypsin/sample (Pierce™ Trypsin Protease, Thermo Fisher Scientific), and 0.6 μg/sample LysC (Pierce™ LysC Protease, Thermo Fisher Scientific). Proteins were then digested for 16 hours at 37°C with constant shaking at 1100 rpm.

To halt protease activity, 1% (v/v) trifluoroacetic acid (TFA) was added the following day and samples were loaded onto homemade StageTips composed of three layers of SDB-RPS matrix (Empore)^116^, previously equilibrated with 0.1% (v/v) TFA. After loading, two washes with 0.1% (v/v) TFA were performed, and peptides were eluted with 80% acetonitrile and 2% ammonium hydroxide. After the eluates were evaporated in a SpeedVac centrifuge, the samples were resuspended in 20 μl 0.1% TFA and 2% acetonitrile. The peptides were completely solubilized by constant shaking for 10 minutes at 2000 rpm, and peptide concentrations were determined on a Nanodrop™ 2000 spectrophotometer (Thermo Fisher Scientific) at 280 nm.

#### 3. Nanoflow LC–MS/MS analysis for proteomes and chromatomes

Peptide separation before MS was accomplished using liquid chromatography on an Easy-nLC 1200 (Thermo Fisher Scientific) with in-house packed 50 cm columns of ReproSilPur C18-AQ 1.9-μm resin. A binary buffer system was used (buffer A: 0.1% formic acid and buffer B: 0.1% formic acid in 80% acetonitrile), with a gradual increase in buffer B concentration (from 5% initially to 95% at the end) to elute the peptides over a 120-minute period at a steady flow rate of 300 nl/min. The peptides were then introduced into an Orbitrap Exploris™ 480 mass spectrometer (Thermo Fisher Scientific) via a nanoelectrospray source. Each set of triplicates were followed by a washing step while the column temperature was constantly at 55°C.

Data-Dependent Acquisition (DDA) runs used a top12 shotgun proteomics method within a range of 300–1650 m/z, a default charge state of 2, and a maximum injection time of 25 ms. Full scan resolution was set at 60000 and the normalized AGC target at 300%. For MS2 scans, the orbitrap resolution was set at 15000 and the normalized AGC target at 100%, with a maximum injection time of 28 ms.

Data-Independent Acquisition (DIA) runs used an orbitrap resolution of 120000 for full scans in a scan range of 350–1400 m/z, with a maximum injection time of 45 ms. For MS2 acquisitions, the mass range was set to 361–1033 with isolation windows of 22.4 m/z. A default window overlap of 1 m/z was used. The orbitrap resolution for MS2 scans was set at 30000, the normalized AGC target at 1000%, and the maximum injection time at 54 ms.

#### 5. MS data quantification

Raw MS data acquired in DIA mode was analyzed using DIA-NN version 1.8.1.^117^ A hybrid approach utilizing a dedicated DDA and DIA library was employed; the DDA library was generated using SpectroMineTM (Pulsar) and the DIA library using DIA-NN. Cross-run normalization was conducted in an RT-dependent manner. Missed cleavages were set to 1, N-terminal methionine excision was activated, and cysteine carbamidomethylation was set as a fixed modification. Proteins were grouped using the additional command ‘–relaxed-prot-inf’. Match-between runs was enabled, and the precursor FDR was set to 1%.

#### 5. Statistical analyses

Raw data outputs were analyzed downstream with Perseus (version 1.6.0.9).^118^ CVs were calculated by filtering out proteins or precursors with fewer than 2 out of 3 valid values. Downstream analyses were conducted after imputation of missing values, which was done based on a Gaussian distribution with respect to the standard deviations of measured values (width of 0.2 and a downshift of 1.8 standard deviations). GO enrichment analyses of differentially enriched proteins were performed against the background of total identified proteins using a Benjamini-Hochberg FDR-corrected Fisher’s Exact test of 0.05. These analyses were carried out individually for each cluster. Student’s t-tests were calculated with a permutation-based FDR of 0.05 and an s0 value of 1, unless stated otherwise. For the multiple sample test based on an ANOVA the FDR was set to 0.05 and the s0 value to 2. ANOVA tests of normalized chromatomes were conducted likewise after calculating pairwise differences of ChAC-DIA and total proteome values.

#### 6. Web application development

Significant changes in chromatome, proteome and relative chromatin binding during axial stem cell differentiation were displayed in an interactive heatmap as mean row differences of log2 intensities.

The web application was programmed using R Shiny with the following libraries besides base R packages for data processing and visualization: shiny (1.7.1), shinydashboard (0.7.2), shinyHeatmaply (0.2.0), plotly (4.10.0), heatmaply (1.3.0) and png (0.1–7). From the tidyverse (1.3.1) family we further utilized tidyr (1.2.0), dplyr (1.0.9), and ggplot2 (3.3.6).

### Reduced representation bisulfite sequencing (RRBS)

#### 1. Genomic DNA isolation and library preparation

Genomic DNA was isolated from cultured cell monolayers using the Quick-DNA Miniprep Kit (Zymo Research) according to the manufacturer’s instructions and eluted sequentially in 30 μL and 20 μL elution buffer. DNA concentration was measured using the Qubit dsDNA High Sensitivity Assay (Thermo Fisher Scientific).

For RRBS library preparation, 200 ng genomic DNA per sample was digested with HaeIII (New England Biolabs) in 1x CutSmart Buffer (50 μL total volume) at 37°C for 16 h, followed by heat inactivation at 80°C for 30 min. Digested DNA was purified using 1.8x MagMAX magnetic beads (Thermo Fisher Scientific) and eluted in 50 μL nuclease-free water.

End repair and A-tailing were performed using the NEBNext Ultra II End Prep Module (New England Biolabs) in a 60 μL reaction containing 7 μL reaction buffer and 3 μL enzyme mix, with incubation at 20°C for 30 min and 65°C for 30 min. DNA was purified using 1.8x MagMAX beads and eluted in 13.5 μL pre-warmed (45°C) nuclease-free water, of which 12.5 μL was used for adapter ligation.

A-tailed DNA was ligated to methylated TruSeq adapters (Microsynth; 100 μM) using Quick T4 DNA Ligase (New England Biolabs) in a 20 μL reaction and incubated overnight at 20°C. Ligated DNA was subjected to bisulfite conversion using the EZ DNA Methylation-Gold Kit (Zymo Research) according to the manufacturer’s protocol.

Bisulfite-converted libraries were enriched by PCR using the Q5 Ultra II U-Ready Master Mix (New England Biolabs). PCR reactions (50 μL) contained P5 and P7 index primers (30 μM each) and 5 μL bisulfite-treated DNA, and were amplified with an initial denaturation at 98°C for 30 s, followed by 8 cycles of 98°C for 10 s, 65°C for 30 s, and 72°C for 30 s, with a final extension at 72°C for 5 min.

Final libraries were quantified using Qubit dsDNA HS assays and assessed for fragment size distribution using an Agilent TapeStation system.

#### 2. Data processing

Paired-end RRBS data were processed using a Snakemake-based workflow (Snakemake 8.15.2)

1. ^119^. Sample metadata, including sample identifiers, FASTQ file paths, and treatment annotations, were provided via a tab-delimited sample sheet. Raw FASTQ files were staged by symbolic linking to standardized filenames.

Adapter and quality trimming were performed with Trim Galore v0.6.10 (https://github.com/FelixKrueger/TrimGalore) in paired-end mode, using a minimum Phred quality score of 20 and a minimum retained read length of 20 nt. To account for RRBS-specific bias, the first 4 bases of both read 1 and read 2 were removed. Trim Galore dependencies included cutadapt v5.2 ^120^ and FastQC v0.12.1 (https://www.bioinformatics.babraham.ac.uk/projects/fastqc/).

Trimmed reads were aligned to a bisulfite-converted reference genome using Bismark v0.25.1 ^121^ with Bowtie2 v2.5.4 ^122^, configured for directional libraries and paired-end alignment. PCR duplicates were removed using deduplicate_bismark in paired-end mode. Methylation calling was performed with bismark_methylation_extractor v0.25.1 using RRBS-specific settings, generating comprehensive cytosine reports and bedGraph files.

Alignment statistics and per-sample HTML reports were generated using bismark2report. Deduplicated BAM files were sorted and indexed with samtools v1.23 ^123^. Genome-wide coverage tracks were generated using deepTools bamCoverage v3.5.6 ^124^ with a bin size of 25 bp, CPM normalization, exclusion of duplicate reads, a minimum mapping quality of 10, and read extension enabled. The effective genome size was calculated from the reference FASTA index. Methylation BigWig tracks were generated by sorting Bismark bedGraph files and converting them using UCSC bedGraphToBigWig v482, with chromosome sizes derived from the reference FASTA index.

Quality control was performed on raw and trimmed reads using FastQC v0.12.1. Alignment-level QC included samtools flagstat and samtools stats and insert size metrics were computed using Picard CollectInsertSizeMetrics v3.4.0 (https://broadinstitute.github.io/picard/). All QC outputs were aggregated using v1.33 ^125^.

#### 3. Differential methylation calling

Coverage files (tables with methylation counts per genomic site) were generated using bismark2bedGraph. Common human single-nucleotide polymorphisms (UCSC snp151Common) were excluded, and the coverage files were used to identify differentially methylated regions (DMRs) in passages 24in, p2, p4, (CS) p6, and p10 (each compared to p0). Passage CFS p6 was excluded because it was identified as an outlier in the PCA. DMRs were identified using the DSS v2.58 ^126^ package with R v.4.5.2 ^127^ in two steps: First, site-wise statistics were calculated with DMLtest with smoothing enabled. Subsequently, DMRs were called with a site p-value threshold of 0.001, a minimum region length of 100 bp, a minimum of 5 CpG sites per region, and a delta value of 0.1.

#### 4. Transcription factor motif enrichment analysis in RRBS data

To search for transcription factors (TFs) that are associated with the methylation changes during cell differentiation (i.e., potential differentiation drivers), the hypo- and hypermethylated DMRs of passage p10 were searched for TF motifs that are found more frequently than expected. This search was carried out using monaLisa v1.16 ^128^ in R v.4.5.2 ^127^ as follows: Vertebrate transcription factor motifs were retrieved from the JASPAR 2026 database ^129^ using JASPAR v0.99.4 ^130^. The DMRs of passage p10 were loaded and divided into five equal-sized groups (bins) based on their mean methylation change. The group containing the DMRs with an absolute mean methylation change of less than 0.3 was defined as null (no-effect) bin. The DMR groups were verified to have roughly equal average lengths and GC content, though it was apparent that DMRs in groups with stronger methylation changes were on average longer and had lower GC content. Finally, motif enrichment per DMR group was calculated using a minimal motif hit score of 10 and the DMRs of the other groups as background. P-values were calculated by monaLisa using Fisher’s exact test and adjusted with Benjamini-Hochberg multiple testing correction. The enrichment results were filtered for significant (p < 10e-4, unless noted otherwise) enrichment in at least one of the non-null groups. Enrichment results did not change when using the (subsampled) whole genome or all sequenced regions instead of the other DMR groups as background.

Many Hox and Cdx motifs found are very similar. To determine whether their enrichment stemmed from a single underlying motif, motif hits were searched in the DMRs using monaLisa findMotifHits with a minimum motif hit score of 10, and co-occurrence was monitored by calculating Pearson correlation between the Hox/Cdx motif hits.

To monitor the enrichment of Hox and Cdx motifs across passages, the DMRs of the other passages were processed as described for passage p10.

To check for demethylation of the promoters of TFs whose motifs are enriched in the hypomethylated DMRs (i.e., of the potential differentiation drivers), the Bismark coverage files of all passages were combined and filtered for positions within a range of 1000 bp upstream and downstream of the transcriptional start site of the TF. Then, median methylation was calculated over these ranges for each passage.

### CRISPR/Cas9 screening for essential gene analysis

#### 1. Plasmid library amplification, virus production and transduction of CFS cells

Previously constructed gRNA library that is cloned into the lentiCRISPR v1^131^ plasmid was transformed into Endura electrocompetent cells (Lucigen) for expansion. To generate the lentiviral pool, the genome-wide sgRNA library, comprising 182,135 gRNA-containing lentiCRISPR v1 plasmids targeting more than 18,000 coding genes with approximately 10 sgRNAs per gene for nearly all targets was transfected into 293T cells together with pCMV-VSV-G and psPAX2 plasmids at a ratio of 13.3:6.6:10 (30 µg total plasmid in every 15cm culture dish), respectively, in the presence of polyethylenimine ‘Max’ (Polysciences). Transfection medium was exchanged with 0.5% BSA-containing 293T growth medium after 16 h. Lentiviral particles were collected 65 h after transfection by an initial centrifugation at 3,000 rpm. for 10 min at 4 °C and filtering through 0.45 µm cellulose acetate filters (Millipore) to get rid of the cellular debris, followed by an ultracentrifugation of the cleared supernatant at 24,000 rpm. in a swing bucket rotor (Beckman Coulter) for 2h at 4 °C. The lentiviral pellet was reconstituted in cold CFS medium and frozen at -70 °C. Functional virus titers were measured by transducing the CFS cells with different volumes of the virus preparation and by calculating the ratio of puromycin-selected cells to the non-selected cells in parallel wells of transduced cells.^132^ A multiplicity of infection (MOI) of 0.3 was used to transduce the expanded CFS cells to enrich for the mutants containing a single gRNA integration, hence a single mutation. Transduced cells were selected for 7 days with puromycin. 1-1.5 days after the transduction, 40 million cells were harvested for an initial reference point in the analysis, representing the starting pool of gRNAs. 6 and 16 days after the transduction, 72 million cells were harvested per time point, leading to a ∼400-fold coverage of the gRNA library, to test the changes in abundance of the mutants within the population as compared to the initial time point.

#### 2. DNA extraction, PCR amplification of sgRNAs and high-throughput DNA sequencing

Genomic DNA extraction was performed using the Blood & Cell Culture DNA Midi Kit (QIAGEN) according to the instructions of the manufacturer. The region containing the gRNA integration was amplified using the following primers, as described previously^49^: 5ʹ-TCGTCGGCAGCGTCAGATGTGTATAAGAGACAGNNNNNNNNNNGGCTTTATATATCTTGT GGAAAGGACG-3ʹ (forward) and 5ʹ-GTCTCGTGGGCTCGGAGATGTGTATAAGAGACAGACG GACTAGCCTTATTTTAACTTGC-3ʹ (reverse). DNA amplicon libraries containing sgRNA constructs were sequenced using P2.200 flowcell on the Illumina NextSeq 1000. The run configuration was 100bp paired end.

#### 3. Data analysis

Fastq files were analyzed using the nf-core pipeline CRISPRseq v2.2.1, employing the screening analysis parameter. Reference sequences from Wang et al.^131^ were utilized for library annotation via CRISPRcleanR within the pipeline. The parameter min_reads was set to 3 to filter sgRNAs based on their average counts in the control sample. The following softwares and packages were used for the processing: MAGeCKFlute, Anaconda, Bioconda, MAGeCK, CRISPRcleanR, BAGEL2, BioContainers, Docker, Singularity.

### Cell doubling times upon PRC2 inhibition

Cells were initially seeded at a uniform density in 6-well plates and incubated under controlled conditions (37°C, 5% CO₂, 5% O2) to ensure standard growth environments. 5 µM GSK126 and 20 µM EED226 exposure began the next day, with daily replacement of the medium. Cell counts were recorded at the time of harvesting and seeding using an automated cell counter. CDT was calculated based on the exponential growth model, applying the formula CDT = [ T × ( ln2 ) ] / [ ln (Xe / Xb) ], where T = the interval between measurements, Xb is the initial cell count, Xe is the final cell count. Each experiment was performed in triplicate to ensure accuracy, and data were averaged to determine mean doubling times. Statistical analysis was conducted using one-way ANOVA to assess significant differences between groups.

### Electrophysiological analysis

#### 1. Patch-clamp recordings

The whole-cell patch-clamp technique was performed to record from cells either derived from hiPSCs (CS and CFS) or hESCs (labeled 04.06) recorded at an early (day 14-21) and late time point (day 49-57) after start of differentiation. During recordings at RT, neurons grown on glass coverslip were continuously perfused by artificial cerebrospinal fluid (ACSF) saturated with 95% O_2_ / 5% CO_2_ at a flow rate of 2 ml/min (Ismatec IPC). ACSF contained 125 mM NaCl, 25 mM NaHCO_3_, 2.5 mM KCl, 1.25 mM NaH_2_PO_4_, 10 mM D-glucose, 2 mM CaCl_2_, and 1 mM MgCl_2_ (pH, 7.3; osmolality, 310 mOsm/kg).

Whole-cell voltage-clamp recordings were performed with a patch-clamp amplifier (EPC 9, HEKA Elektronik). Individual cells were visualized using infrared differential interference contrast video microscopy on a Zeiss Axioskop microscope and were identified based on a spherical and bright cell body bearing 2 or more neurites. Borosilicate glass electrodes with filaments (1B150F-3, 0.84 x 1.5 x 76mm, Science Products) were pulled with a horizontal pipette puller (Model P-97 Flaming Brown Micropipette Puller, Sutter Instrument). Pipettes had resistances between 4–6 MΩ when filled with solution containing (in mM): 115 mM K-gluconate, 20 mM KCl, 10 mM Phosphocreatine-Na, 4 mM Mg-ATP, 0.3 mM GTP, 0.2 mM EGTA, and 10 mM HEPES (pH, 7.3; 290 mOsm).

In voltage-clamp, input resistance was determined at -70 mV with voltage steps of -3 mV before inward and outward currents were evoked with increasing depolarizing steps (20 mV; 300 ms). In current-clamp, the resting membrane potential (RMP) was determined and APs were evoked with increasing current steps (20 pA; 300 ms). The amplitude of the first evoked AP was analyzed. Spontaneous excitatory postsynaptic currents (sEPSCs) were recorded at -70 mV.

#### 2. Data analysis

Offline analysis was done with FitMaster (HEKA Elektronik). Spontaneous synaptic activity was analyzed using Igor Pro (WaveMetrics, Lake Oswego, OR) and MiniAnalysis (Synaptosoft). GraphPad Prism (version 8.2.1) was used to plot the data as mean +/- SEM and to test for statistical differences among means of groups via one-way analysis of variance (ANOVA).

### Teratoma analysis

#### 1. Injection

Animal experiments were performed following the experimental guidelines within the animal protocol approved by the government of Upper Bavaria with nr. ROB-55.2-2532.Vet_02-14-41. In brief immunodeficient SCID mice were injected under the kidney capsule with 1×10^6^ cells resuspended in 50% Matrigel. The mice were kept under observation for the duration of 6 weeks.

#### 2. Histopathology

The left and right kidney were harvested and immediately fixed with 4 % (w/v) neutrally buffered formalin for a maximum of 48 hours. Kidneys were cut in a standardized manner resulting in 4 cross-sections and subsequently routinely embedded in paraffin. Sections of 3 µm were stained with hematoxylin and eosin (H&E) and scanned with an AxioScan.Z1 digital slide scanner (Zeiss) equipped with a 20x and a 40x magnification objective. Images were obtained using the commercially available software netScope Viewer Pro (Net-Base Software GmbH).

### Energy metabolism analysis

The cells (H9, CS and CFS) were dissociated using TrypLE select and seeded in cell culture microplate (Agilent Seahorse XF96) at a density of 50000 cells/well in presence of their full respective media, supplemented by 10 µM of Y-27632. The next day the cells (90% confluence) were incubated with RPMI medium (Sigma) supplemented with L-Glutamine (2 mM) with D-(+)-Glucose (10 mM, for the measurement of Oxidative phosphorylation, or without Glucose (for glycolysis assay) and transferred to an incubator (37 °C) without CO_2_ for one hour before the start of the measurement. For Mito Stress assay, the following treatments were used, respectively: Oligomycin at 1 µM (Cayman Chemical) which inhibits the ATP synthase, Carbonyl cyanide 4-(trifluoromethoxy) phenylhydrazone (FCCP, 1µM, Sigma), an uncoupling agent that disrupts mitochondrial ATP synthesis by interfering with proton gradient across the mitochondrial inner membrane^133^ and a combination of rotenone (Sigma) and antimycin A (Sigma) inhibiting complex I and complex III of the electron transport chain, respectively, each at a concentration of 1 µM. For glycolysis stress test, the cells Glucose (10 mM) was first injected followed by Oligomycin (1 µM) then 2-Deoxy-D-glucose at 10 mM (Sigma). At least 30 wells were assayed for each cell type. The measurements values were then normalized to the total protein content in each well measured by the SRB assay. The data were then visualized using Wave software and analyzed using GraphPad Prism 8.0.

### Embryo grafting and assessment of contribution

#### 1. Mouse strains, staging and husbandry

Wild-type C57BL/6J mice were maintained on a 12 h light/12 h dark cycle. For timed matings, noon on the day of finding a vaginal plug was designated as E0.5. Staging of early mouse embryos was carried out according to Downs and Davies.^134^ All animal experiments were performed under the UK Home Office project license PEEC9E359, approved by the Animal Welfare and Ethical Review Panel of the University of Edinburgh and within the conditions of the Animals (Scientific Procedures) Act 1986.

#### 2. Embryo grafting

Approximately 70% confluent mouse CFS cells were grafted.^24,135^ The cells were topically labelled with DiI or DiO in a 100 µl droplet of M2 medium (Sigma) by expelling approximately 1 µl of labelling solution (Thermo Fisher V22886 and C7001) to the drop. After 30 sec-1 minute labelling, cells were washed 2-4x in 100 µl droplets, special care was taken for DiI to ensure all crystals were washed off. For embryos older than 6 somites, where the hindgut covers the primitive streak, the transfer pipette was introduced to the hindgut lumen by expelling a small volume of medium for better visualisation and to avoid damage to the hindgut. Cells were grafted just dorsal to the hindgut end, which approximates to the node-streak border or anterior primitive streak of these embryos.

#### 3. Immunohistochemistry

Embryo culture, cryosectioning and immunofluorescence were performed as described previously.^136^ To prevent loss of lipophilic dyes, images were acquired before immunostaining counterstaining nuclei with Hoechst 33342 (1:2000, Thermo Fisher). Pre- and post-immunostaining images of each cryosection were registered using the Hoechst as a reference channel. Imaging was performed using a Nikon Eclipse TiE widefield Inverted microscope and Plan Apo 20x/0.8 lens. Images were processed using Photoshop (Adobe) and ImageJ software (NIH).

